# TREM2 Activation by First-in-Class Direct Small Molecule Agonists: DEL Screening, Optimization, Biophysical Validation, and Functional Characterization

**DOI:** 10.1101/2025.05.22.655617

**Authors:** Hossam Nada, Farida El Gaamouch, Sungwoo Cho, Katarzyna Kuncewicz, Laura Calvo-Barreiro, Moustafa T. Gabr

## Abstract

Triggering receptor expressed on myeloid cells 2 (TREM2) is a key regulator of microglial function, and its loss-of-function variants are linked to Alzheimer’s disease (AD) and neurodegenerative disorders. While TREM2 activation is a promising therapeutic strategy, no small molecule agonists acting via direct TREM2 binding have been reported to date. Here, we describe the discovery of first-in-class, direct small molecule TREM2 agonists identified through DNA-encoded library (DEL) screening. The DEL hit (**4a**) demonstrated TREM2 binding affinity, as validated by three biophysical screening platforms (TRIC, MST, and SPR), induced Syk phosphorylation, and enhanced microglial phagocytosis. Preliminary optimization yielded **4i**, which maintained TREM2 engagement with improved selectivity over TREM1 and no cytotoxicity. Molecular dynamics simulations revealed that **4a** stabilizes a transient binding pocket on TREM2, suggesting a novel mechanism for receptor activation. These findings provide the first proof-of-concept for direct pharmacological TREM2 agonism, offering a foundation for developing therapeutics against AD and related disorders.

**Table of Contents graphic:** 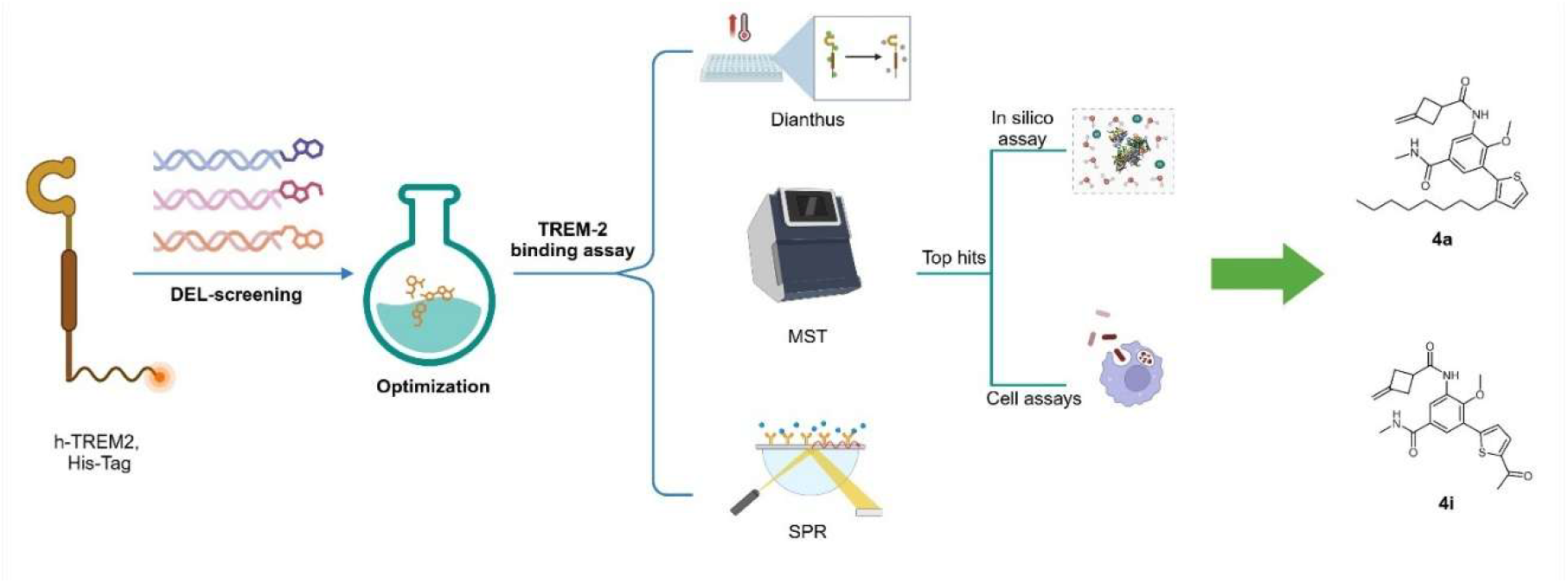

## 1. INTRODUCTION

Triggering receptor expressed on myeloid cells-2 (TREM2) is a transmembrane innate immune receptor which is primarily expressed on myeloid lineage cells such as dendritic cells, tissue-resident macrophages and microglia.^1, 2^ TREM2 is structurally composed of an extracellular V-type immunoglobulin (Ig) domain which is connected to a transmembrane helix via a short stalk region and a cytoplasmic tail devoid of intrinsic signaling motifs.^3, 4^ The transmembrane helix is responsible for the interaction with the adaptor protein DAP12 for intracellular signaling.^5^ The proteolytic cleavage or alternative splicing of TREM2 generates soluble TREM2 (sTREM2) which retains immune-modulating functions.^6^

TREM2 plays a key role in microglial function such as immune surveillance modulation, synaptic pruning, debris clearance and inflammation modulation.^7^ Under normal physiological conditions, TREM2 enhances microglial phagocytosis and clears apoptotic cells, myelin debris and misfolded protein aggregates like amyloid-beta (Aβ).^8^ TREM2 has also been found to be involved in the modulation of microglial polarization, balancing proinflammatory (M1-like) and anti-inflammatory (M2-like) states to maintain central nervous system (CNS) homeostasis.^9, 10^ Figure 1 provides a simplified overview of the TREM2 signaling pathway.

**Figure 1.**
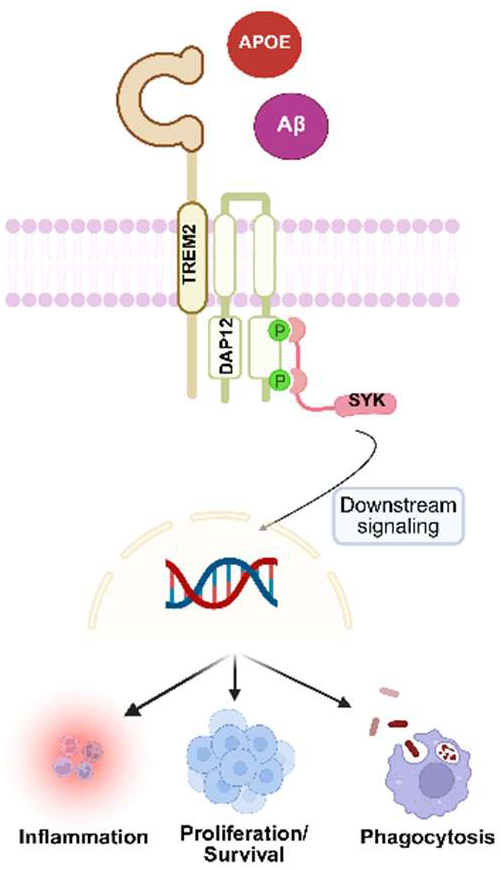
TREM2 Ligands, Signaling, and Functions. Binding of ligands to TREM2 triggers its association with DAP12 which results in the phosphorylation of DAP12 tyrosine residues by Src family kinases, initiating downstream signaling events.

Genetic evidence links TREM2 dysfunction to neurodegeneration and impaired tumor immunity, making it a compelling therapeutic target. In Alzheimer’s disease (AD), TREM2 dysfunction alters microglial activity that leads to chronic neuroinflammation, impaired debris and Aβ clearance and synaptic toxicity.^11^ Loss-of-function genetic mutations of TREM2 (e.g., R47H, R62H, T66M) have been shown to increase AD susceptibility by impairing Aβ phagocytosis, exacerbating Tau pathology and promoting sustained M1like neurotoxic signaling.^12–15^ Furthermore, studies carried out in APPswe/PS1dE9 a broadly used model for AD where mice exhibit high Aβ plaque load content at 6 months old of age, but also an exacerbated neuroinflammation, and cognitive impairments. In that study, authors demonstrated that TREM2 activity modulation could facilitate microglial phagocytosis, and reduce neuroinflammation through TREM2/DAP12 associated signaling pathway.^16^

TREM2 plays a complex role in tumors where it has been reported to act as potential immune suppressor and a modulator of anti-tumor immunity. For example, TREM2-expressing tumor-associated macrophages (TAMs) are often linked to an immunosuppressive microenvironment which limits effective anti-tumor responses. Conversely, the expression of TREM2 was found to be reduced in CNS tumors such as glioblastoma, correlating with impaired phagocytosis, accelerated tumor growth, and heightened T cell exhaustion.^17–20^ Sphingolipid-TREM2 interactions in glioma-associated macrophages drive antitumor immune responses.^17^ While elevated TREM2 expression has been associated with improved patient survival in preclinical studies where adeno-associated virus (AAV)-mediated TREM2 overexpression was showed to enhance antitumor immunity synergistically with anti-programmed cell death protein 1 (PD-1).^17^ Pharmacological TREM2 activation could enhance microglial-mediated clearance, mitigate neurotoxic inflammation, and counter glioblastoma progression.

Existing strategies to activate TREM2 signaling rely on antibodies with limited brain permeability^15^ or indirect small molecule activators that promote receptor clustering rather than direct binding^21^ (**VG-3927** or **I-192**, a TREM2 clustering agent). This gap has fundamentally limited both mechanistic studies and drug discovery efforts, as no small molecules exist that directly engage TREM2 to initiate signaling. Direct TREM2 agonists provide superior therapeutic potential compared to clustering agents by enabling precise, stoichiometric receptor engagement with defined binding kinetics, concentration-dependent signaling, and improved selectivity. Unlike clustering compounds that induce non-specific receptor aggregation, direct agonists maintain native receptor homeostasis while offering a structurally-tractable binding pocket for rational optimization. We employed DNA-encoded library (DEL) screening to discover the first small molecules that directly bind TREM2 and activate downstream signaling. The binding of these new TREM2 agonists was validated using temperature related intensity change (TRIC), microscale thermophoresis (MST), and surface plasmon resonance (SPR). Their biological activity was further assessed using cellular assays for TREM2 agonism and microglial phagocytosis. The binding mode of the TREM2 DEL hit, elucidated via computational studies, reveals a novel transient pocket, offering a template for rational optimization.

## 2. RESULTS AND DISCUSSION

### 2.1. DEL screening to identify TREM2 binders

Screening of 4.2 billion compounds from the DELopen kit (WuXi AppTec) against the human TREM2 protein led to the identification of 10,668 full-length compounds across 17 of the 27 included libraries (*data not shown*). These compounds were selected by eliminating NTC signals (C4) from the target conditions (C1, C2, and C3), applying an enrichment score threshold of 100, and focusing on compounds that bound exclusively to the TREM2 protein (A area; Figure 2A). While many full-length compounds from various libraries bound to TREM2, only certain structural patterns were enriched in a specific library. Based on these observations and the chemotypes most prevalent in the data, we identified a single top binder (Figure 2B).

**Figure 2.**
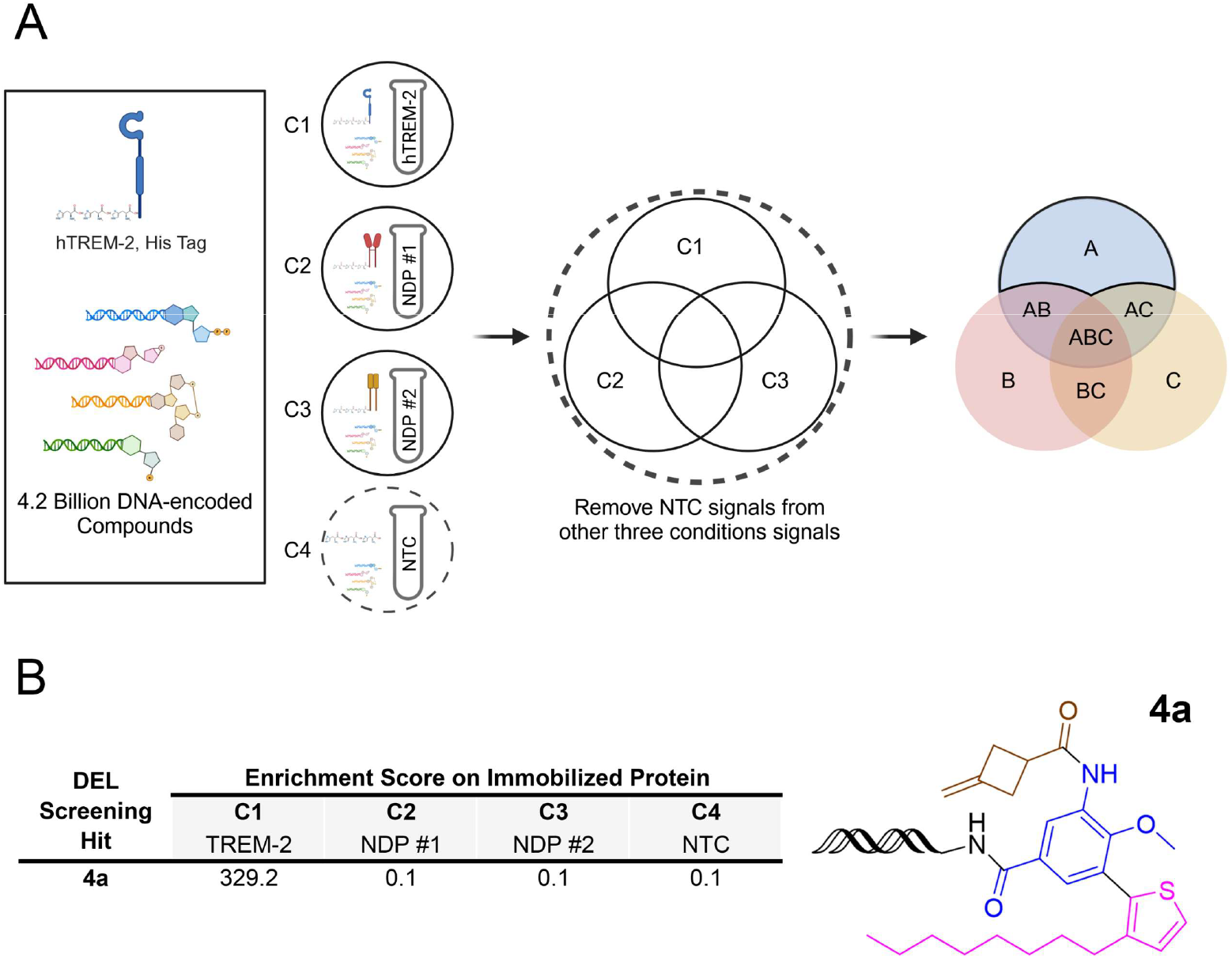
Selection of the Top Small Molecule TREM2 Binder Identified via DEL Screening. **(A)** A total of 27 DNA-encoded libraries, comprising 4.2 billion compounds, were incubated with human TREM2 protein immobilized on HisPur™ Ni-NTA Magnetic Beads (C1), two non-disclosed human proteins immobilized on the same affinity matrix (C2, C3), or the affinity matrix alone as a non-target control (NTC, C4). Following affinity selection, bound small molecules were sequenced to decode their DNA tags. After removing NTC signals (C4) and applying selection criteria, compounds exclusively detected in area A were identified as potential TREM2 binders. (**B)** Enrichment score and structure of the top small molecule binder (**4a**) selected for human TREM2 following affinity selection. Abbreviations: NDP: Non-disclosed protein.

### 2.2. Synthesis of TREM2 DEL hit and structural analogs

Based on the DEL screening results, the substituted thiophene moiety of compound **4a** was selected as the site for derivatization. This was due to the fact that unlike the other two ring moieties of compound **4a** which were shared among multiple hits, the substituted thiophene ring was unique to this particular hit indicating its amenability toward substitution. Accordingly, the DEL-discovered hit (**4a**) and its derivatives (**4b**-**l**) were synthesized according to Scheme 1. The synthetic route began with the bromination of 3-amino-4-methoxy-N-methylbenzamide (**1**) using N-bromosuccinimide (NBS) and hydrochloric acid (HCl) at room temperature for 6 hours to yield the brominated intermediate (**2**). Next, compound **2** was subjected to amide coupling with 3-methylenecyclobutane-1-carboxylic acid in the presence of 1-ethyl-3-(3-dimethylaminopropyl)carbodiimide hydrochloride (EDC·HCl) and 4-dimethylamino-pyridine (DMAP) at room temperature for two hours which afforded the intermediate compound **3**.

Finally, a Suzuki–Miyaura cross-coupling reaction was employed to introduce diverse boronic acid derivatives and generate the final compounds **4a**–**l** (see chemical structures in Table 1). In this step, compound **3** was dissolved in a 5:1 mixture of 1,2-dimethoxyethane (DME) and water and the reaction was catalyzed by tetrakis(triphenylphosphine)palladium(0) [Pd(PPh_3_)_4_] in the presence of sodium bicarbonate (NaHCO_3_) as a base at 100 °C for 6 hours.

**Table 1.**
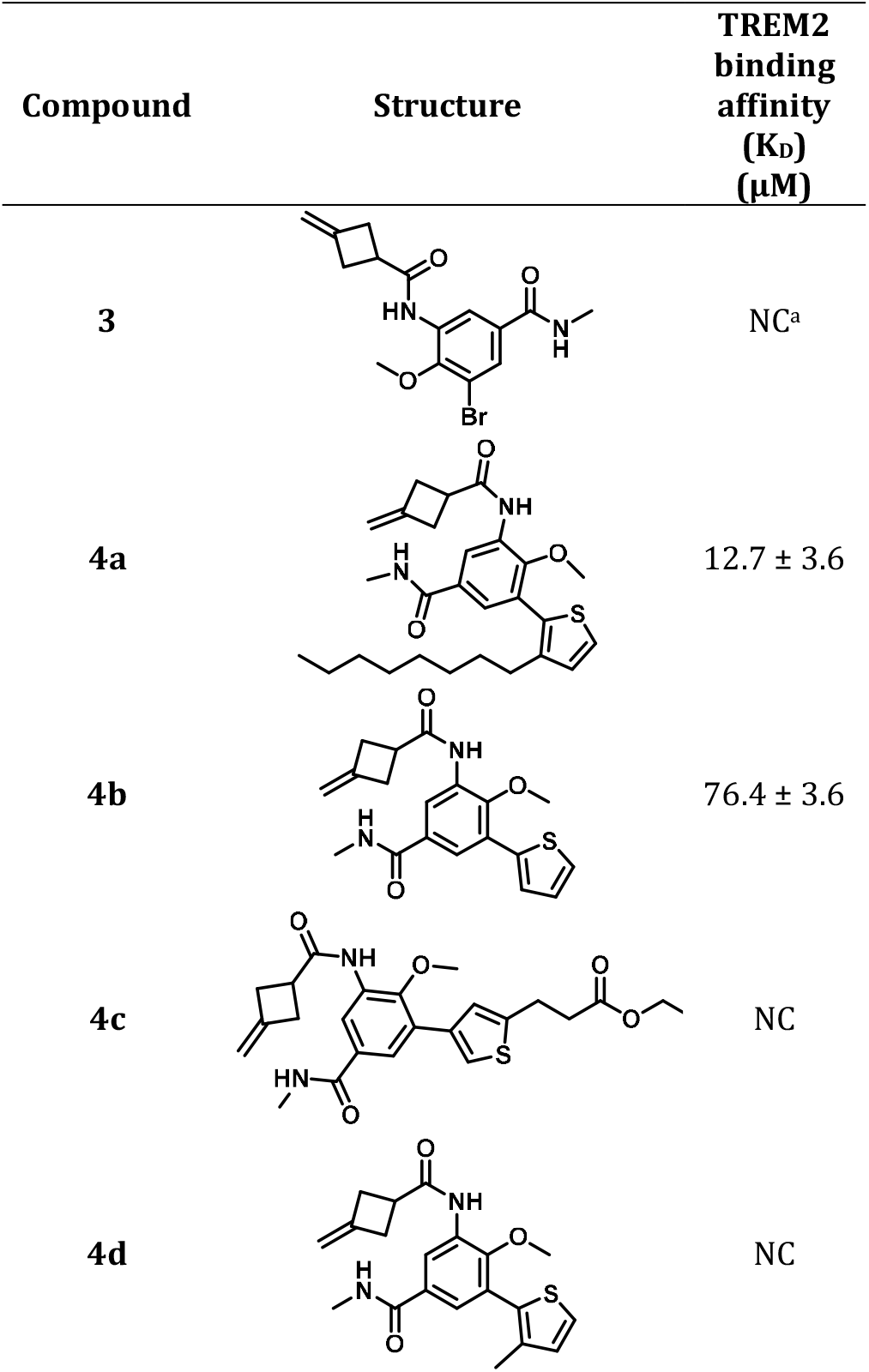
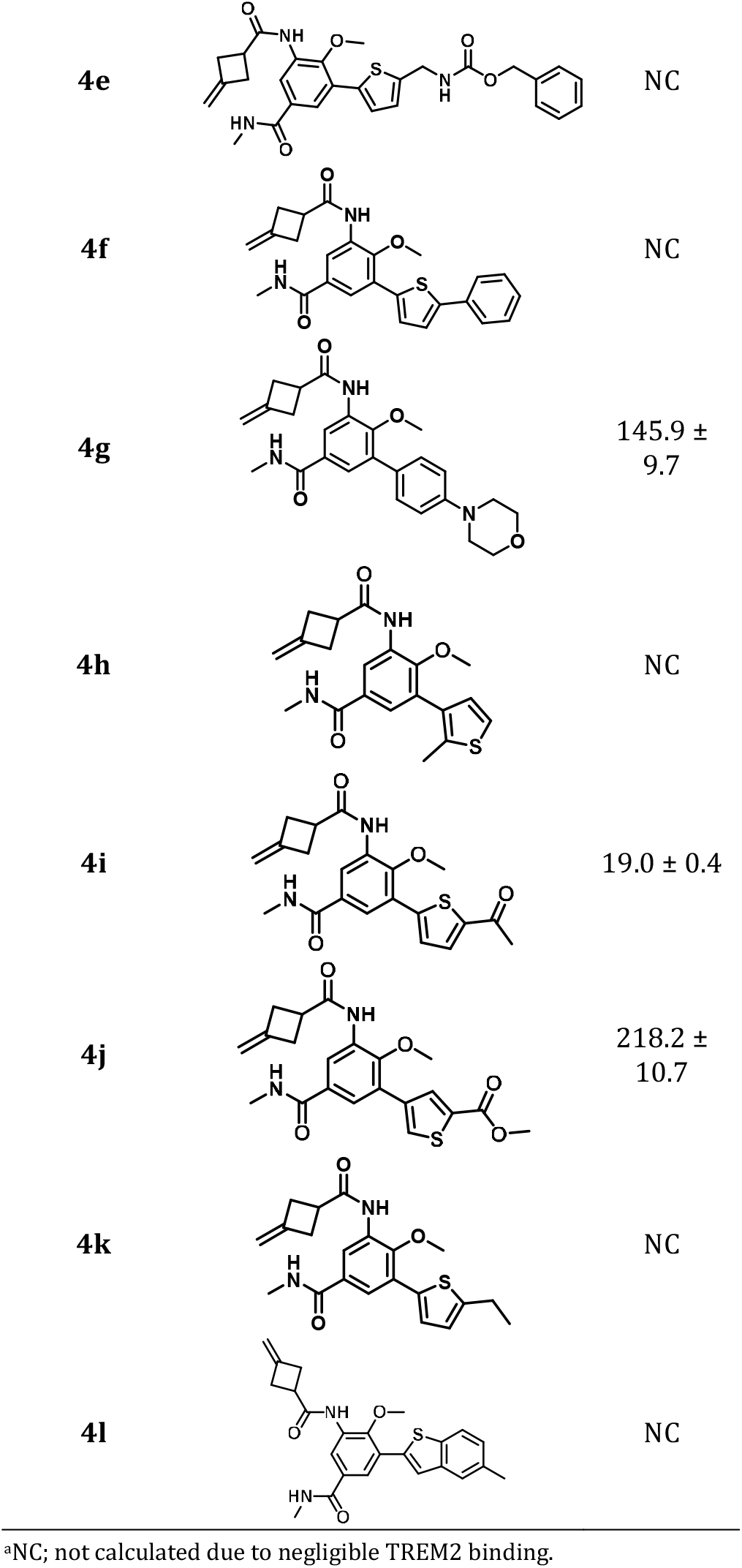
Chemical structures of the synthesized compounds and their corresponding TREM2 binding affinity.

**Scheme 1.**
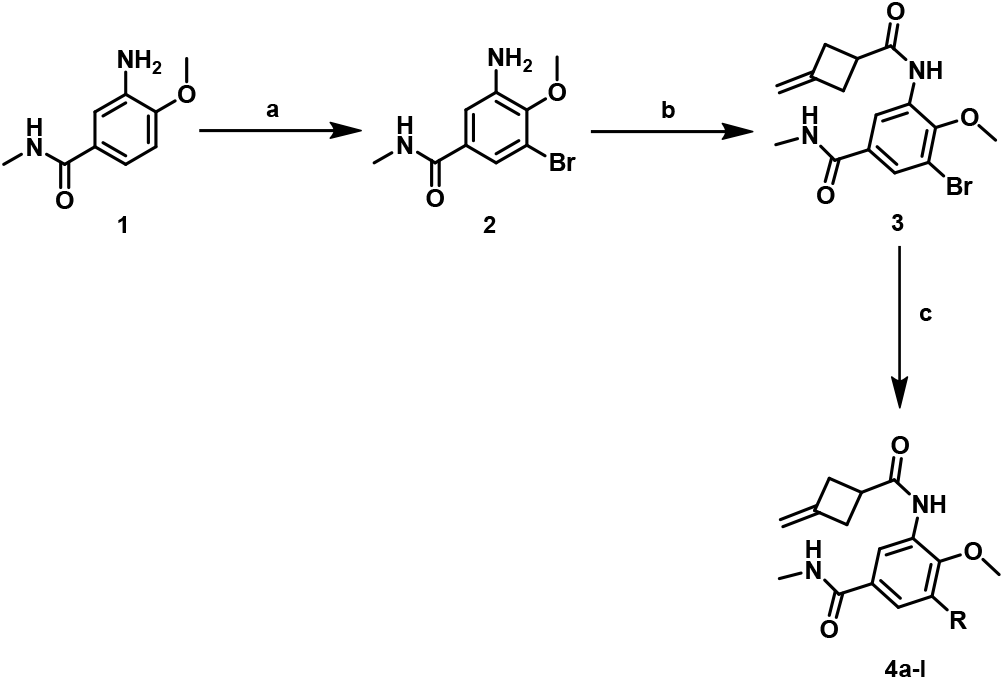
Synthesis of Compounds 4a-l. (**a**) 3-Sulfolene, NBS, HCL, rt, 6h; (**b**) 3-methylenecyclobutane-1-carboxylic acid, EDC·HCl, DMAP, rt, 2h; (**c**) Boronic acid derivatives DME/water (5:1), tertrakis, NaHCO_3_, 100 °C, 6h.

### 2.3. Biophysical evaluation of TREM2 binding affinity

#### 2.3.1. TRIC-based affinity screening

Dianthus is a technology that quantifies molecular interactions by detecting changes in fluorescence intensity that are induced by temperature variations (a principle known as Temperature-Related Intensity Change, or TRIC). Herein, the binding affinity of the synthesized compounds were evaluated using Dianthus in a single-dose assay. The 12 derivatives derived from compound **4a** were incubated with TREM2 at a concentration of 100 μM. Compounds demonstrating a signal strength change exceeding five times the standard deviation of the negative control was classified as hits. Based on the single dose assay, 7 out of 12 synthesized derivatives were deemed as potential TREM2 hits (Figure 3).

**Figure 3.**
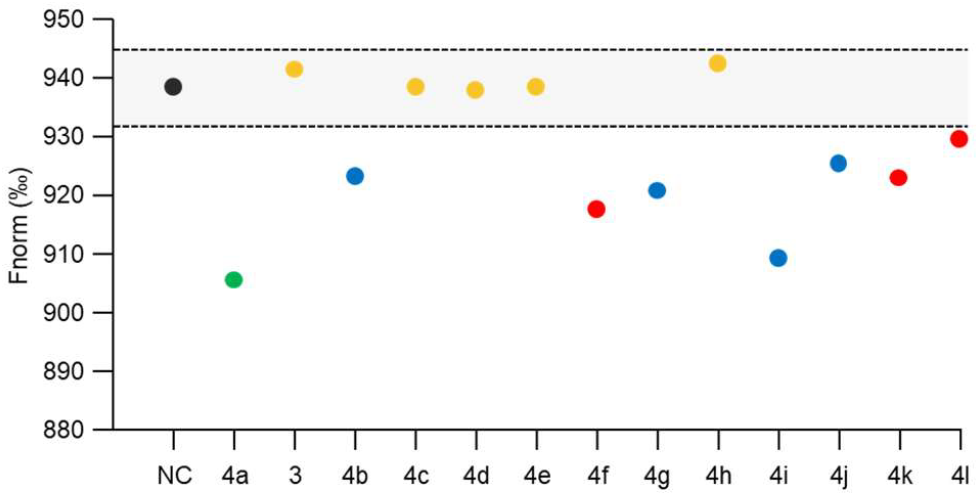
Identification and validation of TREM2-binding hits derived from compound 4a. Initial screening of 12 derivatives against TREM2 (10 nM) at a single concentration (100 μM) using Dianthus. The black dot represents negative control (TREM2 alone), and the green dot represents positive control (TREM2 with **4a**). The shaded grey area denotes the threshold for hit identification, representing signals below 5-fold standard deviation of the negative control. Compounds above this threshold were classified as potential TREM2 binders. TREM2-binding hits identified through this screening are represented by blue dots, while red dots indicate false positives resulting from aggregation.

Subsequent evaluation was conducted to confirm whether the seven identified hits exhibited dose-dependent binding to TREM2. Three out of the 7 suspected hits, compounds **4f, 4k**, and **4l**, were established to be false positives due to compound aggregation and/or their inability to exhibit dose-dependent binding (data not shown). Conversely, compounds **4b, 4g, 4i** and **4j** along with compound **4a** (Figure S1) demonstrated concentration-dependent binding to TREM2. As such these 5 compounds were considered as true binders for subsequent evaluation. To date, the only small molecule TREM2 agonist reported in the literature is **I-192** (**VG-3927**, Figure S2).^21^ However, **I-192** did not exhibit dose-dependent binding to TREM2 in our TRIC-based affinity assay (Figure S1F), suggesting that its previously reported TREM2 agonistic activity^21^ may not result from direct receptor engagement. This observation aligns with the proposed mechanism of action, wherein **I-192** is believed to promote TREM2 clustering rather than acting as a direct ligand.^21^

#### 2.3.2. MST

Based on the promising results from the Dianthus assay, MST, and SPR were used to further characterize the binding interactions of the identified hits and to confirm their binding mechanism. The results of MST assay for compounds **4a, 4b, 4g, 4i** and **4j** are illustrated in Figure 4. The DEL hit, compound **4a**, exhibited a K_D_ value of 12.7 ± 3.6 µM. Meanwhile, the synthesized derivatives, compounds **4b, 4g, 4i** and **4j** demonstrated K_D_ values of 76.4 ± 3.6, 145.9 ± 9.7, 19.0 ± 0.4 and 218.2 ± 10.7 µM, respectively (Figure 4B-D). The results indicate that compounds **4a** and **4i** exhibit the highest affinity for TREM2, thereby establishing them as promising lead compounds.

**Figure 4.**
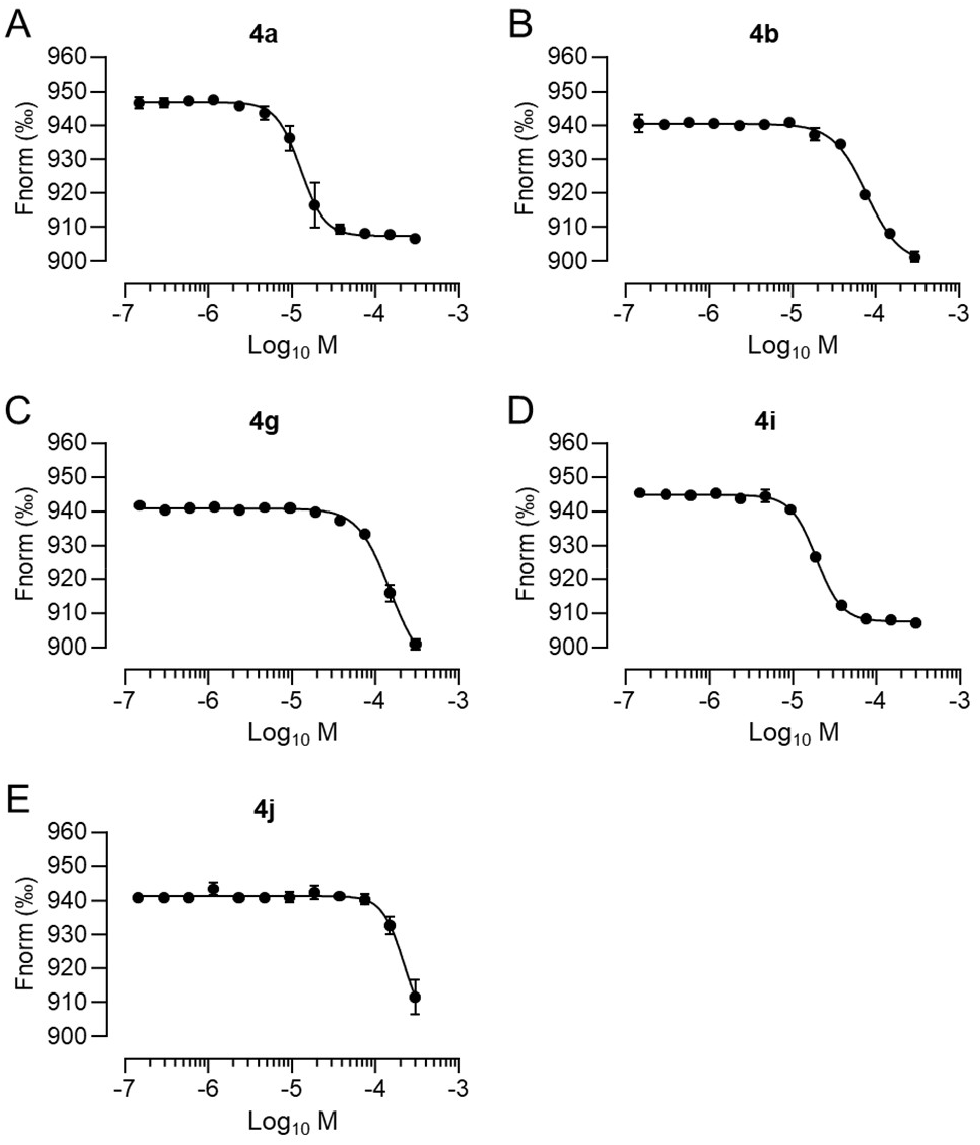
Identification and validation of TREM2 binding using MST. **(A-E)** Dose-dependent binding experiments for 5 hits (**4a, 4b, 4g, 4i**, and **4j**) using MST. Data are presented as Mean ± SEM (n = 3).

Systematic evaluation of the TREM2 binding affinity of the synthesized compounds identified distinct structural requirements for TREM2 binding (Table 1). Compound **3**, the brominated synthetic intermediate, served as the key scaffold for Suzuki-Miyaura diversification but showed negligible TREM2 binding (Table 1). This absence of activity highlights two key design features: (1) the necessity of the thiophene moiety for target engagement, as all active analogs incorporated this ring, and (2) the bromo group’s role as a synthetic handle rather than a pharmacophore. The 3-position of the thiophene ring emerged as critical-compounds with aliphatic substitutions at this position (**4a**) showed higher affinity (K_D_ = 12.7 µM) than unsubstituted (**4b**, K_D_ = 76 µM) or 2-substituted analogs (**4h**, inactive). Notably, bulky aromatic substituents on the thiophene ring, as in compounds **4f** and **4l**, resulted in a complete loss of activity, suggesting that the TREM2 binding pocket disfavors rigid, planar groups at this position. This observation implies a preference for flexible, hydrophobic substituents capable of adapting to the local binding environment and highlights the importance of conformational adaptability in ligand design for this target. The 5-position tolerated moderate substitutions, with an acetyl group (**4i**) maintaining potency (K_D_ = 19 µM) but an ester (**4j**) showing reduced affinity (K_D_ = 218 µM), likely due to steric constraints. Based on the fact that various substitutions with differing physicochemical properties retained the TREM2 binding activity, it can be surmised that the thiophene moiety is amenable to changes which paves the way for future optimization efforts.

#### 2.3.3. SPR

To validate the results obtained from the MST and Dianthus studies, SPR was employed to confirm the binding activity of the two top small molecule TREM2 binders (**4a** and **4i**). Additionally, compound **I-192** was investigated using SPR to further determine whether it functions as a direct binder. SPR provided detailed insights into the kinetics of binding, including the association rate constant (k_a_) and dissociation rate constant (k_d_). For this purpose, a single-cycle kinetics experiment was performed, immobilizing the TREM2 protein on the surface of the sensor chip and analyzing the binding of increasing concentrations of the compound as it flowed across the TREM2 surface. The resulting sensorgrams for the tested compounds are shown in Figure 5, while all kinetic constants are summarized in Table 2. The results confirm that compounds **4a** and **4i** bind to the TREM2 protein with a binding affinity of 45.9 ± 3.7 µM and 24 ± 1.5 µM, respectively. Furthermore, SPR analysis of compound **I-192** revealed no dose-dependent TREM2 binder. Taken together with the previous findings in the Dianthus assay, the results from two independent platforms consistently indicate that **I-192** does not modulate TREM2 activity via direct receptor engagement.

**Table 2.**
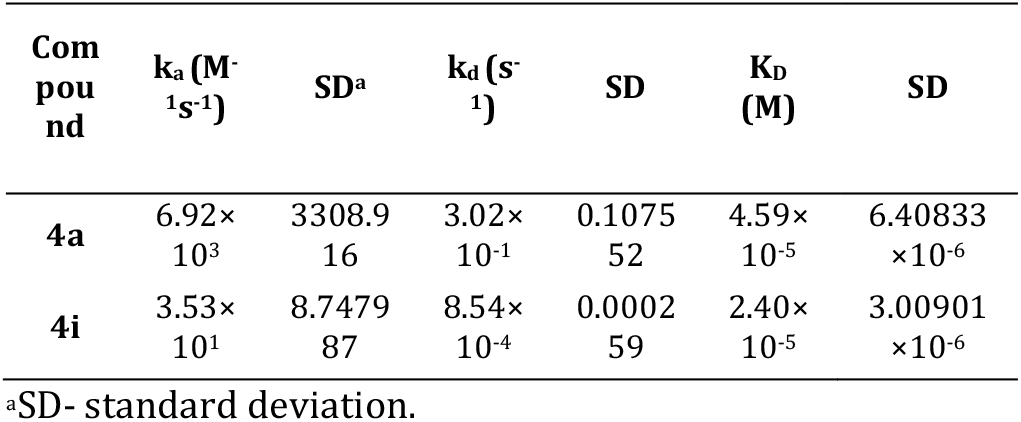
Kinetic constants calculated from SPR data for **4a** and **4i** compounds binding to TREM2 protein, immobilized on SA sensor ship. Constants were calculated with Biacore^™^ Insight Evaluation Software using data from at least three separate analyses. The 1:1 binding model was applied.

**Figure 5.**
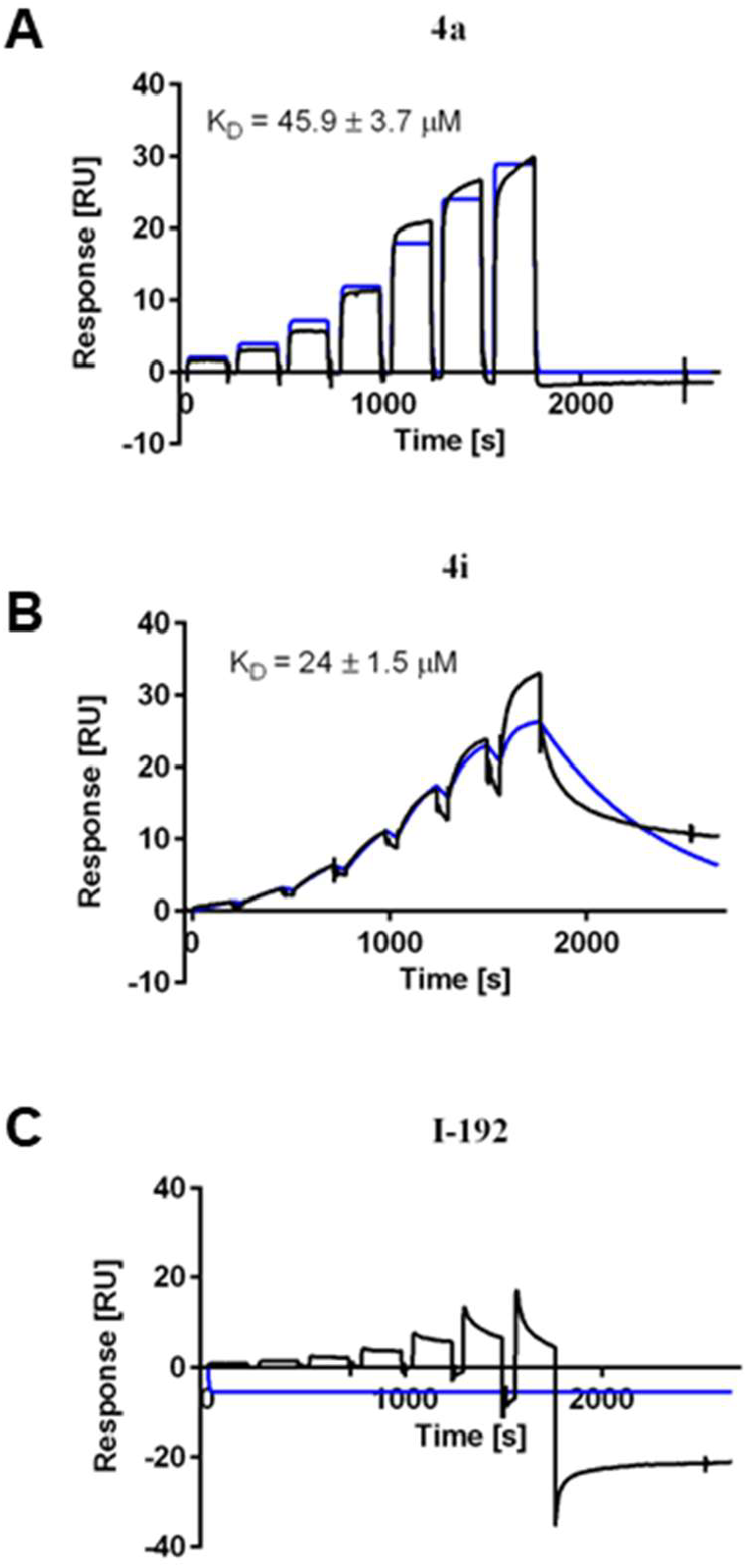
SPR analysis of TREM2 binding affinity for 4a, 4i, and I-192. Single-cycle kinetic sensorgrams of **4a (A), 4i (B)**, and **I-192 (C)** compounds interacting with the TREM2 protein using SPR. Black lines represent the experimental data, while blue lines show the 1:1 kinetic binding model fit. K_D_ values are presented as Mean ± SEM (n = 3).

In addition to evaluating the binding affinity of compounds **4a** and **4i** toward TREM2, the SPR analysis provided detailed insights into their binding kinetics (Table 2). Compound **4a** exhibited an association rate constant (k_a_) of 6.92 × 10^3^ M^−1^s^−1^ and a dissociation rate constant (k_d_) of 3.02 × 10^−1^ s^−1^. In contrast, compound **4i** displayed a markedly slower association rate (k_a_) of 3.53 × 10^1^ M^−1^s^−1^ but a significantly lower dissociation rate (k_d_) of 8.54 × 10^−4^ s^−1^. The lower K_D_ value of compound **4i** compared to **4a** indicates its superior overall binding affinity for TREM2, which is primarily attributed to its substantially slower dissociation rate despite its reduced association rate. These findings suggest that compound **4i** forms a more stable complex with TREM2, highlighting its potential as a promising candidate for further investigation.

#### 2.3.4. TREM2 selectivity and cytotoxicity comparative assays

TREM1 and TREM2 have distinct and often conflicting roles, highlighting the need for establishing the selectivity of any TREM modulator. Accordingly, the two most promising TREM2 binders, compounds **4a** and **4i**, were selected for a comparative binding selectivity assessment against TREM2 and TREM1 using MST (Figure 6). The results of the comparative binding assay showed that **4a** exhibited a K_D_ value of 18.4 ± 1.7 µM for TREM1 which indicates a slight selectivity of around 1.5-fold between TREM1 and TREM2 (TREM2 K_D_ = 12.7 ± 3.6 µM). In contrast, **4i** demonstrated a K_D_ value of 39.8 ± 2.4 µM for TREM1, revealing approximately 2-fold selectivity for TREM2 (TREM2 K_D_ = 19.0 ± 0.4 µM) over TREM1.

**Figure 6.**
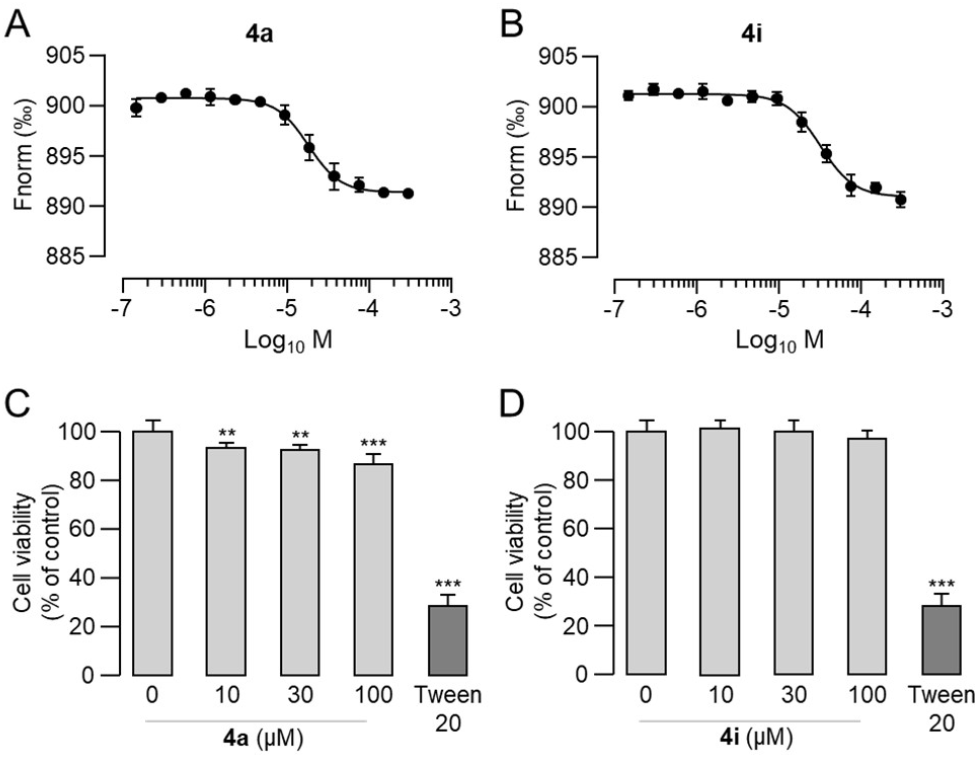
Assessment of TREM2 selectivity and cytotoxicity. (A-B). Dose-dependent binding experiments for **4a** and **4i** in TREM1. Mean ± SEM (n = 3). **(C-D)** Effect of **4a** and **4i** on cell viability in HMC3 cells. Cells were serum-deprived for four hours prior to compound treatment, then treated with compounds for one hour before cell viability was determined by MTS colorimetric assay. Mean ± SEM (n = 5). \*\**p* < 0.01, \*\*\**p* < 0.001 analyzed by Student’s unpaired twotailed t test.

Next, the impact of compounds **4a** and **4i** on cell viability was assessed by3-(4,5-Dimethylthiazol-2-yl)-5-(3-carboxymethoxyphenyl)-2-(4-sulfophenyl)-2H-tetrazolium (MTS) colorimetric cell viability assay in a human microglial cell line that endogenously express TREM2. Briefly, cells were exposed to 10, 30 and 100 μM of each compound. **4a** exhibits a reduction of cell viability by 7% at the lowest concentration (10 μM) while **4i** showed no changes in cell viability at any concentration, making it a potential therapeutic lead.

### 2.4. Computational assay

Biophysical mapping studies of TREM2 have revealed the existence of three distinct binding sites on TREM2 interface: hydrophobic site, basic site and site 2. The hydrophobic surface of TREM2 allows the binding of TREM2 with Apolipoprotein E (APOE). Meanwhile, the basic site was revealed to be responsible for the interaction with charged residues such as IL_34_.^22,23^ Based on these reports, the synthesized compounds were docked into the three binding sites (data not shown). The docking results did not reveal a correlation between the docking scores of the compounds and the three reported binding sites of TREM2.

To investigate the binding dynamics of compound **4a** with TREM2, an unrestricted 1000 ns (1 µs) molecular dynamics (MD) simulation was performed using the DESMOND software (Figure 7A-B, Video S1). The simulation revealed that after an initial 30 ns, the protein underwent a conformational change which led to the formation of a small pocket and potentially representing a transient protein state (Figure 7B). Compound **4a** bound to this transient pocket, stabilizing it and remaining associated throughout the entire molecular dynamic (MD) simulation (Figure 7C, Video S2). The protein-ligand complex exhibited an average root-mean-square deviation (RMSD) of approximately 1.7 Å which further supported the stability of the ligand within this newly identified transient pocket site (TPS).

**Figure 7.**
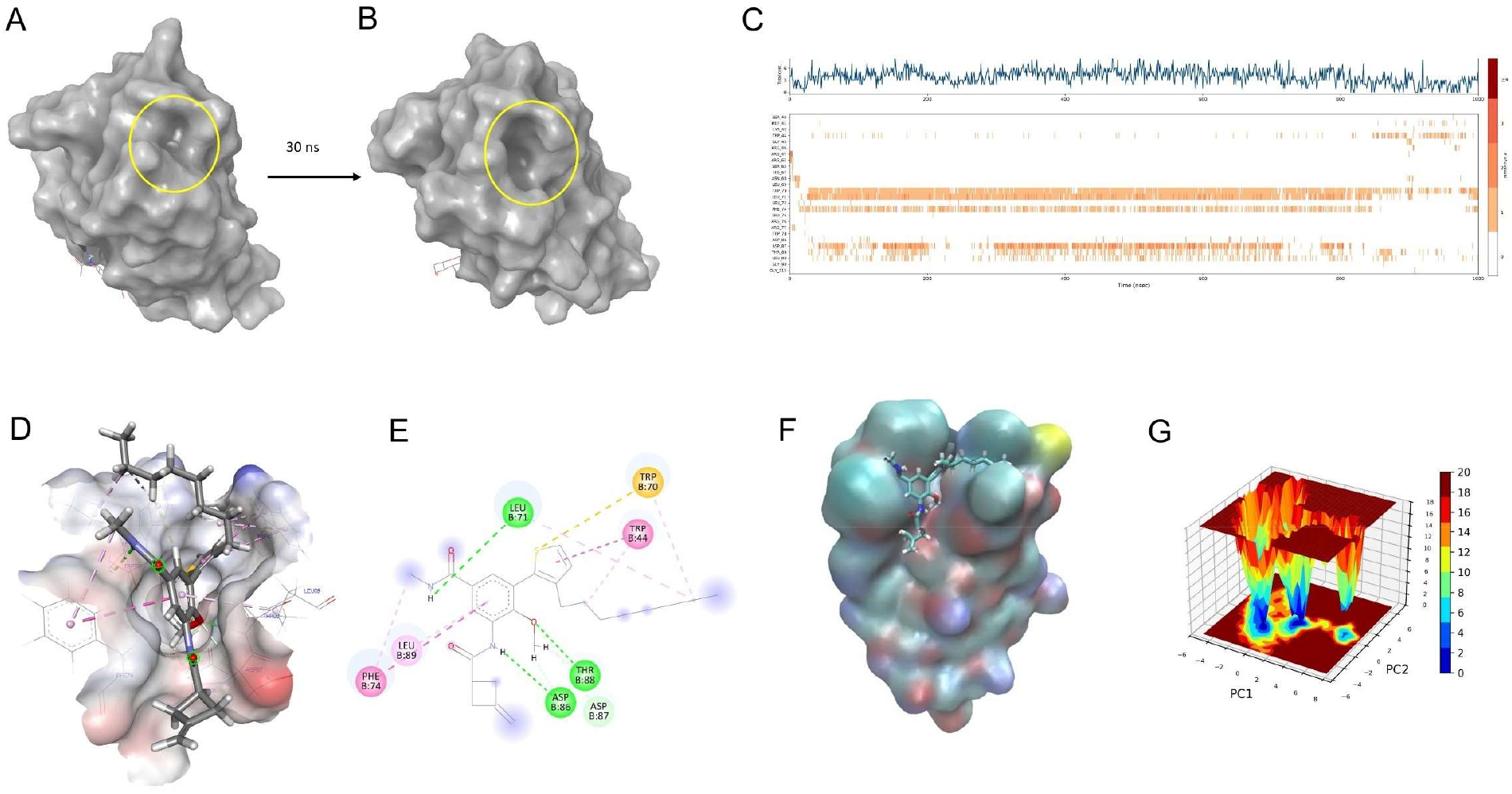
In silico investigation of the binding mechanism of compound 4a with TREM2. **(A)** Surface representation of TREM2 in its native state (PDB ID: 5UD7). **(B)** Formation of a small pocket, potentially representing a transient protein state. **(C)** TREM2/**4a** contact map throughout the 1 ms MD simulation. **(D)** 3D docking representation of compound **4a** within the transient pocket. **(E)** 2D interaction diagram illustrating key binding interactions of **4a** within the transient pocket. **(F)** Snapshot of the hydrophobic groove formed due to the long aliphatic “tail” of **4a. (G)** Gibbs free energy landscape depicting the conformational stability of the TREM2/**4a** complex.

Next, a docking grid was generated employing the stabilized site and compound **4a** was redocked within the TPS to investigate its binding mechanism. The resulting 2D interaction diagram (Figure 7E) illustrated binding interactions between compound **4a** and its target protein. **4a** established multiple interactions with surrounding amino acid residues which may explain the observed stability within the binding pocket. Among the established interactions, hydrogen bonds were observed with ASP 86, ASP 87, THR 88, and LEU B:71 amino acid residues as well as hydrophobic interactions and a π-sulfur interaction with TRP70. The clustering of these interactions in distinct regions suggests an optimized binding mode where both hydrophobic and polar forces play crucial roles in maintaining ligand stability within the transient pocket.

To ensure the robustness and validity of the results, a second 250 ns molecular dynamics (MD) simulation was performed using GROMACS, providing an independent verification of the findings obtained from the primary MD simulation. The TREM2/**4a** complex exhibited an average RMSD of 0.9 Å, indicating high structural stability, and maintained at least two hydrogen bonds throughout the entire simulation, with an average bond distance of 2.85 Å. Notably, the interaction of compound **4a** with TREM2 revealed that the long aliphatic chain of the “tail” moiety induces a conformational change in the stabilized transient site and leading to the formation of a hydrophobic groove. This conformational change enhances the stability of the compound within the binding site (Figure 7F, Video S3). Gibbs free energy analysis (Figure 7G) identified multiple energy minima, with the deepest basins corresponding to the observed conformational change which further supporting its thermodynamic favorability. Collectively, these findings suggest a potential mechanistic basis for the high binding affinity of compound **4a** toward TREM2.

### 2.5. Cellular assay

#### 2.5.1. Measurement of TREM-2 induced activation using intracellular Syk phosphorylation

The two hits from the screening assays, compounds **4a** and **4i**, were selected to assess their ability to activate TREM2 *in vitro*. Compounds **4a** and **4i** were evaluated in a human immortalized cell line (Human Embryonic Kidney) HEK 293 over expressing the human TREM2 receptor and its associated transmembrane adaptor protein DAP12 (HEK293-hTREM2/DAP12). The selective activation of TREM2/DAP12 will lead to the recruitment and phosphorylation of the spleen tyrosine kinase (SYK) which will then promote the activation of multiple downstream mediators and mechanisms that includes microglial phagocytosis, cell proliferation and reduction of inflammation.

The commercial AlphaLISA detection kit was used to assess SYK activation by measuring Tyrosine 525/526 phosphorylation levels following treatment with compounds **4a** and **4i**. The SPARK plate reader (TECAN) with the AlphaLISA technology settings was employed to measure the fluorescence intensities for each condition. Syk phosphorylation changes were quantified and compared to control condition (Figure 8A). Compound **4a** significantly increases Syk phosphorylation: phospho-Syk level is approximately 2.3-fold higher following treatment with compound **4a** compared to control condition (\**p*<0.05). Similarly, compound **4i** induced an increase in phospho-SYK levels, though to a lesser extent than compound **4a**.

**Figure 8.**
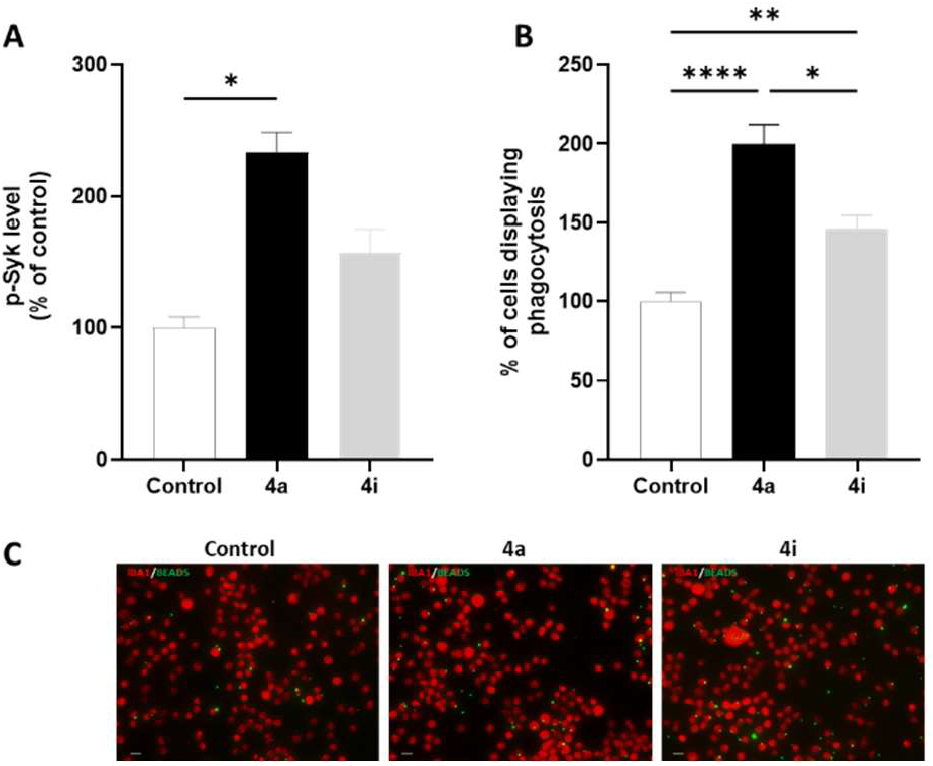
In vitro validation of the TREM2 binding activity. **(A)** Histogram representing the quantification of phospho-Syk levels, measured using AlphaLisa technique, in untreated condition or after treatment with compound **4a** and **4i** (25 uM-1 h). Phospho-Syk changes in treated conditions are expressed as percentage of control (N=3). (**B)** Histogram representing the quantification of BV2 cells showing at least one fluorescent latex bead in their cell body (percentage of control) after treatment with compound **4a** and **4i** (25 μM-1 h) (n=36-40 pictures/group (N=5)). Statistical analyses were performed using One-Way ANOVA followed by a Kruskal-Wallis multiple comparisons test was done. Error bars represent means ± SEM. \**p* < 0.05, \*\**p* < 0.005, \*\*\*\**p* < 0.0001. (**C)** Representative images of BV2 cells immuno-labelled with antibody directed against IBA1 (red) and containing fluorescent beads (green) in their cell body (scale bar=20 μM), in control condition or following treatment with compound **4a** or **4i**.

#### 2.5.2. Assessment of selected compound’s role in glial phagocytosis

Glial phagocytosis is a protective mechanism associated with several neurodegenerative diseases: glial cells, the immune cells of the brain will act as cleaners thanks to their ability to sense their environment. Here, we examined whether compounds **4a** and **4i** could play a role in the phagocytic activity of microglia cells and therefore be potential actors in neurodegenerative disease modulation. Phagocytic function was measured in BV2 cells, a murine microglia cell line, using fluorescent latex beads. BV2 cells were serum deprived prior treatment with either compound **4a** or **4i**, and beads addition. Following treatment with compound **4a**, we observed twice as many cells containing 1 or more beads compared to the control condition. Compound **4i** also increased beads phagocytosis by approximately 46% in comparison with control, however its effect on glial phagocytosis remains significantly lower than what is observed for compound **4a** (Figure 8B).

## 3. Conclusion

This work describes the discovery and characterization of the first direct small molecule agonists of TREM2, establishing a new pharmacological approach for TREM2 agonism. Through DEL screening of 4.2 billion compounds followed by rigorous triage using three orthogonal biophysical methods (TRIC, MST, and SPR), we identified hit compound **4a** as a TREM2 binder. Structure-activity relationship (SAR) studies yielded optimized analog **4i**, which maintained TREM2 engagement while exhibiting improved selectivity over TREM1 and reduced cytotoxicity. Our study overcomes a key limitation in small molecule-based TREM2 activation, where previously reported small molecule TREM2 agonists exhibited indirect agonism via receptor clustering. By contrast, our compounds display mechanistically defined, receptor-selective agonism, enabling clearer pharmacological interrogation and enhanced translational potential.

The comprehensive validation cascade developed here-employing TRIC for initial hit confirmation, MST for affinity determination in solution, and SPR for full kinetic characterization-provides a robust workflow (Figure 9) to validate TREM2 binders from promiscuous or aggregating compounds. We anticipate that the biophysical screening workflow established in this study will serve as a valuable framework for future TREM2-targeted drug discovery efforts.

**Figure 9.**
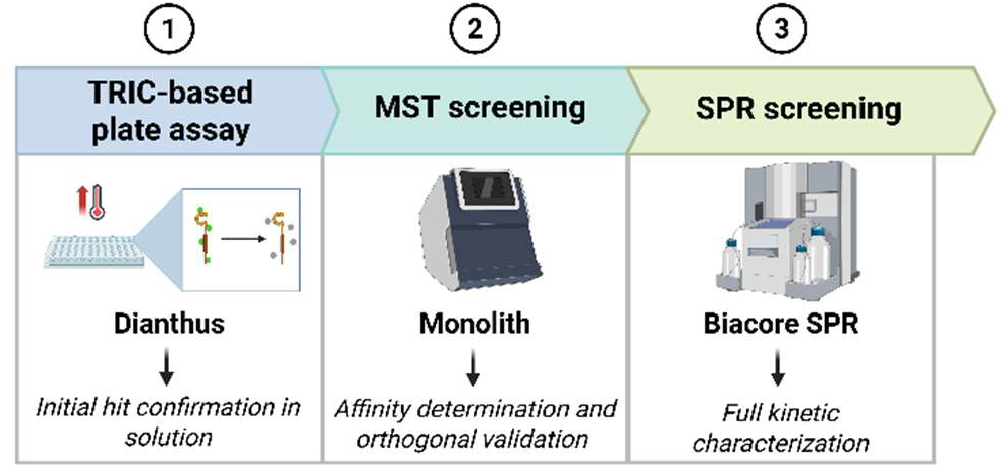
Established workflow for biophysical validation of potential small molecule TREM2 binders. Following a screening campaign to identify small molecule TREM2 binders, a TRIC-based assay using Dianthus as an initial step for hit confirmation. Subsequently, Monolith enables orthogonal validation and affinity determination in solution. Finally, full kinetic characterization of the true hits can be achieved using SPR.

Functional characterization in cellular models confirmed that both compounds (**4a** and **4i**) act as TREM2 agonists, inducing SYK phosphorylation and significantly enhancing microglial phagocytic activity. MD simulations revealed these compounds stabilize a novel transient binding pocket through specific interactions with Asp86, Thr88 and Leu71 residues in TREM2. This work advances the field by moving beyond indirect clustering strategies to achieve small molecule mediated activation of TREM2 a critical step toward developing targeted therapies for neurodegenerative diseases and other conditions involving TREM2 dysfunction. The compounds and characterization methods described herein provide a foundation for future medicinal chemistry optimization and preclinical development of TREM2-targeted therapeutics.

## 4. EXPERIMENTAL

### 4.1. DEL screening

A DEL containing 4.2 billion compounds, provided by WuXi AppTec, was screened against the human triggering receptor expressed on myeloid cells 2 (TREM2) protein. Prior to affinity selection, a protein capture experiment was conducted to ensure stable immobilization of the TREM2 protein (Cat. # 11084-H08H, Sino Biological, Beijing, China) onto the affinity matrix. Based on this preliminary assessment, the selection buffer was optimized to 1x PBS, 0.05% Tween-20, 0.1 mg/mL sssDNA, and 10 mM imidazole, while the elution buffer comprised 1x PBS and 0.05% Tween-20. Each selection round utilized 5 µg of TREM2 protein and HisPur™ Ni-NTA Magnetic Beads (Cat. # 88831, Thermo Fisher, Waltham, MA, USA) as the affinity matrix.

Affinity selection, including protein immobilization and subsequent selection rounds, was performed following the manufacturer’s protocol. The DELopen kit provided four identical copies of 27 distinct libraries, collectively encompassing 4.2 billion compounds. One copy was used for TREM2 screening, two other copies for separate human protein targets, and the fourth as a non-target control (NTC) using HisPur™ Ni-NTA Magnetic Beads without protein to evaluate nonspecific binding.

The heated eluted sample obtained after the second round of selection was designated as the post-selection sample based on the manufacturer’s self-QC step. Briefly, this real-time, semi-quantitative procedure ensures that the amount of selected DEL molecules in the sample is less than 5×10^8^ copies (standard control) and that no cross-contamination is present. This sample was sent to WuXi AppTec, which confirmed that it met sequencing quality standards. Following this verification, the sample proceeded to next-generation sequencing (NGS) to decode the DNA tags associated with each individual small molecule.

Data analysis identified a top-binding compound, whose structure was disclosed to the authors by WuXi AppTec. Upon request, the tag-free version of the compound was synthesized for further study.

### 4.2 Chemistry

Commercially available reagents and solvents were employed in the experiments without prior purification. All intermediates were verified using ^1^H NMR and mass spectrometry (MS), while the final products were characterized by ^1^H and ^13^C NMR, as well as high-resolution mass spectrometry (HRMS). ^1^H and ^13^C NMR spectra were acquired on a Bruker Avance III 500HD spectrometer (500 or 125 MHz, Billerica, MA, USA) using DMSO-*d*6 (Sigma Aldrich) as a solvent at room temperature (r.t.). Chemical shifts are reported in parts per million (ppm) relative to tetramethylsilane (*δ* 0.00, s), with coupling constants given in hertz (Hz). Signal multiplicities are denoted as s (singlet), d (doublet), t (triplet), q (quartet), and m (multiplet). MS spectra were obtained using an SQ Detector 2 mass spectrometer (Waters, USA), while HRMS spectra were recorded on a Waters LCT Premier XE mass spectrometer (USA).

#### 4.2.1. General Procedure for the Synthesis of Compound 2 (3-amino-5-bromo-4-methoxy-*N*-methylbenzamide)

3-amino-4-methoxy-*N*-methylbenzamide (1 eqv.) was dissolved in DMF (20 mL) in a 25 mL oven-dried round-bottom flask equipped with a stirring bar. N-Bromosuccinimide (NBS, 1 eqv.) was slowly added to the solution, followed by the addition of butadiene sulfone (0.7 eqv.) and aqueous HCl (prepared by diluting 2 mL of concentrated HCl with 12.5 mL of water) at room temperature. The reaction mixture was stirred for 6 hours, and the progress of the reaction was monitored by thin-layer chromatography (TLC). Upon completion, the mixture was poured into water and extracted with ethyl acetate. The combined organic layers were concentrated under reduced pressure, and the resulting residue was purified by column chromatography on silica gel using a hexane-ethyl acetate gradient to yield compound 2.

^1^H NMR (500 MHz, DMSO) *δ* 8.03 (d, J = 4.7 Hz, 1H), 7.97 (s, 1H), 6.95 (s, 1H), 6.67 (s, 1H), 5.02 (s, 2H), 3.80 (s, 3H), 2.91 (s, 3H). MS: m/z calcd for [M-H]^2+^ 260.0, found 260.23.

#### 4.2.2. General Procedure for the Synthesis of Compound 3

3-methylenecyclobutane-1-carboxylic acid (1 eqv.) was dissolved in DMF and cooled down to 0–5 °C using an ice bath. Next, EDC.HCL was added to the reaction mixture and stirred for few minutes at 0 °C to allow the activation of the carboxylic acid, forming an O-acylisourea intermediate. 2 eqv. Of DMAP were then slowly added to the reaction mixture which was slowly followed by the addition of 3-amino-5-bromo-4-methoxy-*N*-methylbenzamide (1 eqv.) to yield compound 3 (3-bromo-4-methoxy-N-methyl-5-(3-methylenecyclobutane-1-carboxamido)benzamide). ^1^H NMR (500 MHz, DMSO) *δ* 8.03 (d, *J* = 4.8 Hz, 1H), 7.97 (s, 1H), 6.94 (s, 1H), 6.67 (s, 1H), 5.02 (s, 2H), 3.80 (s, 3H), 2.91 (s, 4H), 2.71 (d, *J* = 4.7 Hz, 3H). ^13^C NMR (126 MHz, DMSO) *δ* 173.54, 167.90, 162.79, 150.73, 145.49, 131.49, 127.13, 121.86, 115.73, 113.60, 106.83, 56.88, 35.49, 31.26, 26.49. MS: m/z calcd for [M-H]^2+^ 352.04, found 355.16. HRMS: m/z calcd for [M+Na]^+^ 377.0300, found 377.0305 (^81^Br).

#### 4.2.3. General Procedure for the Synthesis of Compound 4a-l

In a sealed tube, Compound **3** (1 equiv) was dissolved in DME and degassed water (5:1). Tetrakis (0.15 equiv), the appropriate boronic acid derivative (2 equiv), and Na_2_CO_3_ were then added and the resulting reaction mixture was degassed with nitrogen for 15 minutes. The reaction vessel was sealed and heated at 80 °C overnight. Upon completion, the aqueous layer was extracted with CH_2_Cl_2_, and the organic layer was dried over anhydrous MgSO_4_. The solvents were removed under reduced pressure, and the crude product was purified by flash chromatography on silica gel using a pentane/diethyl ether (9:1) gradient to afford the desired product.

##### 4.2.3.1. Compound 4a (4-methoxy-*N*-methyl-3-(3-meth-ylenecyclobutane-1-carboxamido)-5-(3-octylthiophen-2yl)benzamide)

Sticky orange solid (67mg, 50% yield). ^1^H NMR (500 MHz, DMSO) *δ* 9.27 (s, 1H), 8.23 (s, 1H), 7.74 (d, *J* = 4.7 Hz, 1H), 7.42 (d, *J* = 5.2 Hz, 1H), 6.97 (d, *J* = 5.1 Hz, 1H), 6.88 (s, 1H), 4.81 (t, *J* = 2.5 Hz, 2H), 3.93 (s, 2H), 3.47 (q, *J* = 8.1 Hz, 1H), 2.98 – 2.78 (m, 4H), 2.58 (d, *J* = 4.6 Hz, 3H), 2.43 – 2.35 (m, 2H), 1.48 (d, *J* = 7.7 Hz, 2H), 1.35 – 1.12 (m, 12H). ^13^C NMR (126 MHz, DMSO) *δ* 173.49, 168.83, 149.67, 145.56, 139.78, 135.63, 131.21, 128.76, 127.23, 124.89, 121.30, 114.45, 106.83, 73.99, 56.48, 35.52, 34.44, 31.74, 30.46, 29.38, 29.26, 29.13, 28.52, 26.52, 25.44, 22.54, 14.41. HRMS: m/z calcd for [M+Na]^+^ 491.2345, found 491.2324.

##### 4.2.3.2. Compound 4b (4-methoxy-*N*-methyl-3-(3-meth-ylenecyclobutane-1-carboxamido)-5-(thiophen-2-yl)benzamide)

white powder (30mg, 43% yield). ^1^H NMR (500 MHz, DMSO) *δ* 9.26 (s, 1H), 8.21 – 7.98 (m, 2H), 7.55 (dd, *J* = 5.1, 1.2 Hz, 1H), 7.22 (dd, *J* = 3.6, 1.2 Hz, 1H), 7.15 – 7.04 (m, 2H), 4.80 (t, *J* = 2.5 Hz, 2H), 3.93 (s, 2H), 2.96 – 2.78 (m, 4H), 2.66 (d, *J* = 4.6 Hz, 3H). ^13^C NMR (126 MHz, DMSO) *δ* 173.50, 169.73, 149.79, 145.56, 141.88, 129.75, 127.96, 127.55, 127.10, 126.69, 126.42, 121.10, 112.17, 106.82, 56.50, 35.53, 34.44, 26.52. HRMS: m/z calcd for [M+Na]^+^ 379.1093, found 379.1109.

##### 4.2.3.3. Compound 4c (ethyl 3-(4-(2-methoxy-5-(methylcarbamoyl)-3-(3-methylenecyclobutane-1-carboxamido)phenyl)thiophen-2-yl)propanoate)

Pink powder (15mg, 19% yield). ^1^H NMR (500 MHz, DMSO) *δ* 9.20 (s, 1H), 8.07 (s, 1H), 7.93 (d, *J* = 4.7 Hz, 1H), 7.34 (d, *J* = 1.4 Hz, 1H), 7.06 (s, 1H), 6.99 (d, *J* = 1.5 Hz, 1H), 4.80 (t, *J* = 2.4 Hz, 2H), 4.10 (q, *J* = 7.1 Hz, 2H), 3.92 (s, 3H), 3.07 (t, *J* = 7.2 Hz, 2H), 2.96 – 2.78 (m, 4H), 2.68 (t, *J* = 7.3 Hz, 2H), 2.64 (d, *J* = 4.6 Hz, 2H), 1.21 (t, *J* = 7.1 Hz, 3H). ^13^C NMR (126 MHz, DMSO) *δ* 173.38, 172.23, 170.16, 149.91, 145.60, 142.93, 140.39, 131.93, 129.41, 126.28, 120.93, 111.93, 106.79, 60.51, 56.43, 35.78, 35.53, 34.43, 26.50, 25.18, 14.57. HRMS: m/z calcd for [M+Na]^+^ 479.1617, found 479.1606.

##### 4.2.3.4. Compound 4d (4-methoxy-*N*-methyl-3-(3-meth-ylenecyclobutane-1-carboxamido)-5-(3-methylthiophen-2-yl)benzamide)

^1^H NMR (500 MHz, DMSO) *δ* 9.27 (s, 1H), 8.21 (s, 1H), 7.76 (d, *J* = 4.8 Hz, 1H), 7.42 (dd, *J* = 5.1, 1.2 Hz, 1H), 6.94 – 6.89 (m, 2H), 4.83 – 4.78 (m, 2H), 3.89 (d, *J* = 1.3 Hz, 3H), 3.48 (p, *J* = 8.0 Hz, 1H), 2.97 – 2.79 (m, 4H), 2.60 (dd, *J* = 4.6, 1.2 Hz, 3H), 1.30 – 1.16 (m, 1H). ^13^C NMR (126 MHz, DMSO) *δ* 173.50, 169.06, 149.78, 145.57, 135.80, 134.63, 133.65, 132.53, 131.12, 130.32, 127.18, 124.71, 121.40, 114.25, 106.83, 56.54, 35.53, 31.17, 26.61, 14.75. HRMS: m/z calcd for [M+Na]^+^ 393.1249, found 393.1254.

##### 4.2.3.5. Compound 4e (benzyl ((5-(2-methoxy-5-(methylcarbamoyl)-3-(3-methylenecyclobutane-1-carboxamido)phenyl)thiophen-2-yl)methyl)carbamate)

^1^H NMR (500 MHz, DMSO) *δ* 9.25 (s, 1H), 8.09 (d, *J* = 4.8 Hz, 2H), 7.95 (t, *J* = 6.2 Hz, 1H), 7.41 – 7.31 (m, 5H), 7.04 – 7.02 (m, 2H), 6.90 (d, *J* = 3.6 Hz, 1H), 5.08 (s, 2H), 4.80 (s, 2H), 4.37 (d, *J* = 6.1 Hz, 2H), 3.92 (s, 3H), 2.96 – 2.88 (m, 2H), 2.85 – 2.82 (m, 1H), 2.65 (d, *J* = 4.5 Hz, 3H). ^13^C NMR (126 MHz, DMSO) *δ* 173.49, 169.73, 156.64, 149.78, 145.56, 143.57, 140.83, 137.55, 133.64, 132.83, 132.53, 132.51, 132.01, 129.57, 129.20, 128.85, 128.25, 127.63, 127.01, 125.86,121.11, 111.91, 106.82, 65.95, 56.47, 35.53, 34.44, 26.53. HRMS: m/z calcd for [M+Na]^+^ 542.1726, found 542.1738.

##### 4.2.3.6. Compound 4f (4-methoxy-*N*-methyl-3-(3-meth-ylenecyclobutane-1-carboxamido)-5-(5-phenylthiophen-2-yl)benzamide)

^1^H NMR (500 MHz, DMSO) *δ* 9.29 (s, 1H), 8.23 – 8.09 (m, 2H), 7.72 – 7.64 (m, 2H), 7.50 (d, *J* = 3.8 Hz, 1H), 7.45 (t, *J* = 7.8 Hz, 2H), 7.34 (d, *J* = 7.4 Hz, 1H), 7.23 (d, *J* = 3.8 Hz, 1H), 7.15 (s, 1H), 4.81 (p, *J* = 2.4 Hz, 2H), 3.95 (s, 3H), 3.49 (t, *J* = 8.1 Hz, 1H), 2.87 – 2.79 (m, 2H), 2.70 (d, *J* = 4.6 Hz, 3H). ^13^C NMR (126 MHz, DMSO) *δ* 173.54, 169.73, 149.83, 145.55, 143.60, 141.37, 134.06, 129.67, 128.16, 127.66, 127.30, 127.23, 125.63, 124.59, 121.11, 111.90, 106.83, 56.56, 35.54, 34.46, 26.61. HRMS: m/z calcd for [M+Na]^+^ 455.1406, found 455.1428.

##### 4.2.3.7. Compound 4g (6-methoxy-*N*-methyl-5-(3-meth-ylenecyclobutane-1-carboxamido)-4’-morpholino-[1,1’-biphenyl]-3-carboxamide)

^1^H NMR (500 MHz, DMSO) *δ* 9.19 (s, 1H), 8.08 (s, 1H), 7.79 (q, *J* = 4.6 Hz, 1H), 7.34 – 7.25 (m, 2H), 7.01 – 6.91 (m, 3H), 4.80 (p, *J* = 2.4 Hz, 2H), 3.91 (s, 3H), 3.76 (dd, *J* = 6.1, 3.6 Hz, 4H), 3.46 (t, *J* = 8.1 Hz, 1H), 3.18 – 3.12 (m, 4H), 2.86 – 2.78 (m, 2H), 2.59 (d, *J* = 4.6 Hz, 3H). ^13^C NMR (126 MHz, DMSO) *δ* 173.33, 170.22, 150.55, 150.19, 145.63, 135.50, 131.23, 129.38, 125.95, 121.73, 115.05, 112.38, 106.78, 66.59, 56.38, 48.74, 35.54, 34.41, 26.49. HRMS: m/z calcd for [M+Na]^+^ 458.2056, found 458.2061.

##### 4.2.3.8. Compound 4h (4-methoxy-*N*-methyl-3-(3-meth-ylenecyclobutane-1-carboxamido)-5-(2-methylthiophen-3-yl)benzamide)

^1^H NMR (500 MHz, DMSO) *δ* 9.22 (s, 1H), 8.18 (s, 1H), 7.64 (d, *J* = 4.7 Hz, 1H), 7.26 (d, *J* = 5.2 Hz, 1H), 6.92 – 6.85 (m, 2H), 4.81 (p, *J* = 2.3 Hz, 2H), 3.88 (s, 3H), 3.47 (t, *J* = 8.1 Hz, 1H), 2.87 – 2.79 (m, 2H), 2.58 (d, *J* = 4.6 Hz, 3H), 2.32 (s, 3H). ^13^C NMR (126 MHz, DMSO) *δ* 173.40, 169.42, 150.02, 145.60, 137.52, 134.68, 130.63, 130.29, 130.00, 126.47, 121.74, 113.34, 106.81, 60.24, 56.48, 35.53, 26.56, 14.10. HRMS: m/z calcd for [M+Na]^+^ 393.1249, found 393.1248.

##### 4.2.3.9. Compound 4i (3-(5-acetylthiophen-2-yl)-4-meth-oxy-*N*-methyl-5-(3-methylenecyclobutane-1-carboxamido)benzamide)

^1^H NMR (500 MHz, DMSO) *δ* 9.34 (s, 1H), 8.24 (d, *J* = 4.8 Hz, 1H), 8.20 (s, 1H), 7.90 (d, *J* = 3.9 Hz, 1H), 7.29 (d, *J* = 3.9 Hz, 1H), 7.17 (s, 1H), 4.83 – 4.79 (m, 2H), 3.95 (s, 3H), 3.50 (t, *J* = 8.1 Hz, 1H), 2.84 (d, *J* = 9.3 Hz, 2H), 2.67 (d, *J* = 4.6 Hz, 3H), 2.55 (s, 3H). ^13^C NMR (126 MHz, DMSO) *δ* 191.12, 173.68, 169.36, 150.43, 145.50, 143.79, 134.60, 133.64, 132.83, 132.51, 131.93, 129.29, 127.76, 112.34, 106.87, 56.64, 35.53, 34.47, 26.87, 26.58. HRMS: m/z calcd for [M-H]^-^ 397.1300, found 397.1204.

##### 4.2.3.10. Compound 4j (methyl 4-(2-methoxy-5-(methylcarbamoyl)-3-(3-methylenecyclobutane-1-carboxamido)phenyl)thiophene-2-carboxylate)

^1^H NMR (500 MHz, DMSO) *δ* 9.26 (s, 1H), 8.14 (s, 1H), 8.08 (d, *J* = 4.7 Hz, 1H), 7.90 (d, *J* = 22.2 Hz, 2H), 7.14 (s, 1H), 4.81 (s, 2H), 3.94 (s, 3H), 3.86 (s, 3H), 2.92 – 2.89 (m, 1H), 2.86 – 2.78 (m, 2H), 2.64 (d, *J* = 4.6 Hz, 3H).^13^C NMR (126 MHz, DMSO) *δ* 173.50, 169.87, 162.38, 150.05, 145.57, 141.55, 134.53, 132.61, 130.65, 129.44, 128.64, 127.10, 121.13, 112.28, 106.82, 56.56, 52.78, 35.53, 34.44, 26.51. HRMS: m/z calcd for [M+Na]^+^ 437.1147, found 437.1141.

##### 4.2.3.11. Compound 4k (3-(5-ethylthiophen-2-yl)-4-meth-oxy-N-methyl-5-(3-methylenecyclobutane-1-carboxamido)benzamide)

^1^H NMR (500 MHz, DMSO) *δ* 9.24 (s, 1H), 8.08 (t, *J* = 2.3 Hz, 2H), 7.06 – 7.00 (m, 2H), 6.81 (d, *J* = 3.6 Hz, 1H), 4.80 (t, *J* = 2.4 Hz, 2H), 3.91 (s, 3H), 3.47 (t, *J* = 8.1 Hz, 1H), 2.83 (dd, *J* = 7.5, 1.0 Hz, 4H), 2.66 (d, *J* = 4.6 Hz, 3H), 1.27 (t, *J* = 7.5 Hz, 3H). ^13^C NMR (126 MHz, DMSO) *δ* 173.45, 169.83, 149.79, 147.41, 145.57, 139.12, 129.43, 127.80, 126.80, 126.05, 124.65, 121.14, 111.79, 106.81, 56.46, 35.53, 34.43, 26.55, 25.44, 24.97, 23.24, 16.28. HRMS: m/z calcd for [M+Na]^+^ 407.1406, found 407.1413.

##### 4.2.3.12. Compound 4l (4-methoxy-N-methyl-3-(5-methylbenzo[b]thiophen-2-yl)-5-(3-methylenecyclobutane-1-carboxamido)benzamide)

^1^H NMR (500 MHz, DMSO) *δ* 9.32 (s, 1H), 8.23 – 8.14 (m, 2H), 7.84 (d, *J* = 8.2 Hz, 1H), 7.65 (s, 1H), 7.42 (s, 1H), 7.20 (d, *J* = 8.5 Hz, 2H), 4.81 (p, *J* = 2.3 Hz, 2H), 3.95 (s, 3H), 3.50 (t, *J* = 8.1 Hz, 1H), 2.87 – 2.80 (m, 2H), 2.67 (d, *J* = 4.6 Hz, 3H), 2.44 (s, 3H). ^13^C NMR (126 MHz, DMSO) *δ* 173.61, 169.54, 149.78, 145.53, 142.37, 140.68, 137.05, 134.24, 130.26, 127.73, 126.50, 123.93, 122.28, 121.08, 112.54, 106.85, 56.57, 35.54, 34.47, 26.61, 21.48. HRMS: m/z calcd for [M+Na]^+^ 443.1406, found 443.1395.

### 4.3. Purity assessment

The purity of the key compound reported in this study (**4a**) was determined to be >95% as assessed by LC-MS (see Figure S9).

### 4.4. Biophysical evaluation

#### 4.4.1. SPR for TREM2 binding assessment

Binding affinities of the **4a** and **4i** for biotinylated human TREM2 (Cat.: 11084-H49H-B, Sino Biological, Beijing, China) were determined by SPR (Biacore 8K, Cytiva). The biotinylated TREM2 protein was immobilized on SA Sensor Chip (Cytiva) in a PBS-P (0.2 M phosphate buffer with 27 mM KCl, 1.37 M NaCl and 0.5% Surfactant P20, pH 7.4, Cytiva) to a response level of 4,343 RU ± 399 RU. Kinetic measurements were run using a single cycle kinetic approach. A two-fold dilution series of each compound, consisting of seven concentrations ranging between 200 µM and 3.12 µM, were prepared in running buffer (PBS-P supplemented 2% DMSO) and injected over the prepared surface of the SA Sensor Chip. Experiments were conducted at 25°C employing the following parameters: flow rate of 30 µl/min, contact time of 200 s, and dissociation time of 900 s. After each analysis an additional wash with 50% DMSO solution was performed. Each interaction was investigated at least in triplicate. The results are presented as sensorgrams obtained after subtraction of the background response signal from a reference flow cell and from a control experiment with buffer injection. The obtained data were analyzed using Biacore^™^ Insight Evaluation Software (Cytiva). Binding parameters (k_a_, k_d_, K_D_) were obtained using 1:1 binding model.

#### 4.4.2. Temperature-Related Intensity Change (TRIC)

Temperature-Related Intensity Change measurements were performed using a Dianthus NT.23PicoDuo instrument (NanoTemper Technologies, Munich, Germany) based on the Temperature-Related Intensity Change (TRIC) method. Recombinant TREM2 protein (BioTechne, MN, USA) was labeled using a His-Tag Labeling Kit RED-tris-NTA 2nd Generation (NanoTemper Technologies, Munich, Germany) according to the manufacturer’s protocol. All binding experiments were conducted in PBST buffer (154 mM NaCl, 5.6 mM Na2HPO4, 1.05 mM KH2PO4, pH 7.4, 0.005% Tween-20). For binding measurements, labeled TREM2 protein was used at a final concentration of 10 nM. Compounds were prepared at the indicated concentrations and incubated with labeled TREM2 for 10 minutes at room temperature prior to measurements. The Dianthus instrument was configured with the following parameters: 85% LED excitation power, picomolar detector sensitivity off, and a laser on-time of 5 seconds. All measurements were performed in triplicate. Initial data processing was performed using the Dianthus Analysis software and subsequent data analysis and curve fitting were conducted using GraphPad Prism (GraphPad Software, San Diego, CA, USA).

#### 4.4.3. MST for TREM2 binding assessment

Binding affinity measurements were performed using MST on a Monolith NT.115 system (NanoTemper Technologies, Munich, Germany). TREM1 and TREM2 proteins (BioTechne, MN, USA) were labeled with RED-trisNTA His-tag labeling kit (NanoTemper Technologies, Munich, Germany) according to manufacturer’s instructions. TREM1 assays used buffer containing 154 mM NaCl, 5.6 mM Na_2_HPO_4_, 1.05 mM KH_2_PO_4_, pH 7.4 with 0.05% Tween-20, while TREM2 assays used the same buffer with 0.005% Tween-20. Labeled proteins (40 nM) were incubated with compounds at indicated concentrations for 10 minutes at room temperature. Measurements were performed using red filter, 100% LED power, and medium MST power. Data were analyzed with MO. Affinity Analysis software and GraphPad Prism (GraphPad Software, CA, USA) to determine KD values.

### 4.5. Cellular assay

#### 4.4.1 Cell viability assay

The cytotoxic effects of test compounds were evaluated using the MTS assay (Cell Titer 96® AQueous One Solution, Promega, WI, USA). HMC3 microglial cells were seeded in 96-well plates and maintained in complete growth medium containing 10% FBS until reaching approximately 40% confluence (24 h). After washing, cells were serum-deprived in FBS-free medium for 4 h. Cells were then treated with compounds and Tween-20 for 1 h followed by MTS analysis according to the supplier’s protocol. Absorbance readings were obtained at 490 nm using an Infinite M1000 Pro Microplate Reader (Tecan, Männedorf, Switzerland).

#### 4.4.2. AlphaLISA pSyk (525/526) assay

Phospho-AlphaLISA assay measures a protein target when phosphorylated at a specific residue. The assay uses two antibodies which recognize the phospho epitope and a distal epitope on the targeted protein. In the presence of phosphorylated protein, luminescent Alpha signal is generated. The amount of light emission is directly proportional to the quantity of phosphoprotein present in the sample (from Revvity website). Briefly, HEK – hTREM2/DAP12 cells were seeded at 50,000 cells per well in 96-well plate, in a final volume of 100 μl of growth media composed of DMEM (Gibco) supplemented with 10% FBS (Gibco) and incubated for 24 h at 37 °C, 5% CO_2_. Compounds **4a** and **4i** were diluted in media at a concentration of 25uM, growth media was manually removed from the wells and subsequently replaced with the test compounds containing media. Control wells were treated with the vehicle buffer used to solubilize the several compounds. After 1h, the media was gently removed, lysis buffer was added and after complete lysis, alphalisa assay was performed according to manufacturer instructions (Revvity, USA).

#### 4.4.3. Phagocytosis Assay

Phagocytosis assay was performed in BV2 cells using green fluorescent latex beads (Sigma #L1030). Prior use, beads were opsonized in FBS (1 h at 37 °C), the final concentrations for beads and FBS in DMEM were 0.01% (v/v) and 0.05% (v/v) respectively. Briefly, BV2 cells were submitted to serum starvation prior treatment with either compound **4a, 4i** (25 μM for 30 min) or solubilization buffer (control condition) and beads were added to the media for an additional 30 min. Cultures were then washed 3 times with ice-cold PBS and fixed in 4% paraformaldehyde (PFA) (Thermofisher, MA, USA) before undergoing immunocytochemistry. Following Immunostaining, pictures were taken for each condition under 20X objective, total number of cells was counted manually using DAPI staining and each IBA1 immunolabelled-cell containing at least one bead was counted as positive for phagocytosis.

#### 4.4.4. Immunocytochemistry

Fixed cells were permeabilized with 0.25% Triton X-100-containing PBS and subsequently blocked with 1% BSA. Immunocytochemistry was then performed using primary anti IBA1 (Wako, Richmond, VA) antibody followed by secondary 594 Alexa fluorescent antibody (Thermofisher, MA, USA).

### 4.5. Computational assay

The crystal structure of human TREM2 was obtained from the Protein Data Bank (PDB ID: 5UD7, accessed February 2025).^24^ Protein preparation was performed using the Protein Preparation Wizard in Maestro Schrödinger Suite (v2021.2). This involved the addition of missing hydrogen atoms, assignment of bond orders, optimization of hydrogen-bonding networks, and removal of crystallographic water molecules except those mediating critical interactions. Protonation states of ionizable residues were adjusted to reflect physiological pH (7.4) using PROPKA. This was followed by energy minimization with the OPLS3e force field to relieve steric clashes.

A 1ms MD simulation of **4a**/TREM2 complex was conducted using the DESMOND module (Schrödinger Suite) under the same conditions as our previous work.^25^ For validation, a second 250 ns MD simulation was carried out using GROMACS (v2024.3) under the CHARMM27 all-atom force field (CHARM22 plus CMAP for proteins) according to the same methodology of our previous study.^26^ This crossvalidation strategy mitigates software-specific artifacts and confirms the robustness of observed conformational dynamics. Data Analysis and Visualization were carried out using the VMD, Grace (V.5.1.) and discovery studio visualizer software.^27^ Gibbs free energy landscapes were reconstructed using in-house Python scripts based on principal component analysis (PCA) of backbone atoms.

## ASSOCIATED CONTENT

### Supporting Information

The Supporting Information is available free of charge on the ACS Publications website.

NMR spectra, mass spectrometry charts, purity assessment, binding curves, and chemical structure of **I-192** (PDF)

## AUTHOR INFORMATION

### Author Contributions

The manuscript was written through contributions of all authors. All authors have given approval to the final version of the manuscript.

## Abbreviations

AAV: Adeno-associated virus
AD: Alzheimer’s disease
Aβ: amyloid-beta
APOE: Apolipoprotein E
ka: association rate
CNS: central nervous system
DMAP: 4-dimethylaminopyridine
DEL: DNA-encoded library
k_d_: dissociation rate
K_D_: dissociation rate constant
HEK: Human Embryonic Kidney
Ig: immunoglobulin
MST: microscale thermophoresis
MD: molecular dynamics
PD-1: programmed cell death protein 1
SAR: Structure-activity relationship
sTREM2: soluble TREM2
SPR: surface plasmon resonance
TRIC: Temperature-Related Intensity Change
TREM2: Triggering Receptor Expressed on Myeloid cells 2

## Notes

The authors declare no competing financial interests.

