## Supporting Information for "TREM2 Activation by First-in-Class Direct Small Molecule Agonists: DEL Screening, Optimization, Biophysical Validation, and Functional Characterization"

Corresponding:

^*^Moustafa T. Gabr: Department of Radiology, Molecular Imaging Innovations Institute (MI3), Weill Cornell Medicine, New York, NY 10065, USA.

**Table of Content**

I. Supplementary figures	[Page 3](#_Toc189481170)

II. [Spectral data	Pages 5-30](#_Toc189481172)

**I. Supplementary Figures**

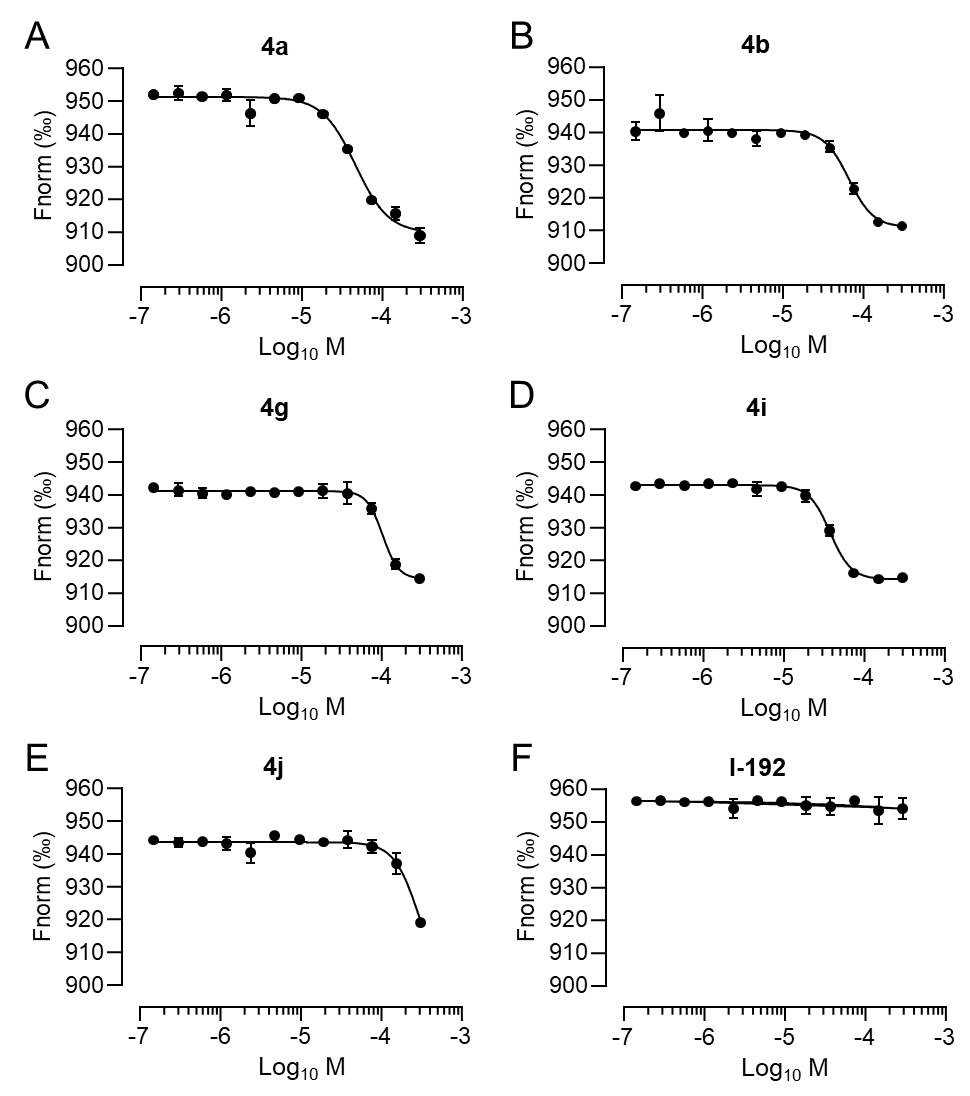

**Figure S1. Dose-dependent binding analysis of compound 4a and its four derivatives using Dianthus (A-F)** Dose-dependent binding experiments for 5 hit candidates and TREM2 agonist I-192 using Dianthus. Compounds were incubated at room temperature for 10 minutes at the indicated concentrations before measurements. Data are presented as Mean ± SEM (n = 3).

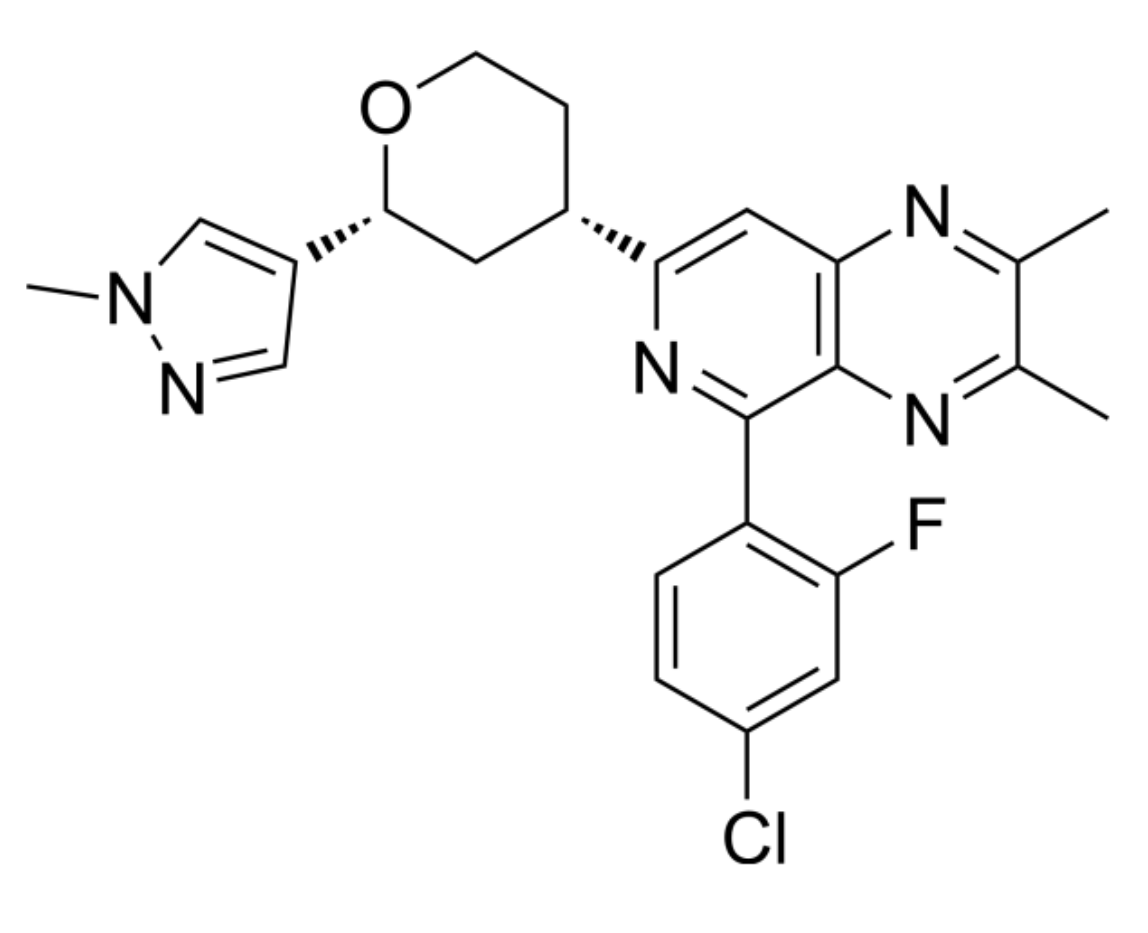

**Figure S2. Chemical structure of small molecule TREM2 agonist, I-192 (VG‐3927).**

**II. Spectral data**

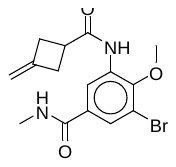

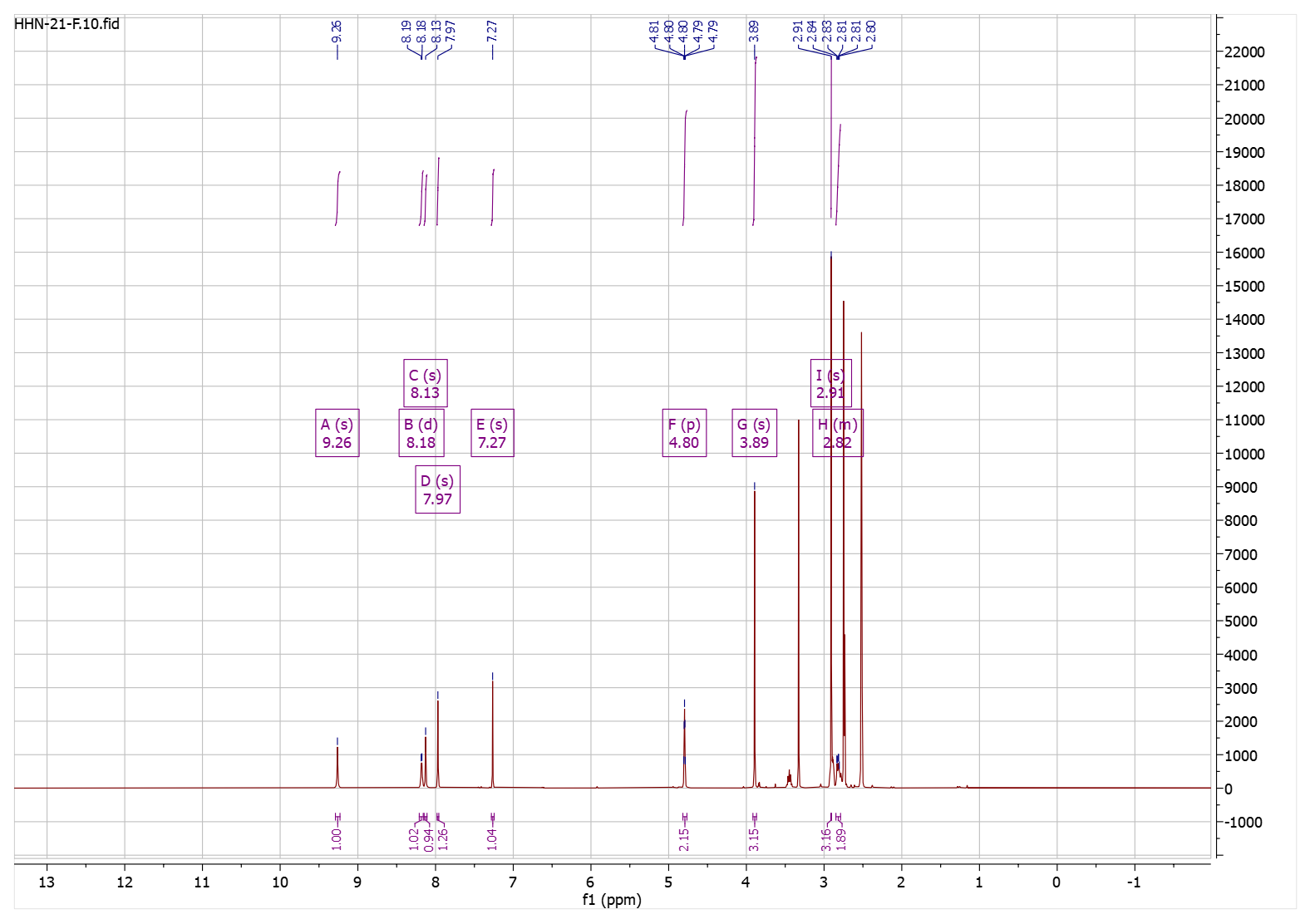

**Figure S3.** ^1^H-NMR spectrum of compound **3**.

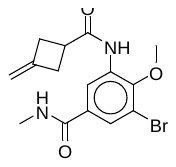

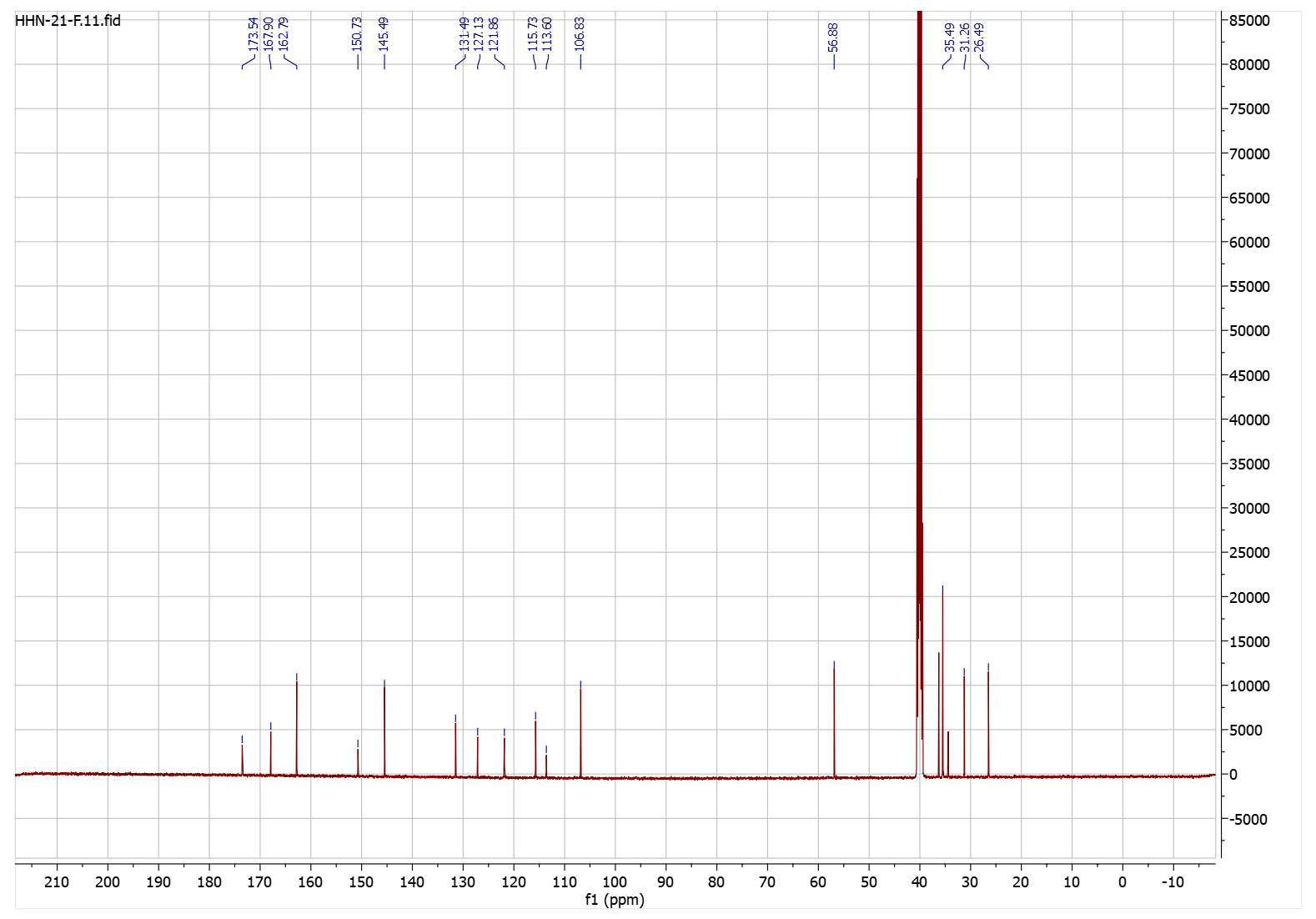

**Figure S4.** ^13^C-NMR spectrum of compound **3**.

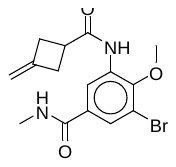

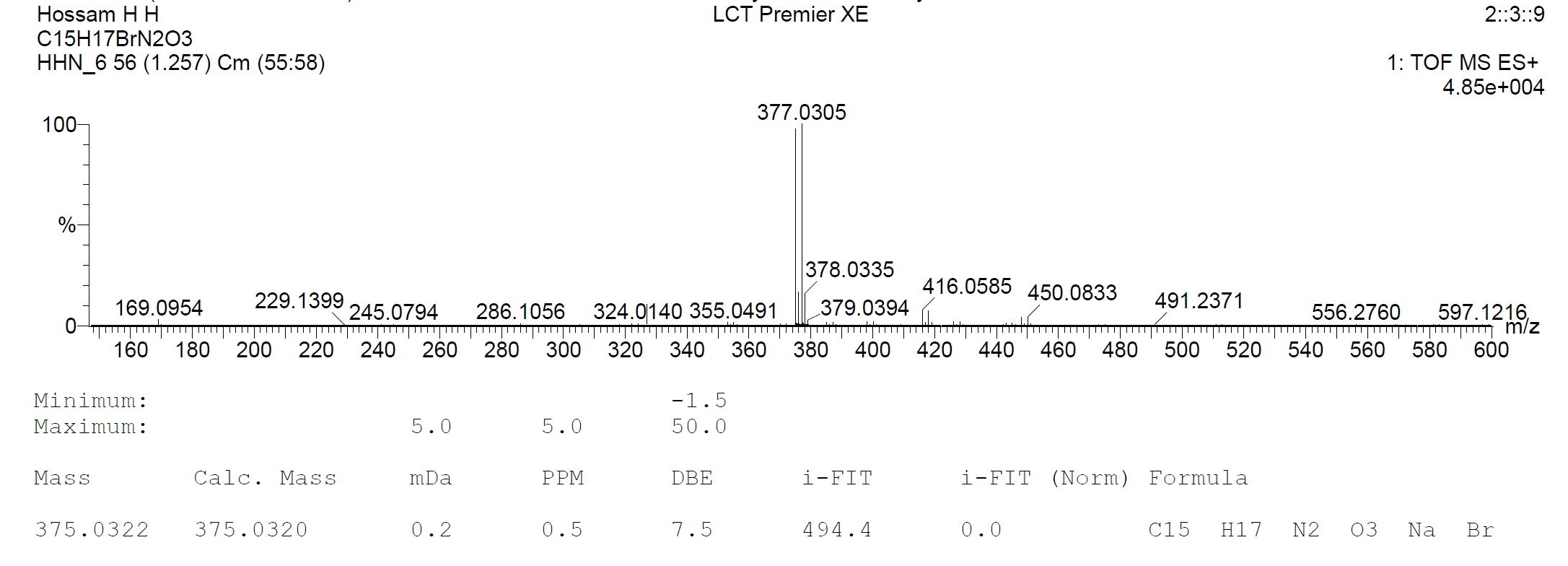

**Figure S5.** HRMS spectrum of compound **3**.

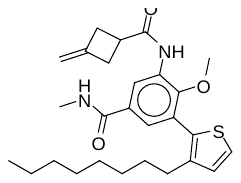

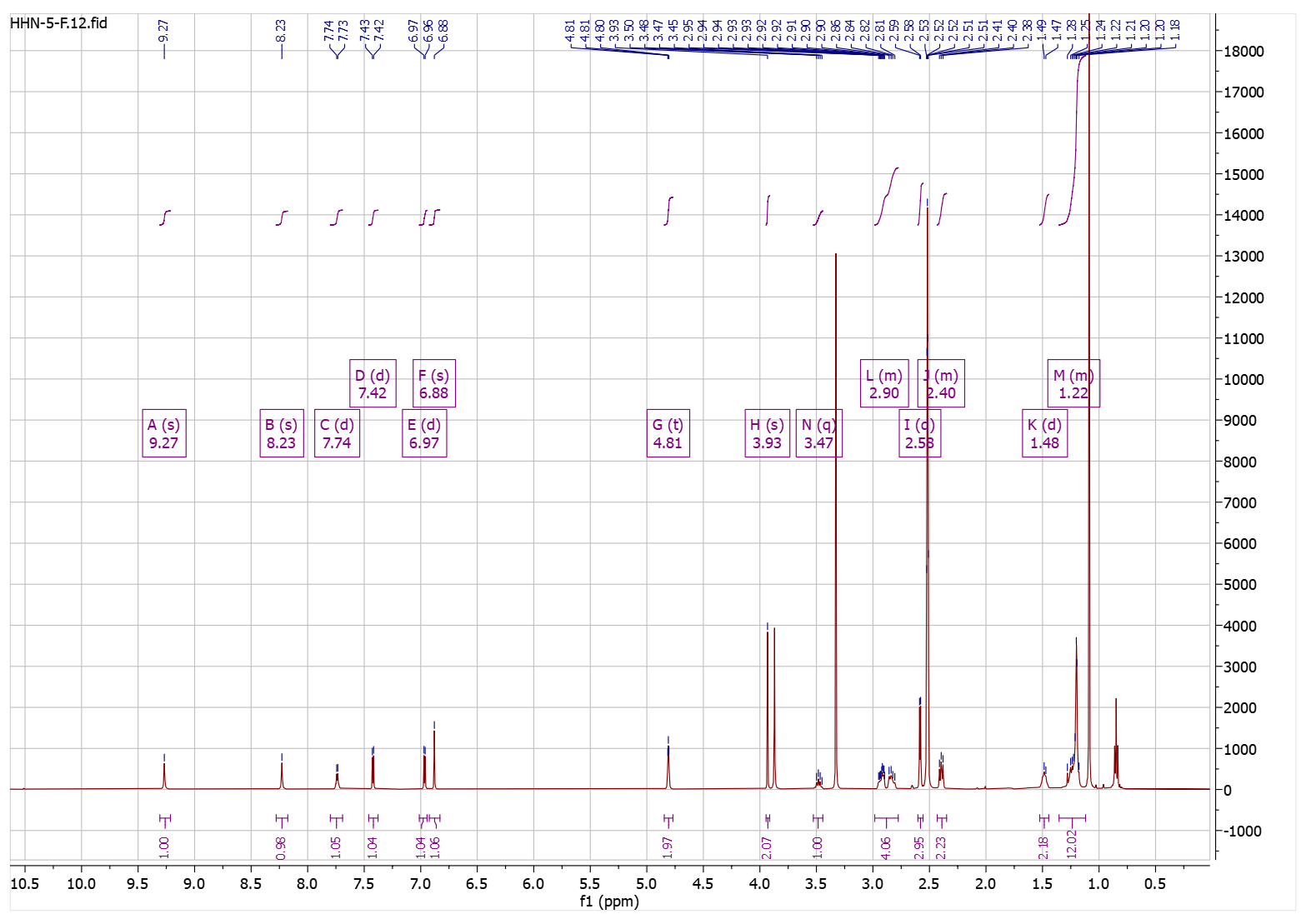

**Figure S6.** ^1^H-NMR spectrum of compound **4a**.

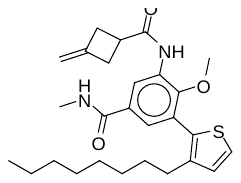

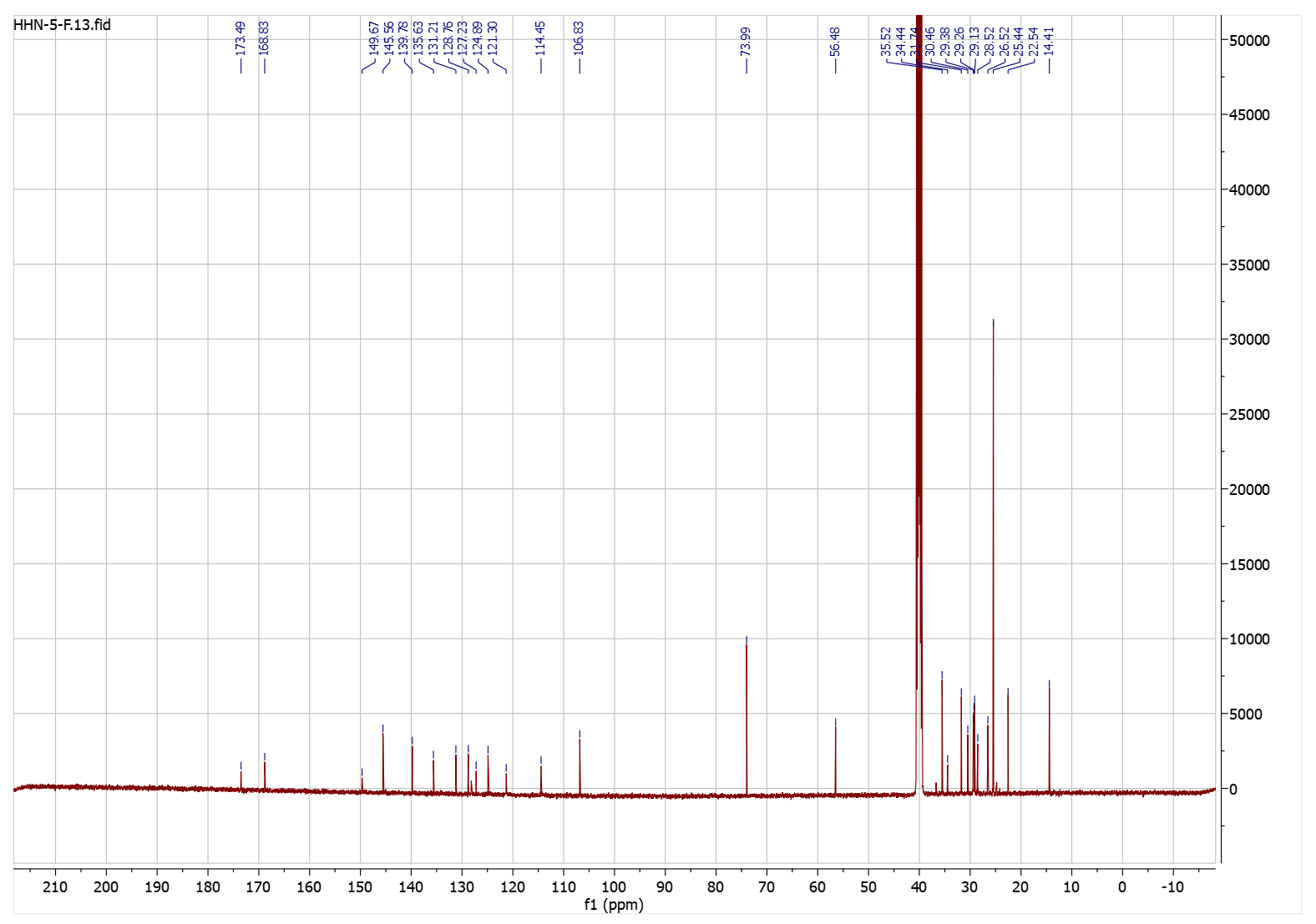

**Figure S7.** ^13^C-NMR spectrum of compound **4a**.

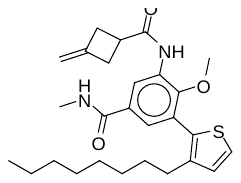

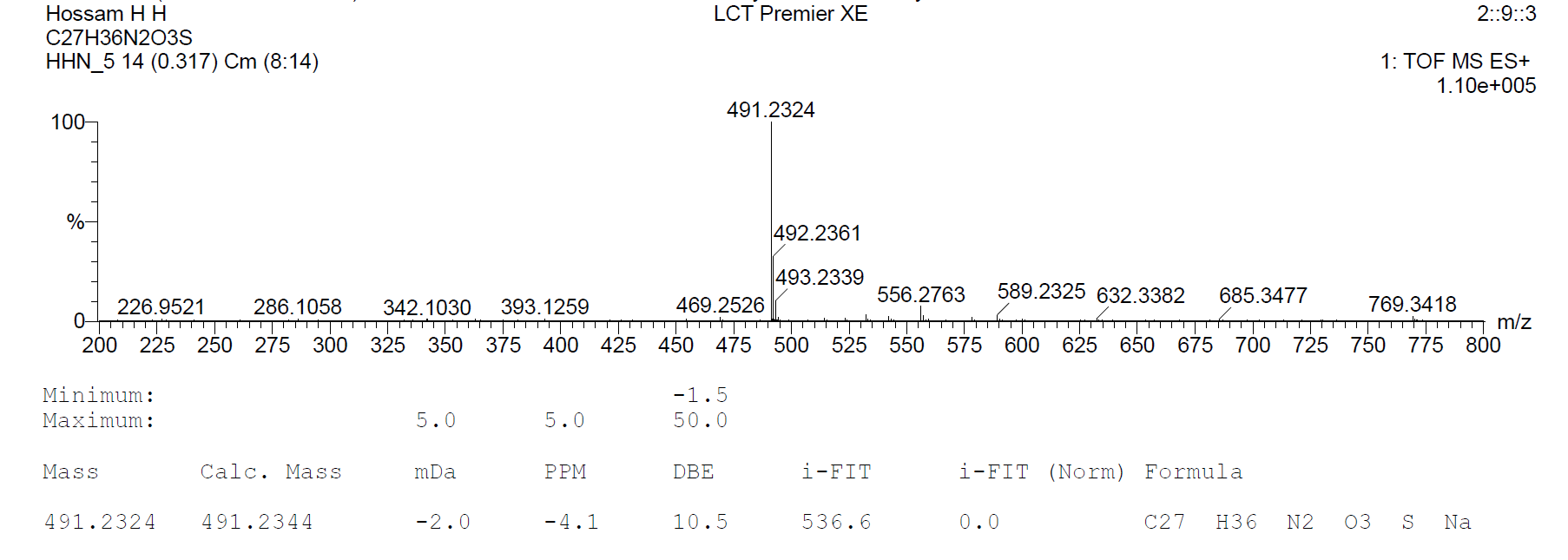

**Figure S8.** HRMS spectrum of compound **4a**.

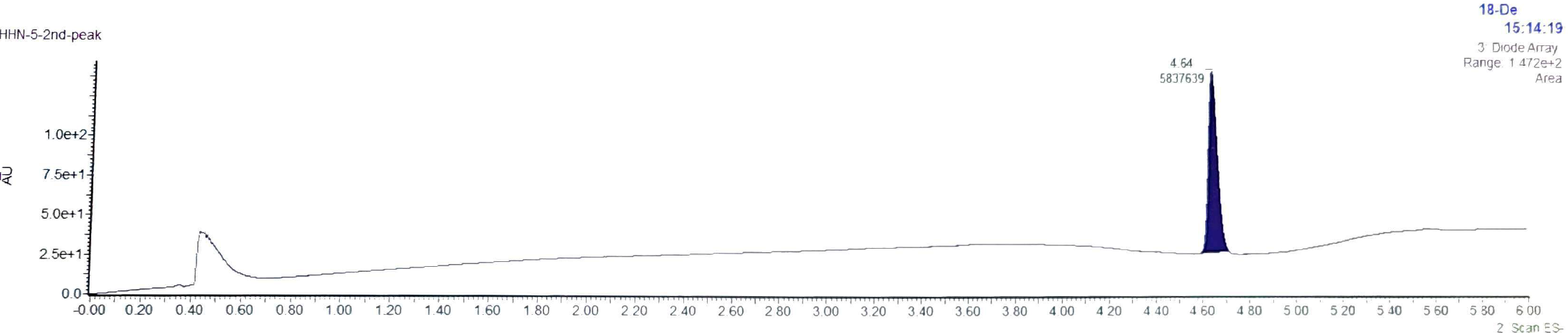

DMF

**Figure S9.** LCMS spectrum of compound **4a** showing 100% purity.

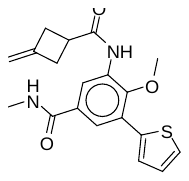

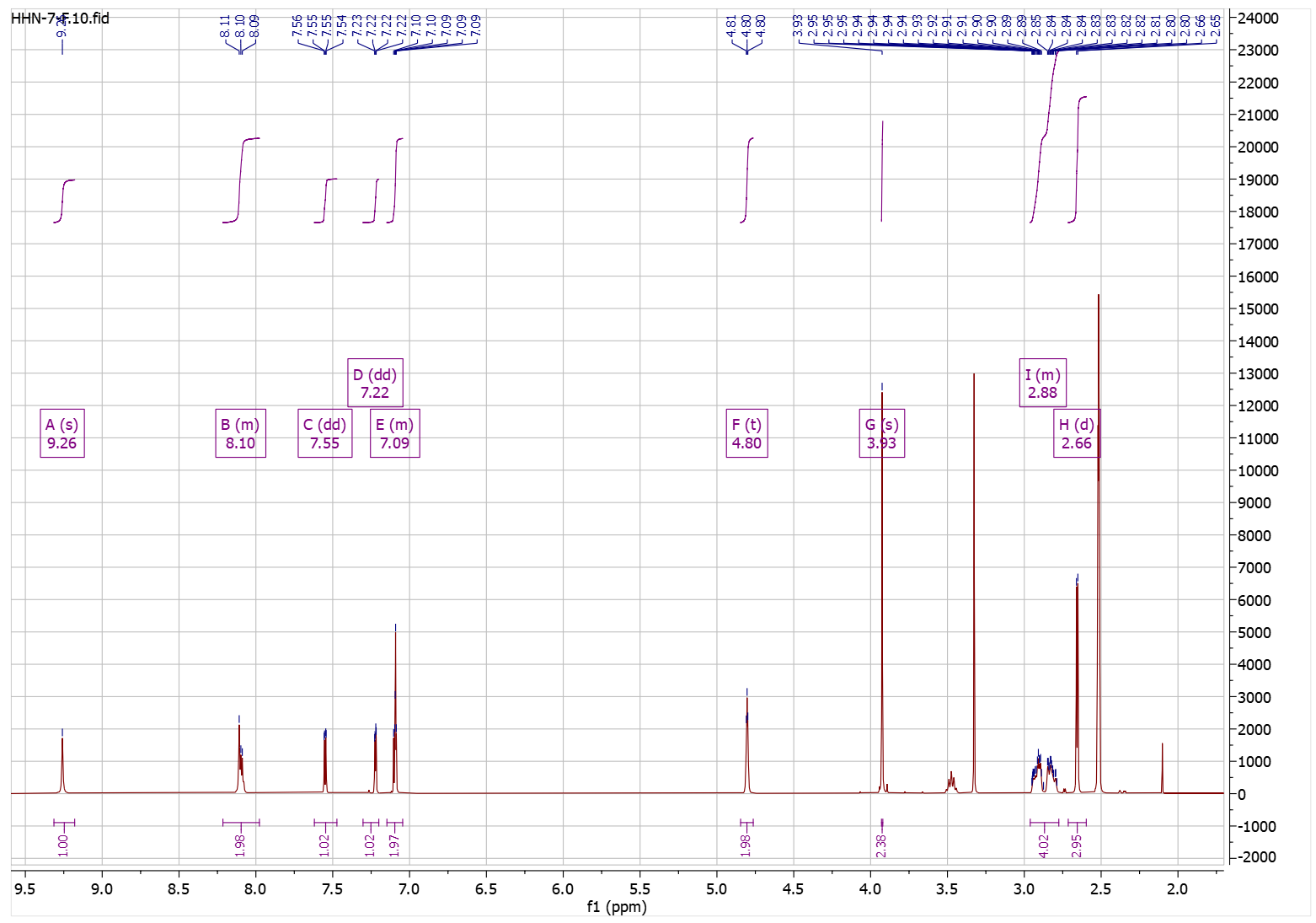

**Figure S10.** ^1^H-NMR spectrum of compound **4b**.

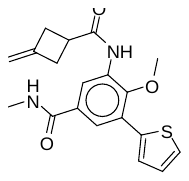

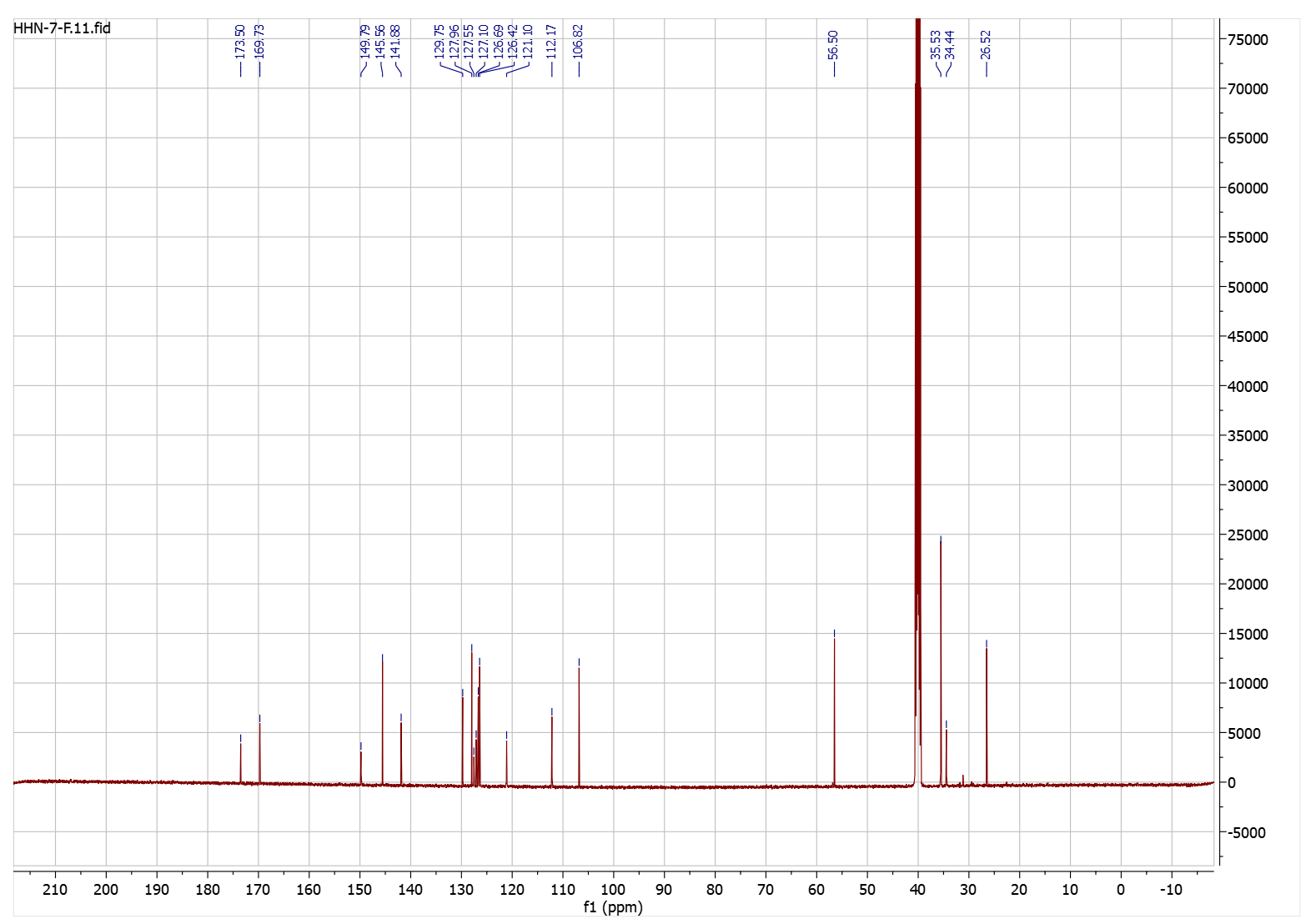

**Figure S11.** ^13^C-NMR spectrum of compound **4b**.

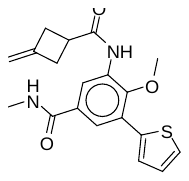

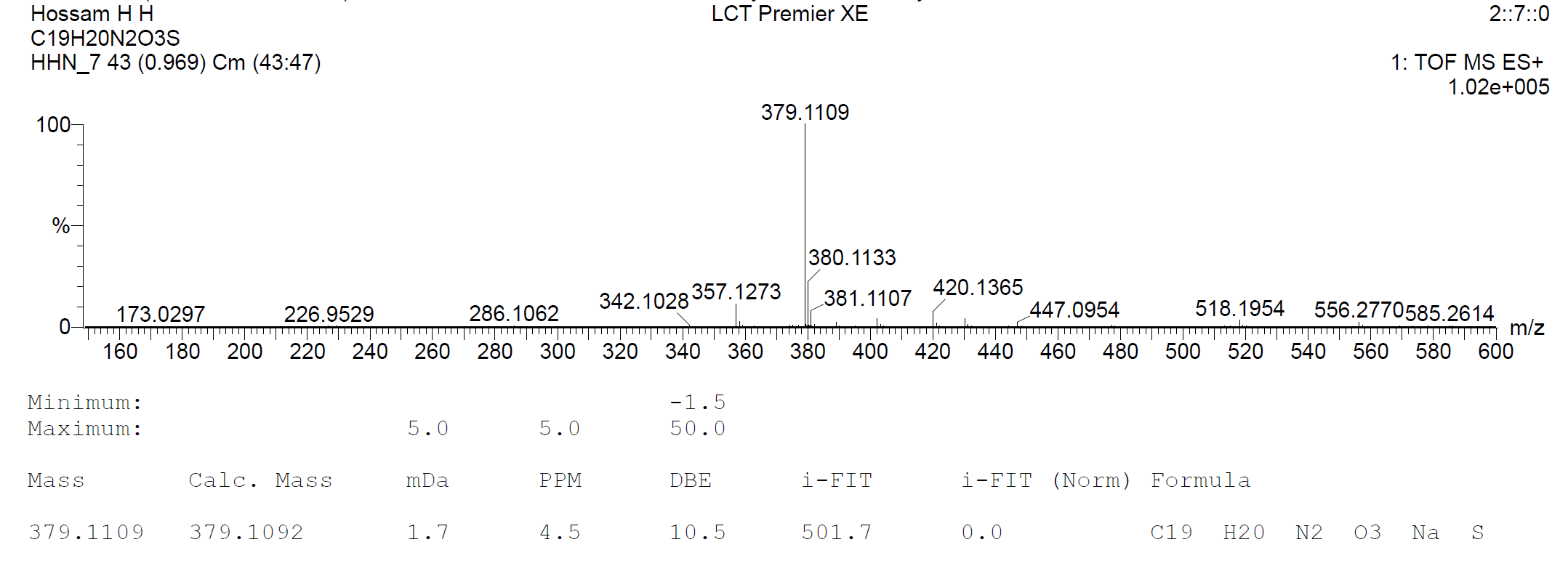

**Figure S12.** HRMS spectrum of compound **4b**.

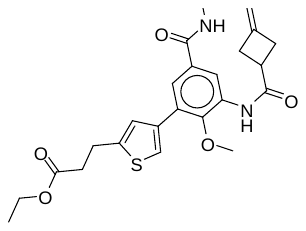

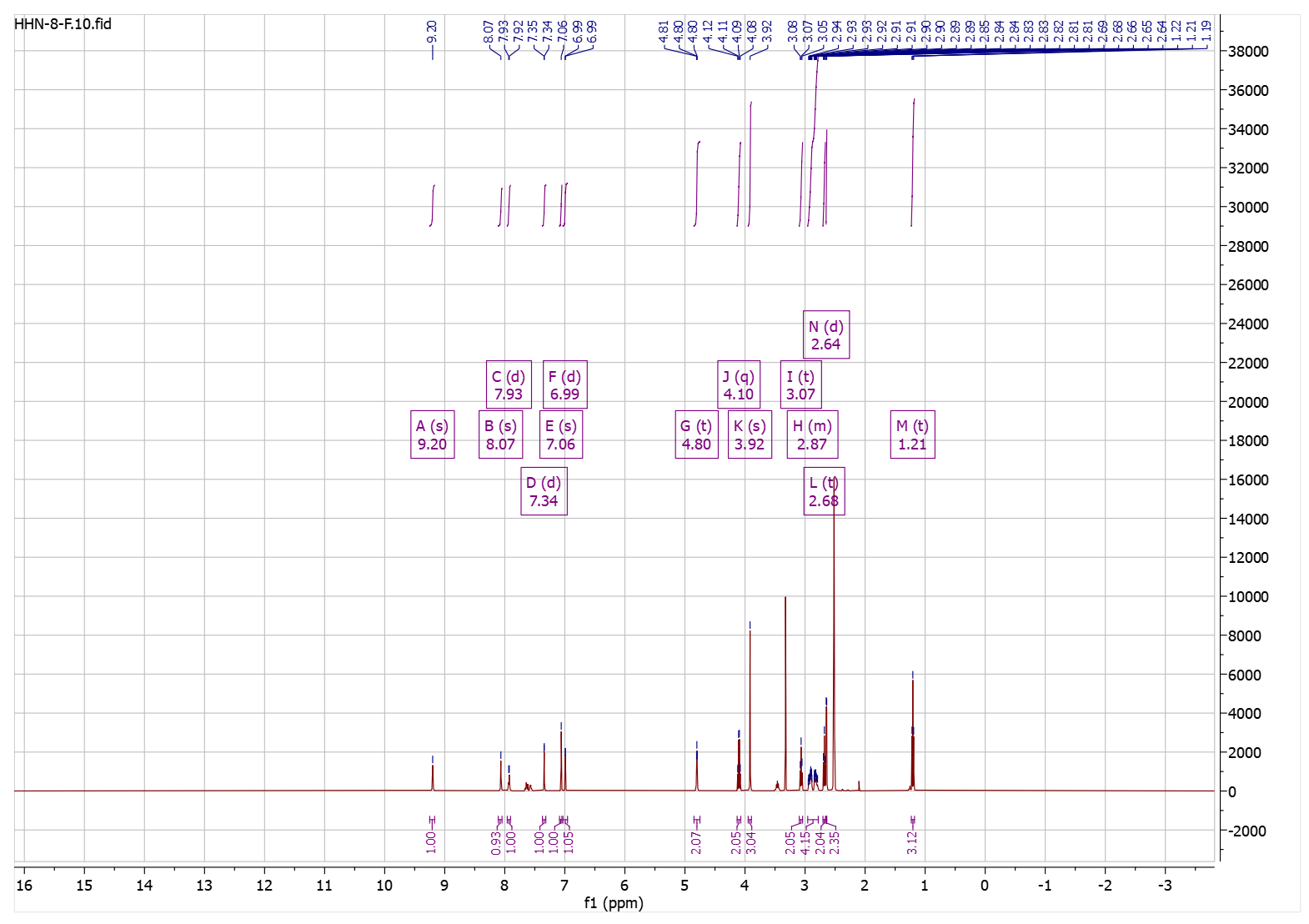

**Figure S13.** ^1^H-NMR spectrum of compound **4c**.

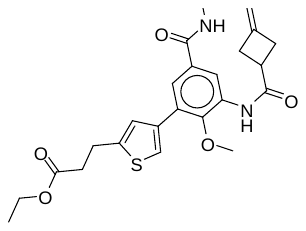

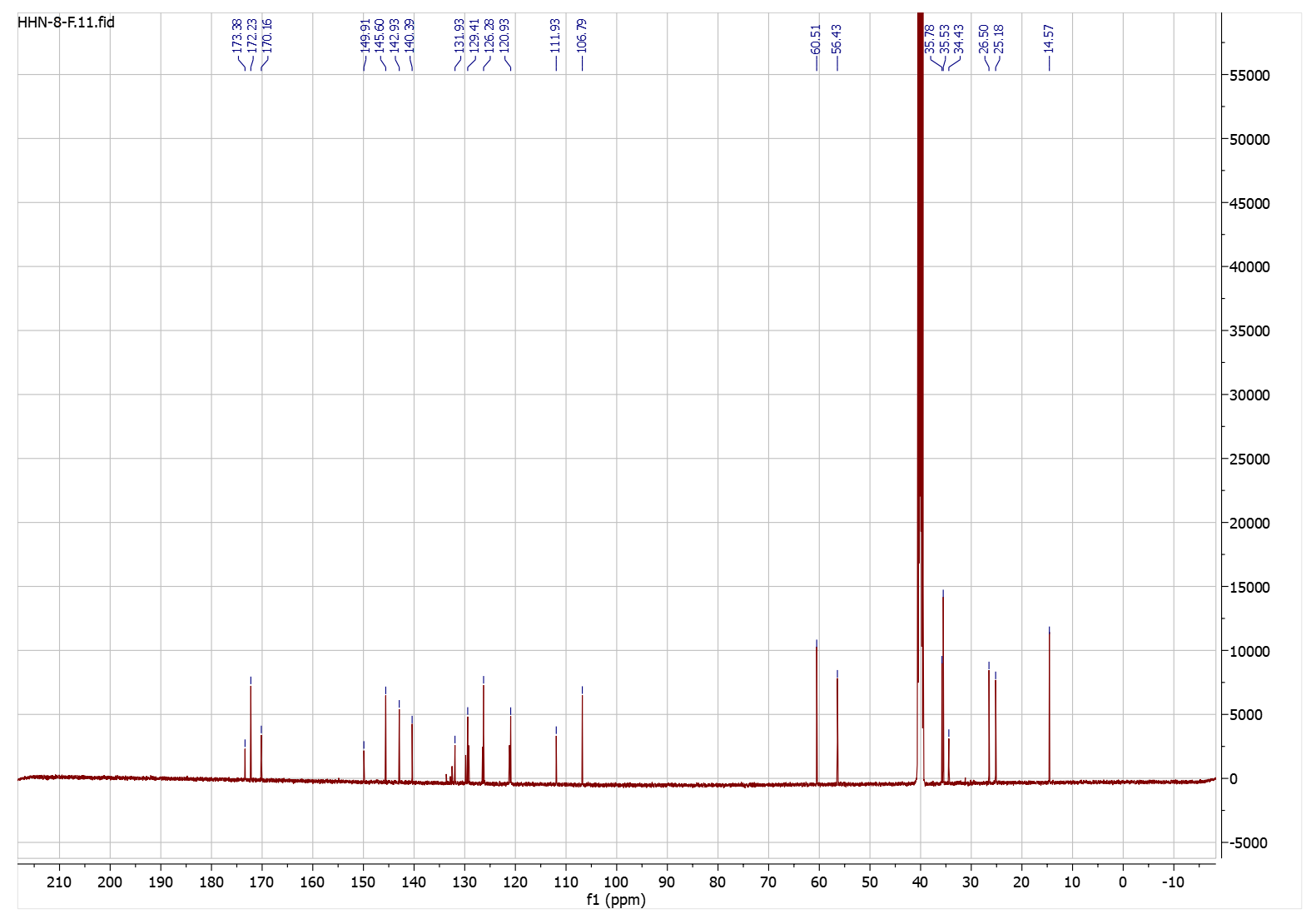

**Figure S14.** ^13^C-NMR spectrum of compound **4c**.

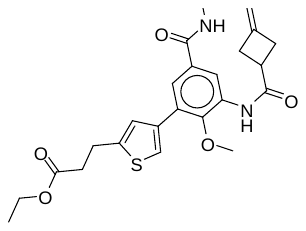

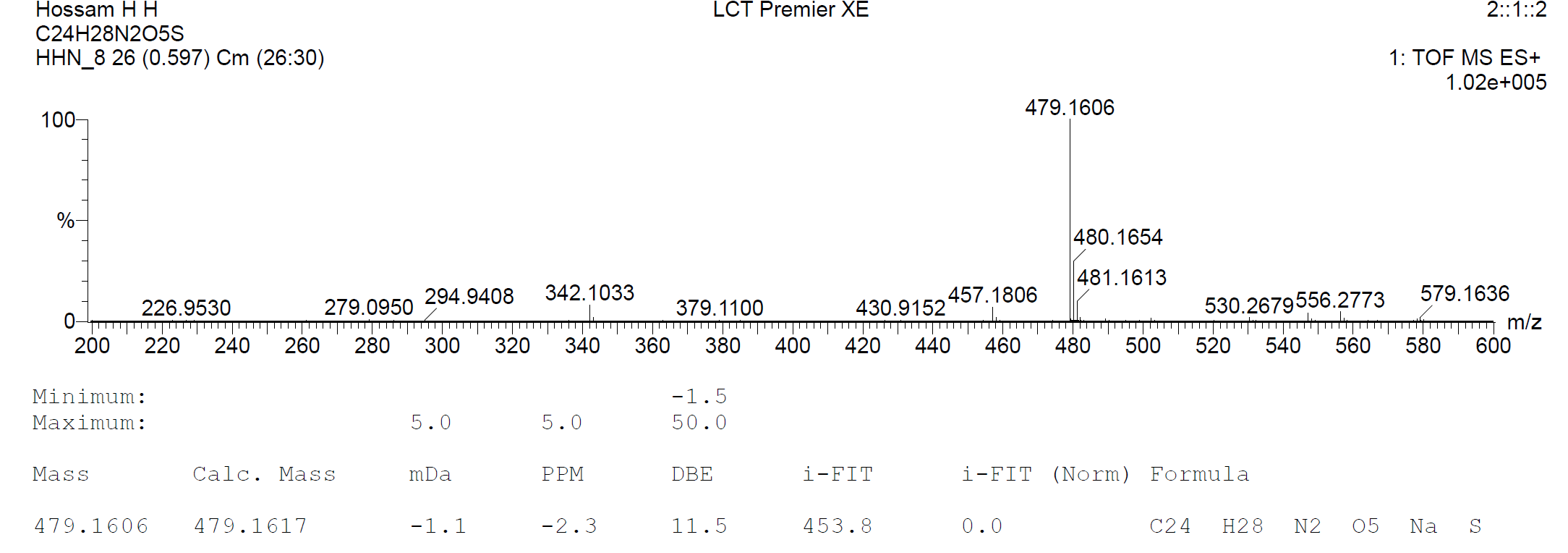

**Figure S15.** HRMS spectrum of compound **4c**.

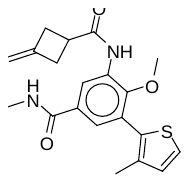

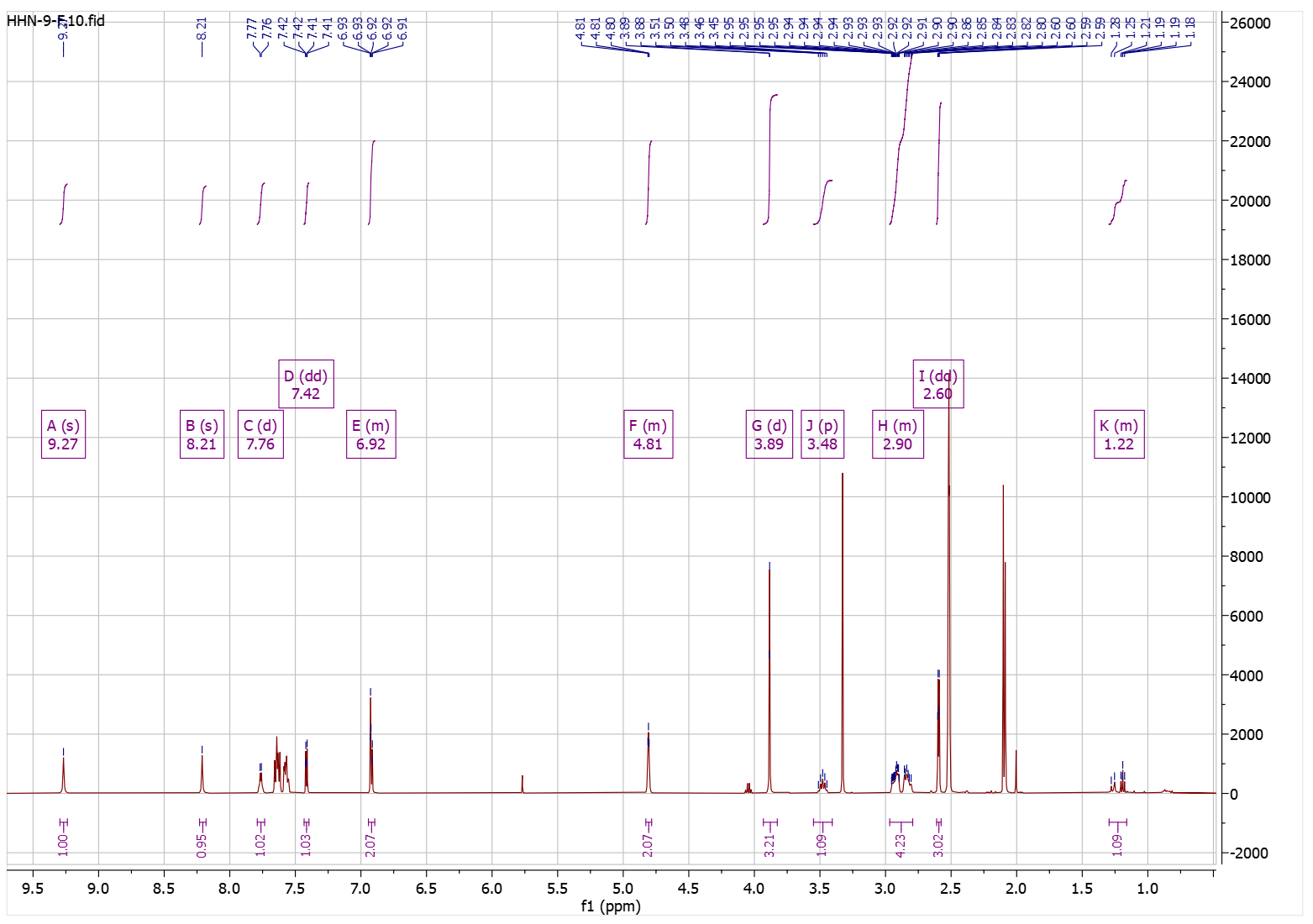

**Figure S16.** ^1^H-NMR spectrum of compound **4d**.

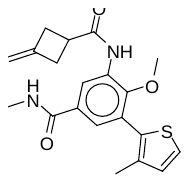

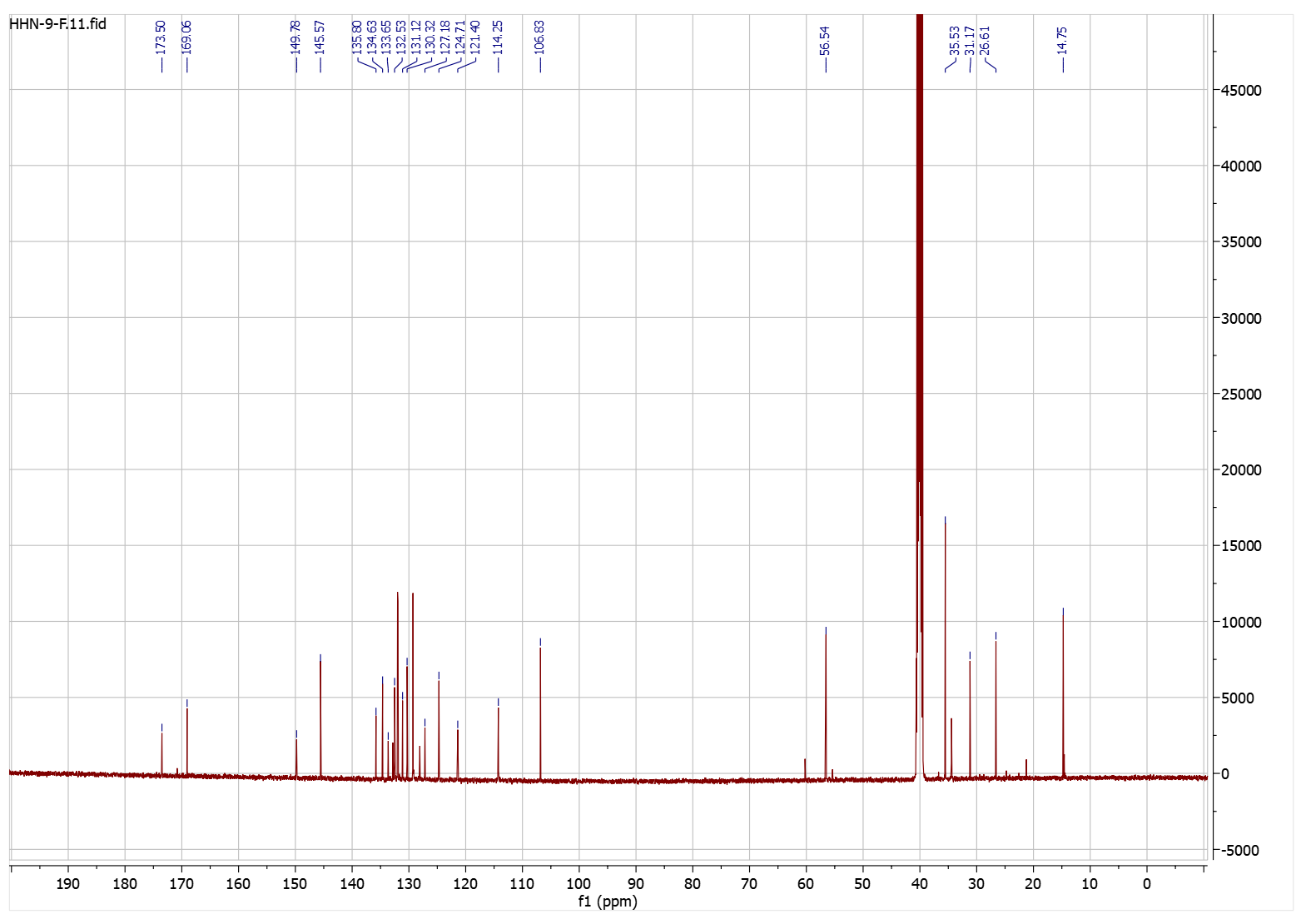

**Figure S17.** ^13^C-NMR spectrum of compound **4d**.

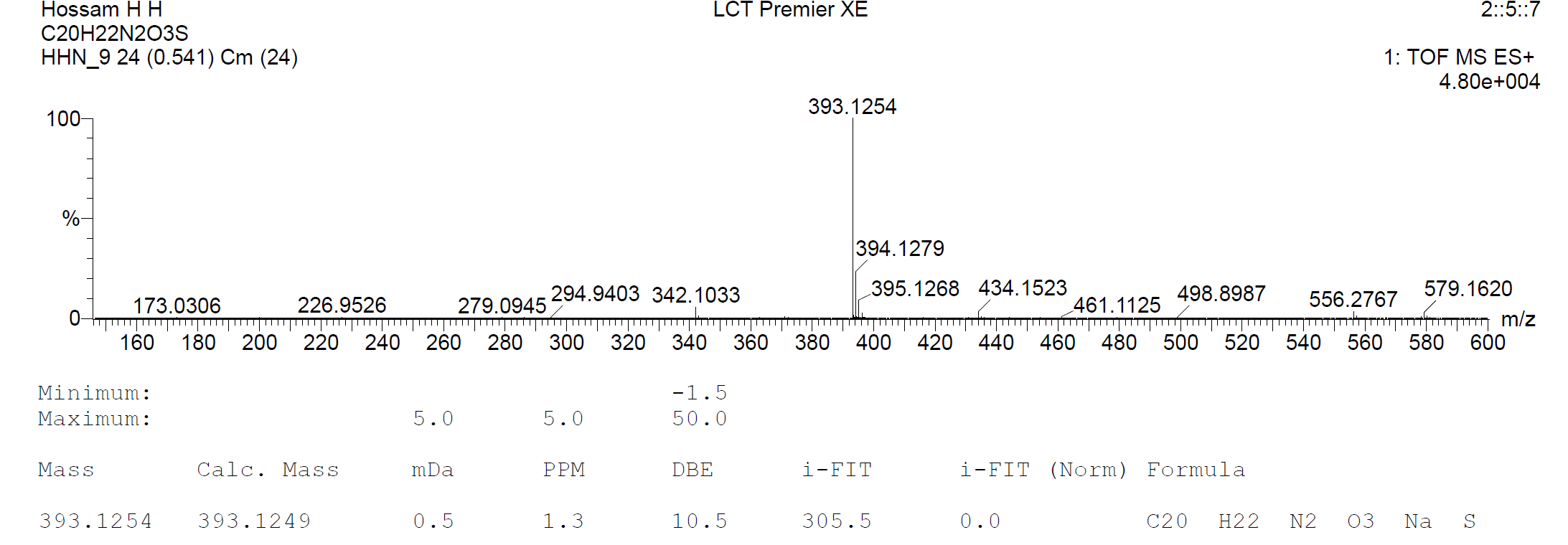

**Figure S18.** HRMS spectrum of compound **4d**.

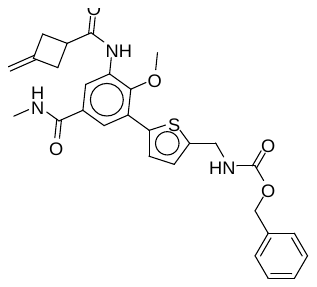

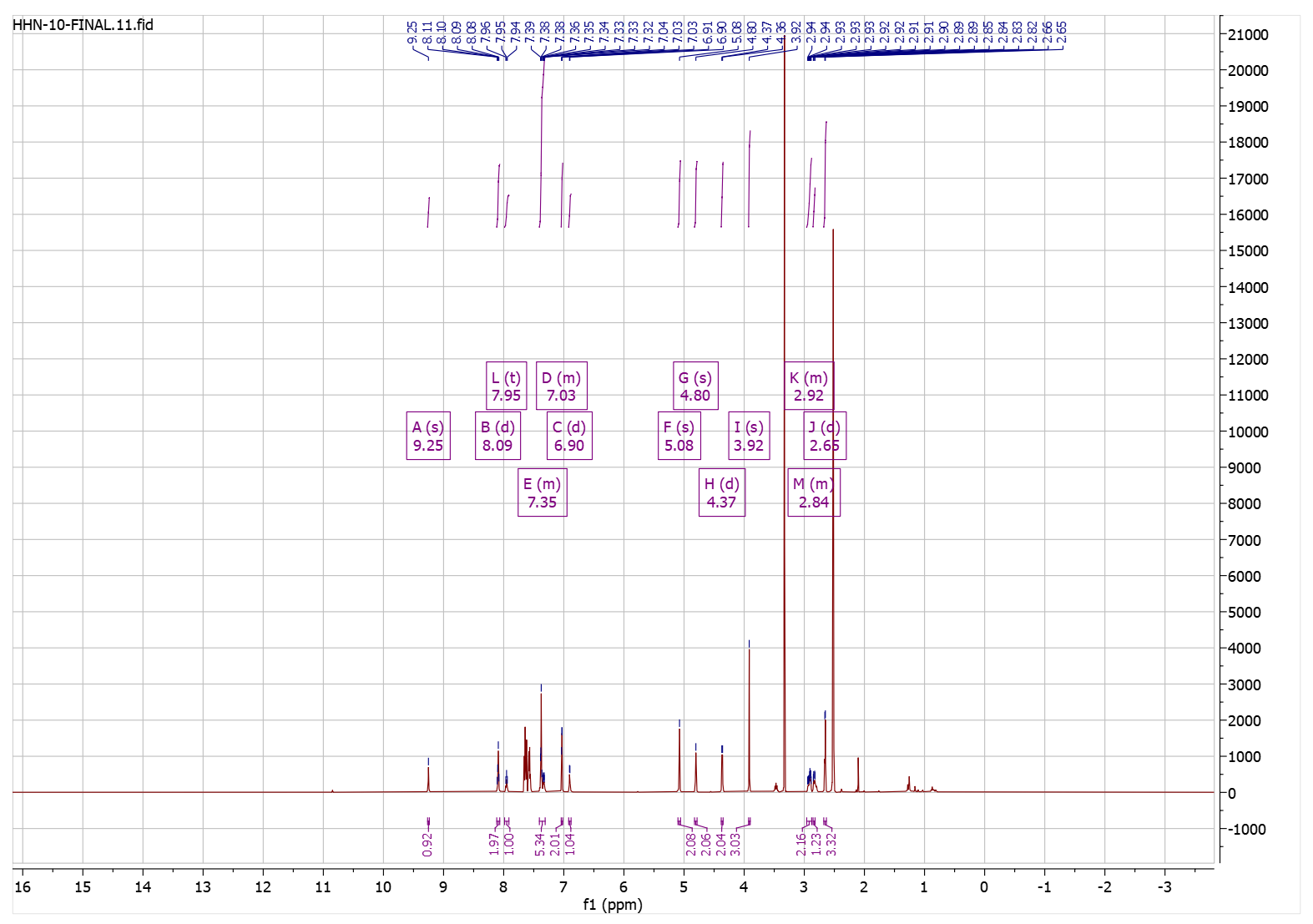

**Figure S19.** ^1^H-NMR spectrum of compound **4e**.

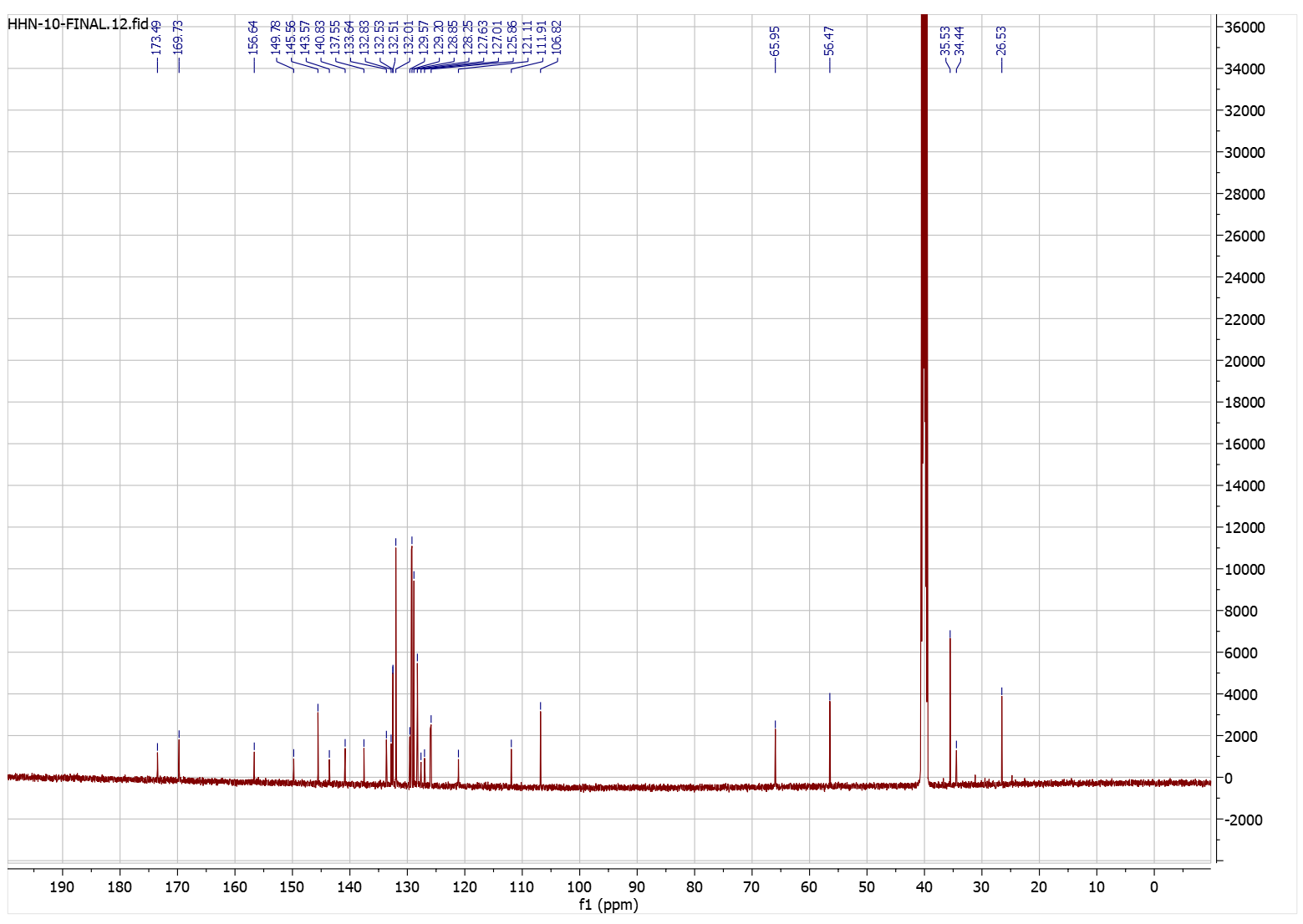

**Figure S20.** ^13^C-NMR spectrum of compound **4e**.

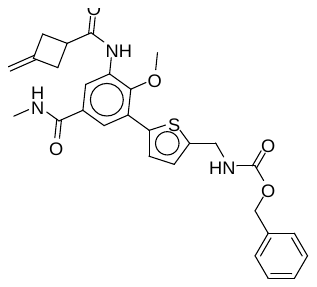

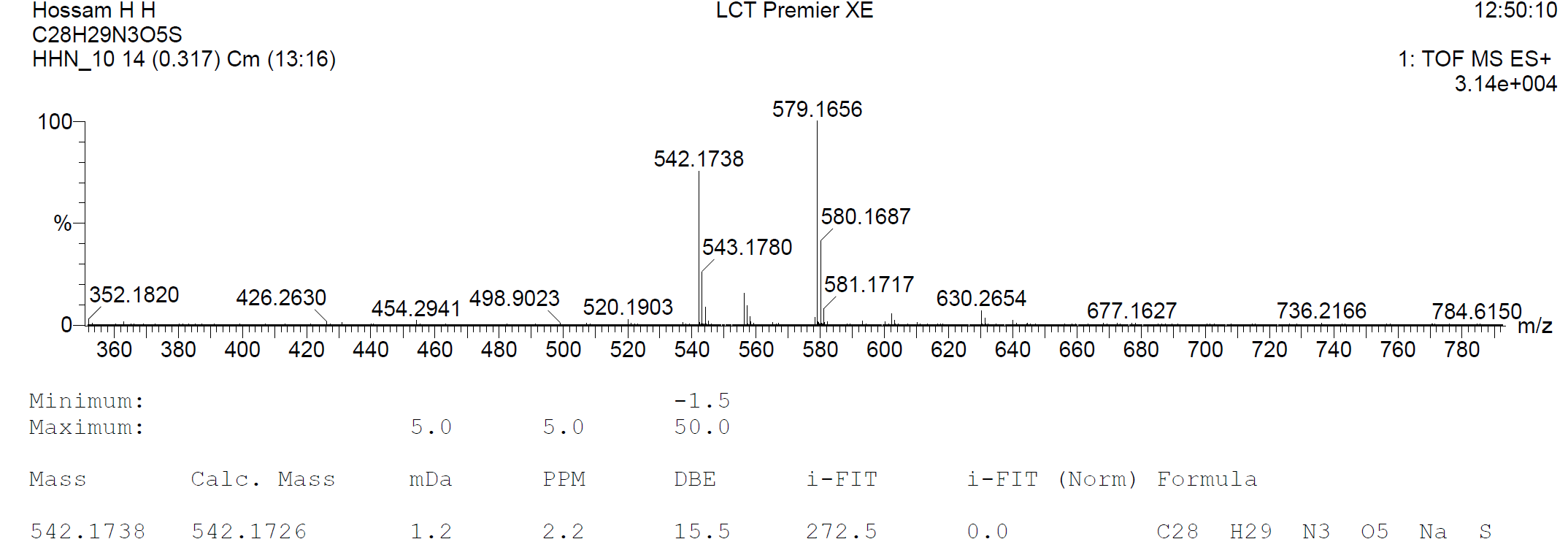

**Figure S21.** HRMS spectrum of compound **4e**.

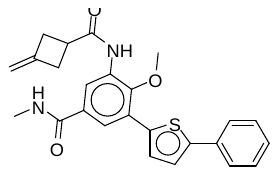

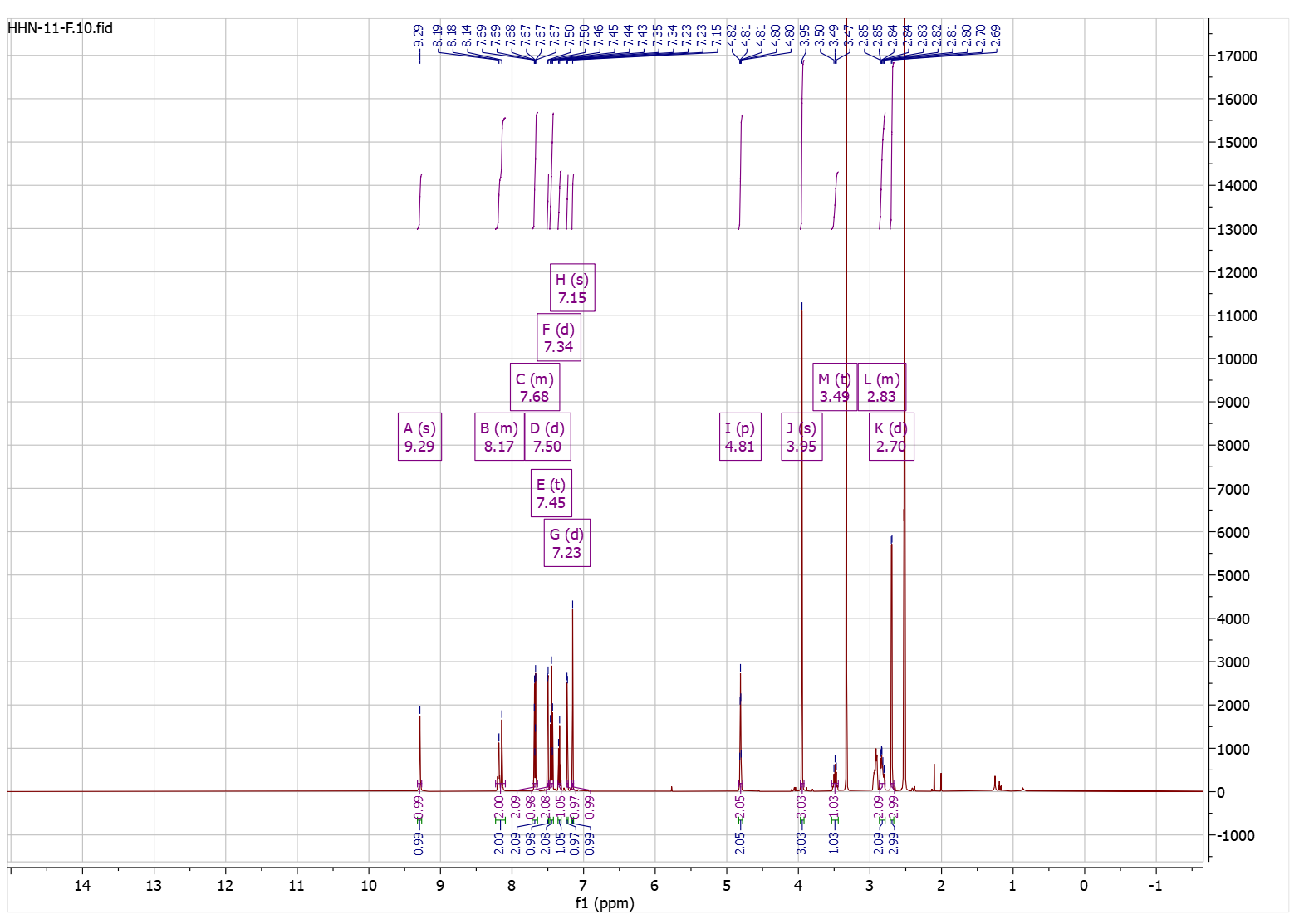

**Figure S22.** ^1^H-NMR spectrum of compound **4f**.

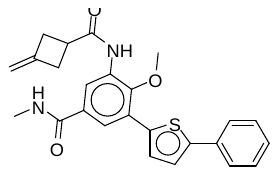

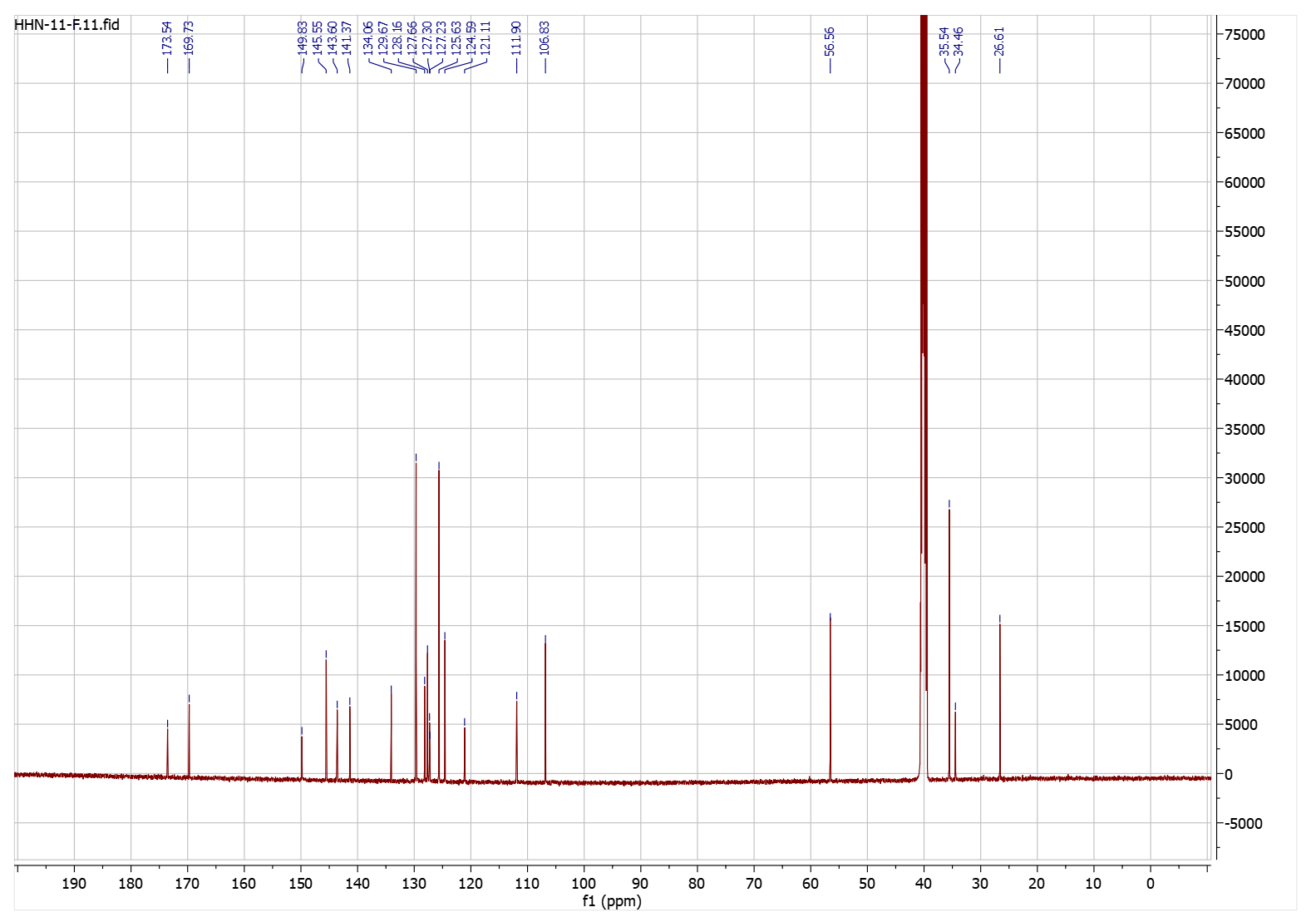

**Figure S23.** ^13^C-NMR spectrum of compound **4f**.

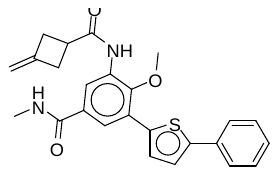

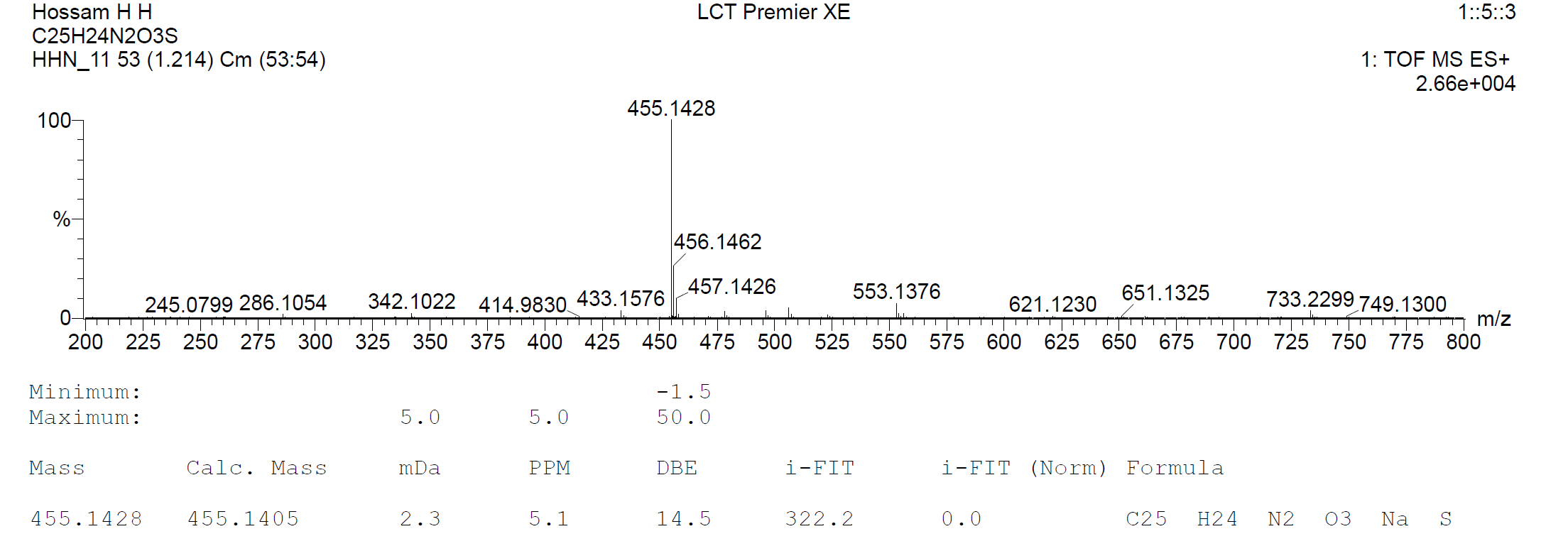

**Figure S24.** HRMS spectrum of compound **4f**.

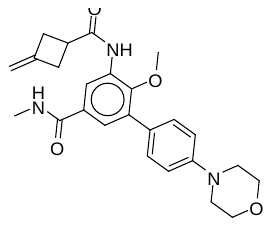

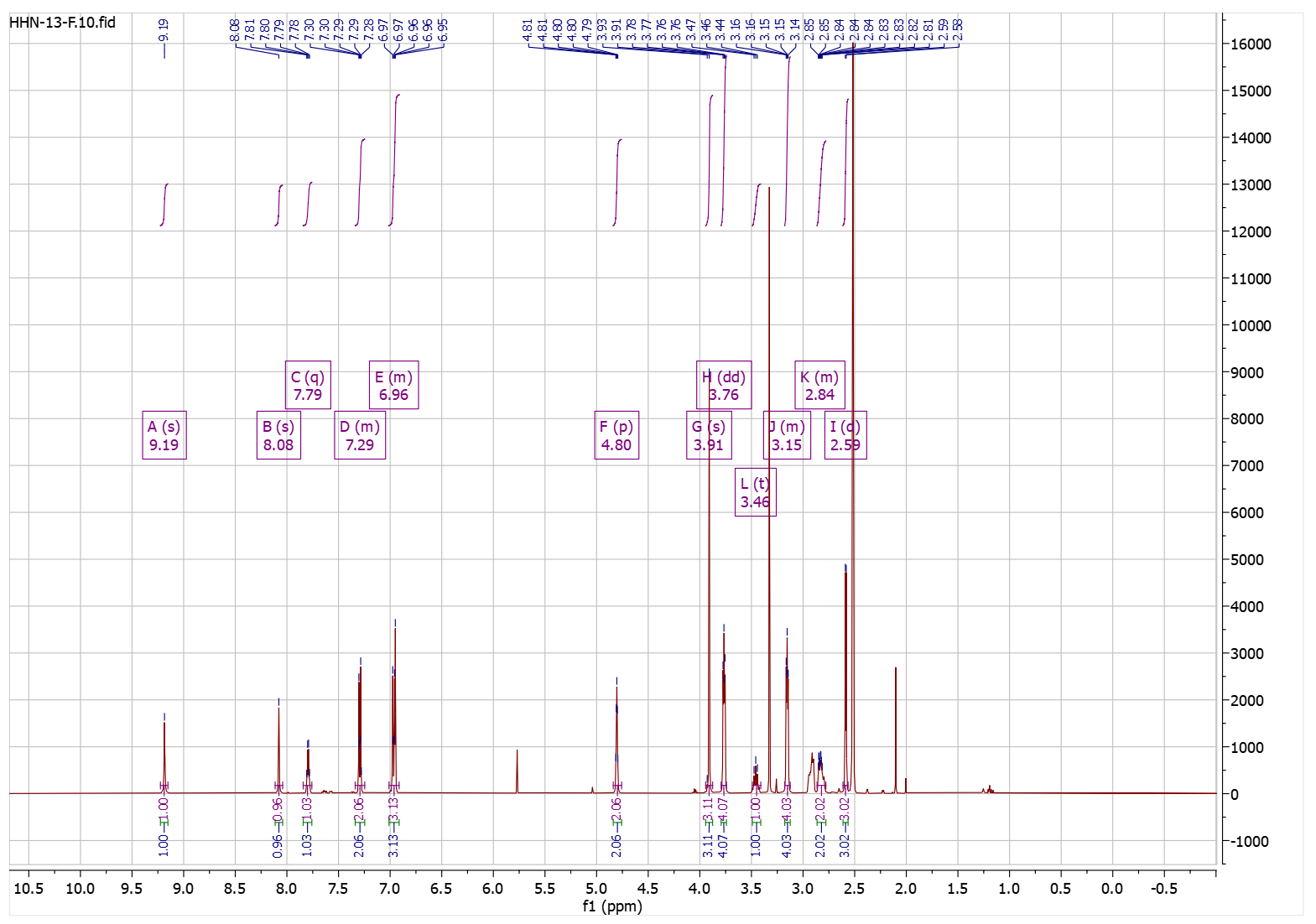

**Figure S25.** ^1^H-NMR spectrum of compound **4g**.

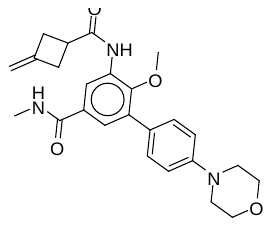

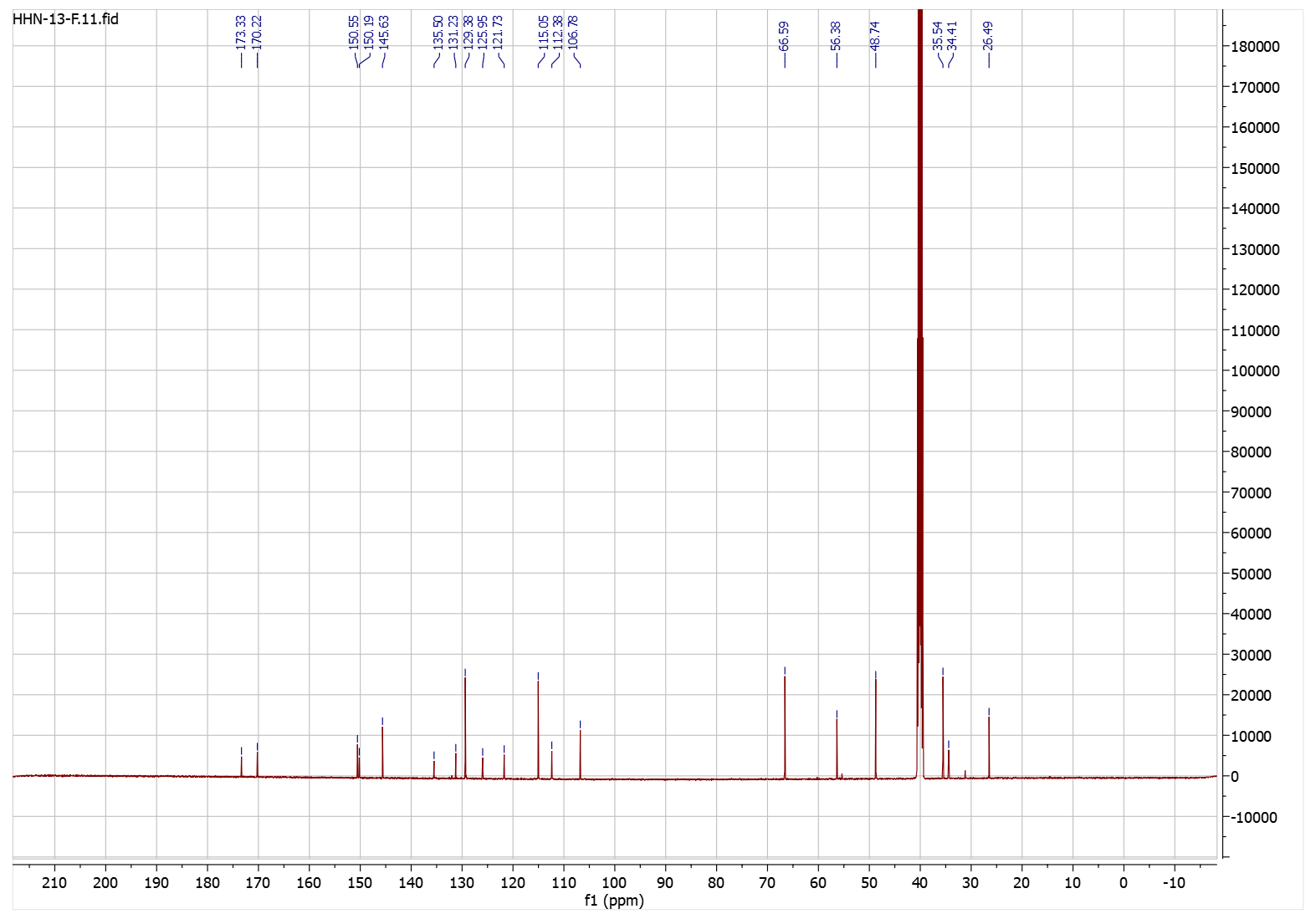

**Figure S26.** ^13^C-NMR spectrum of compound **4g**.

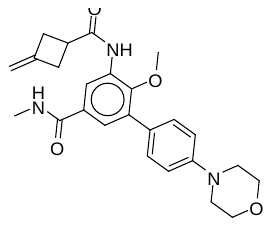

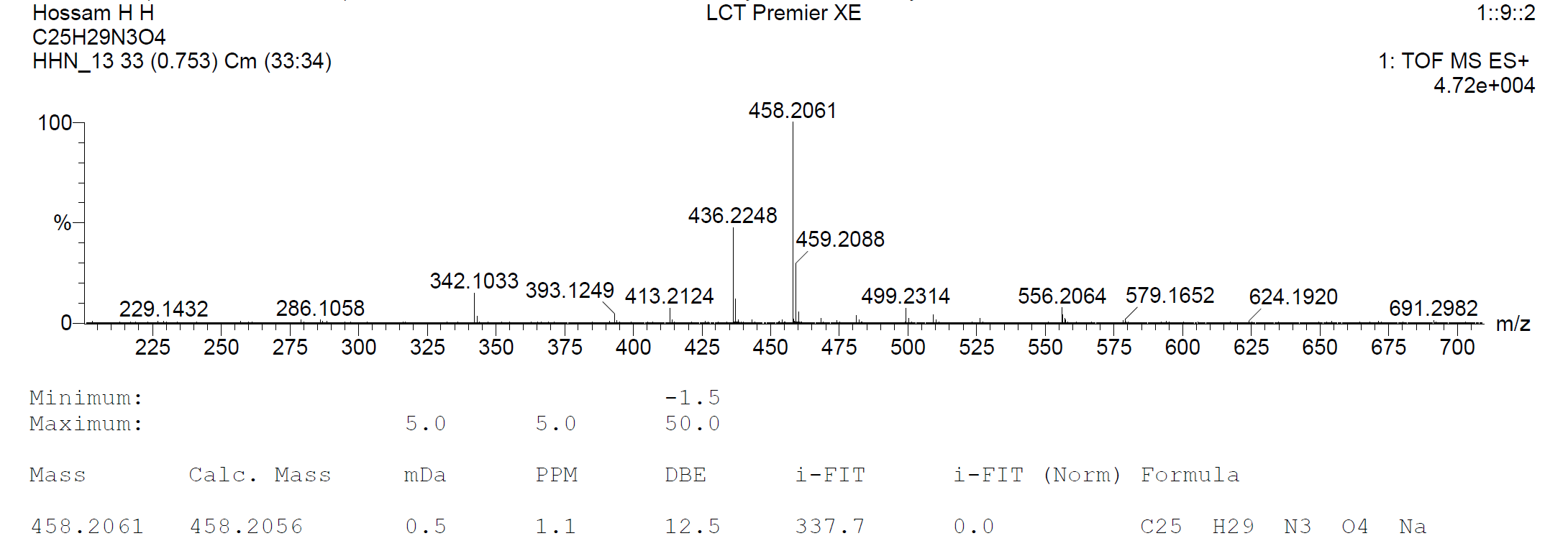

**Figure S27.** HRMS spectrum of compound **4g**.

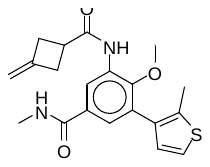

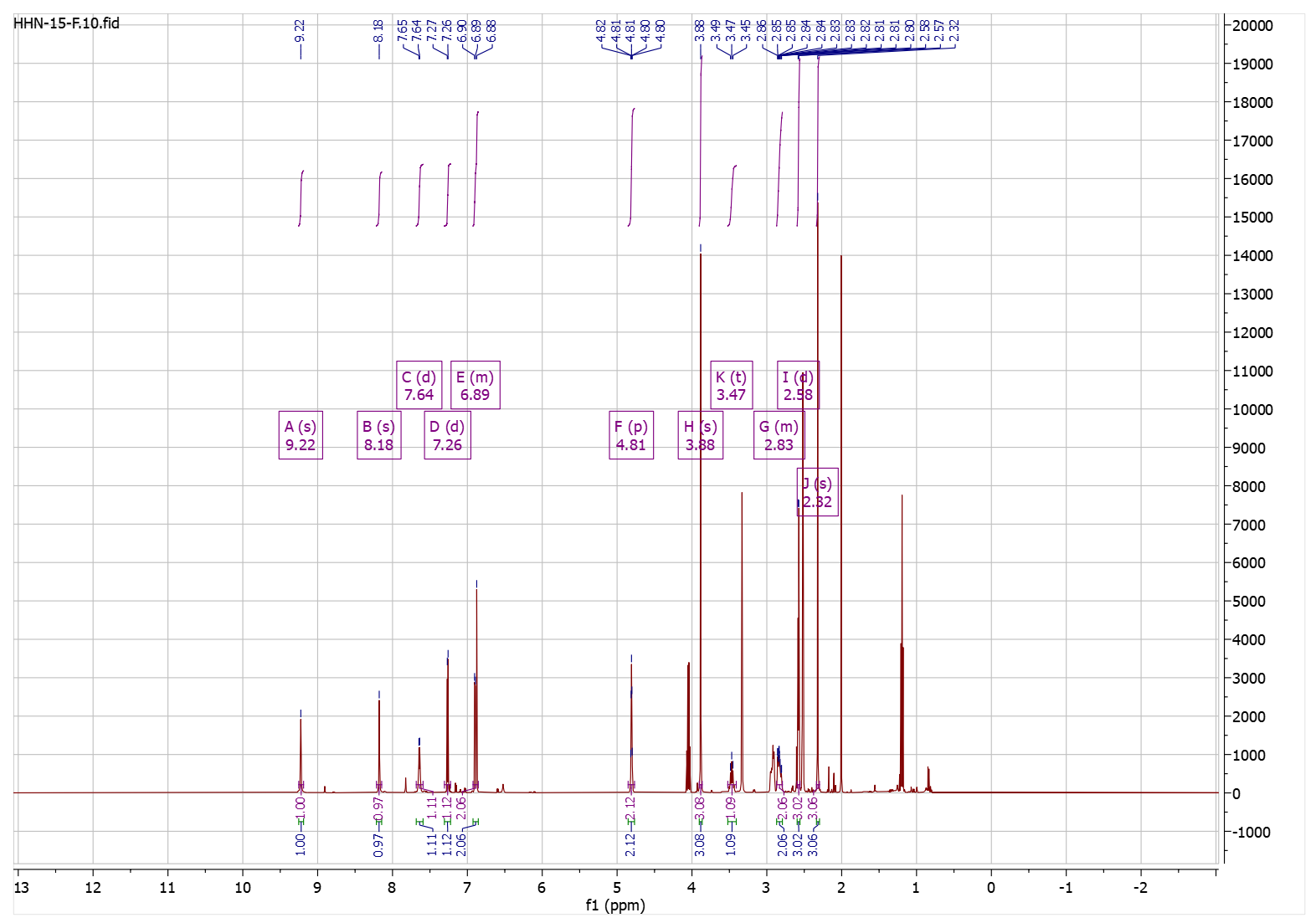

**Figure S28.** ^1^H-NMR spectrum of compound **4h**.

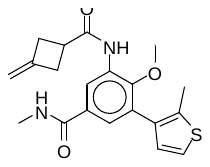

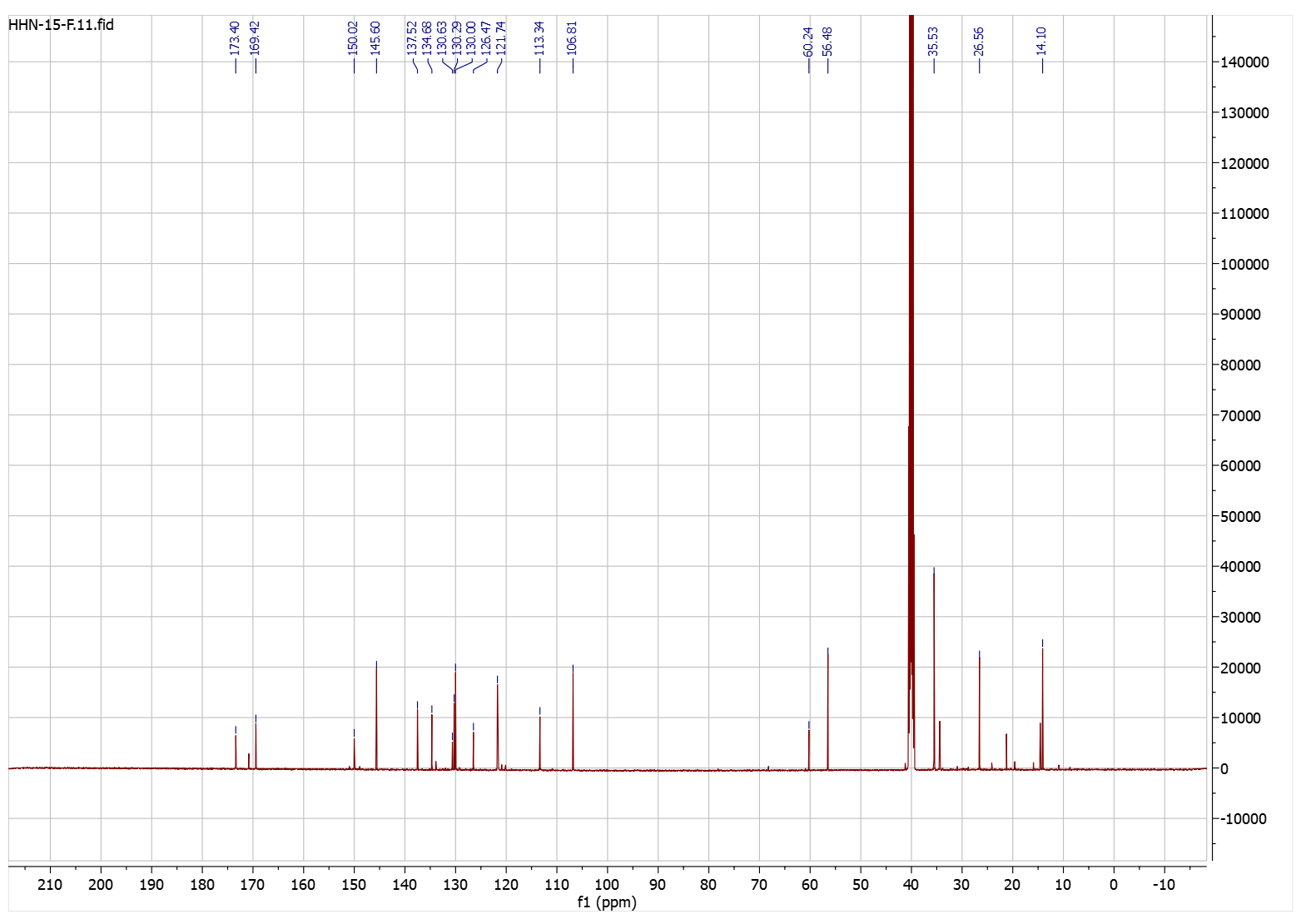

**Figure S29.** ^13^C-NMR spectrum of compound **4h**.

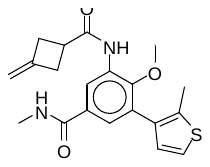

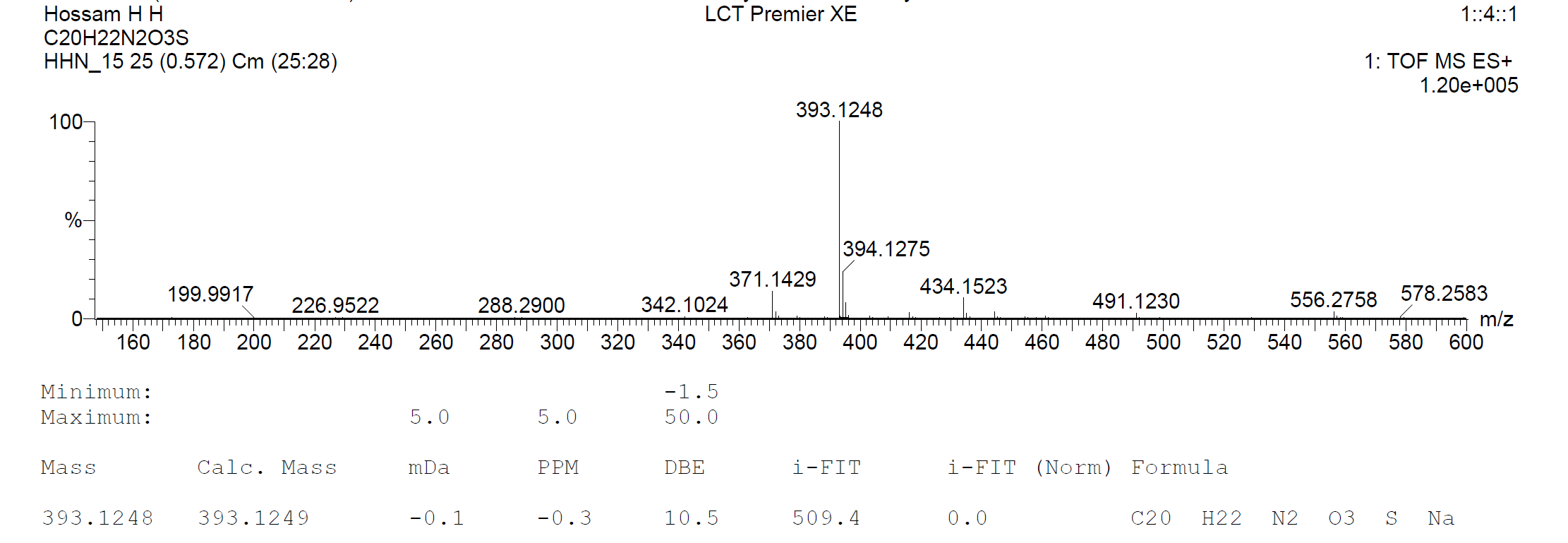

**Figure S30.** HRMS spectrum of compound **4h**.

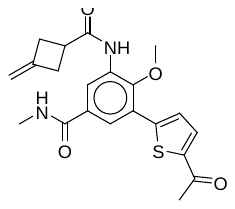

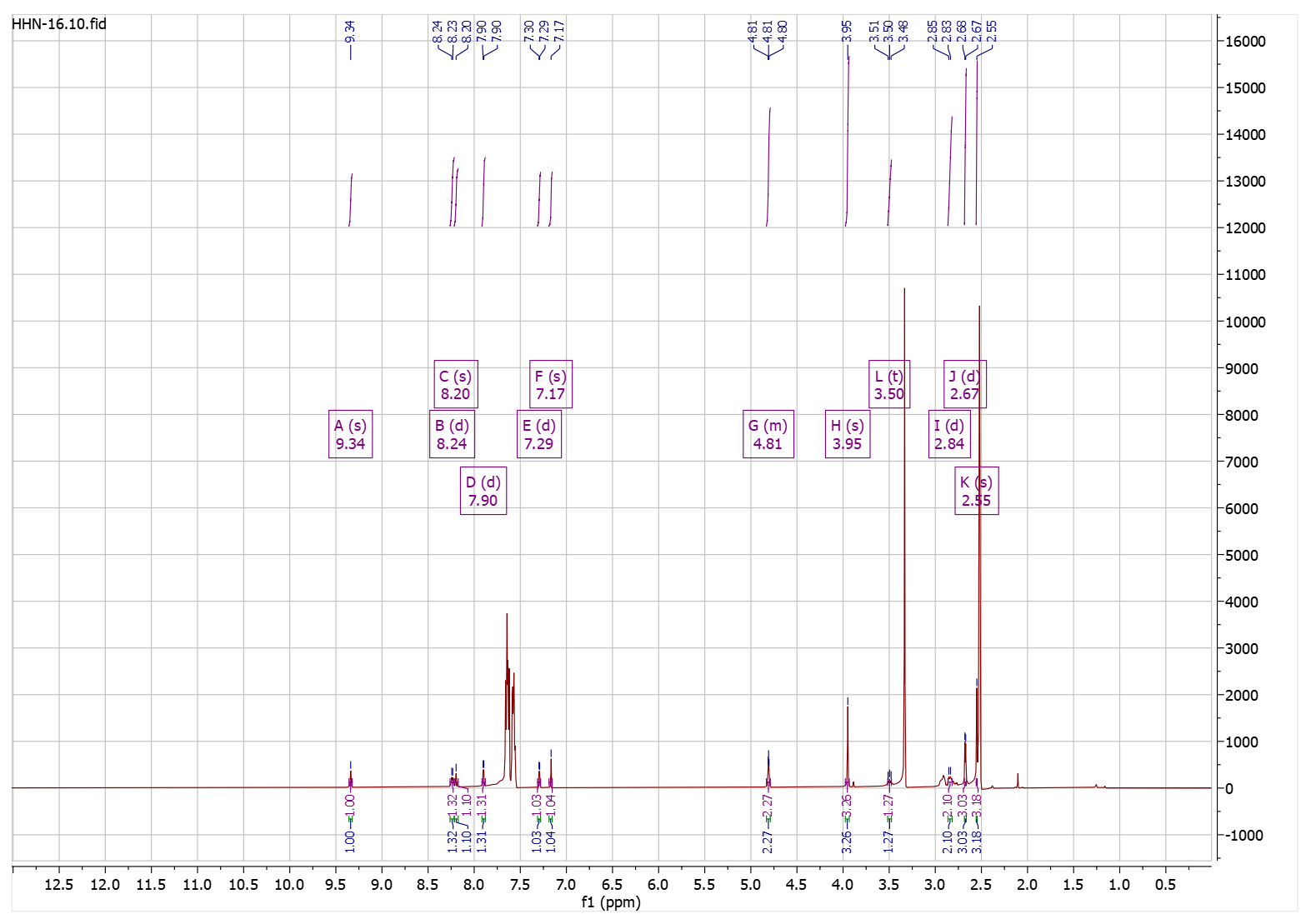

**Figure S31.** ^1^H-NMR spectrum of compound **4i**.

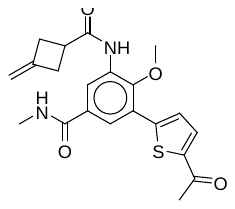

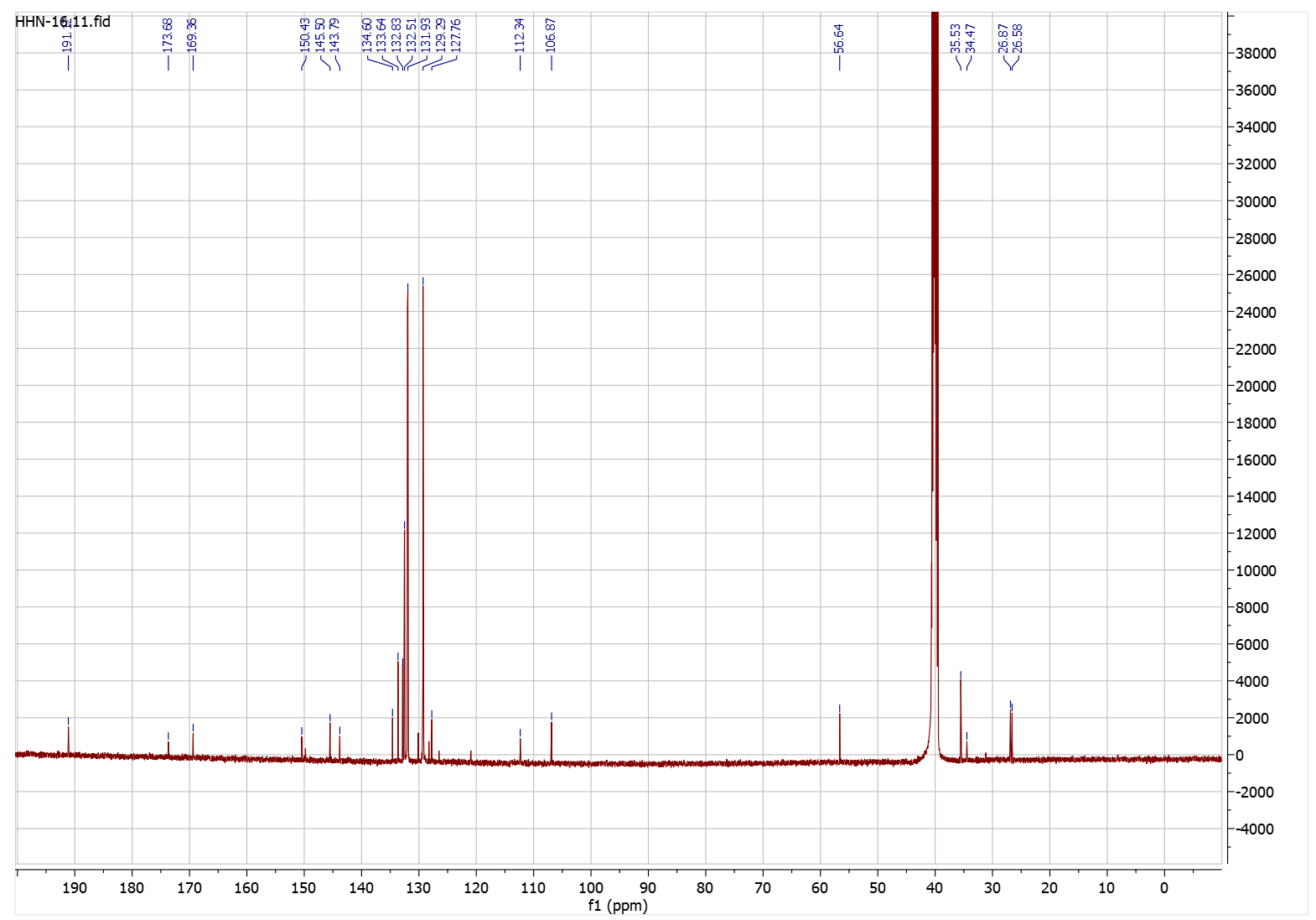

**Figure S32.** ^13^C-NMR spectrum of compound **4i**.

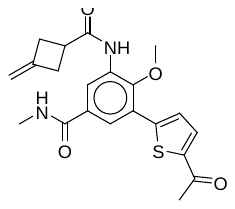

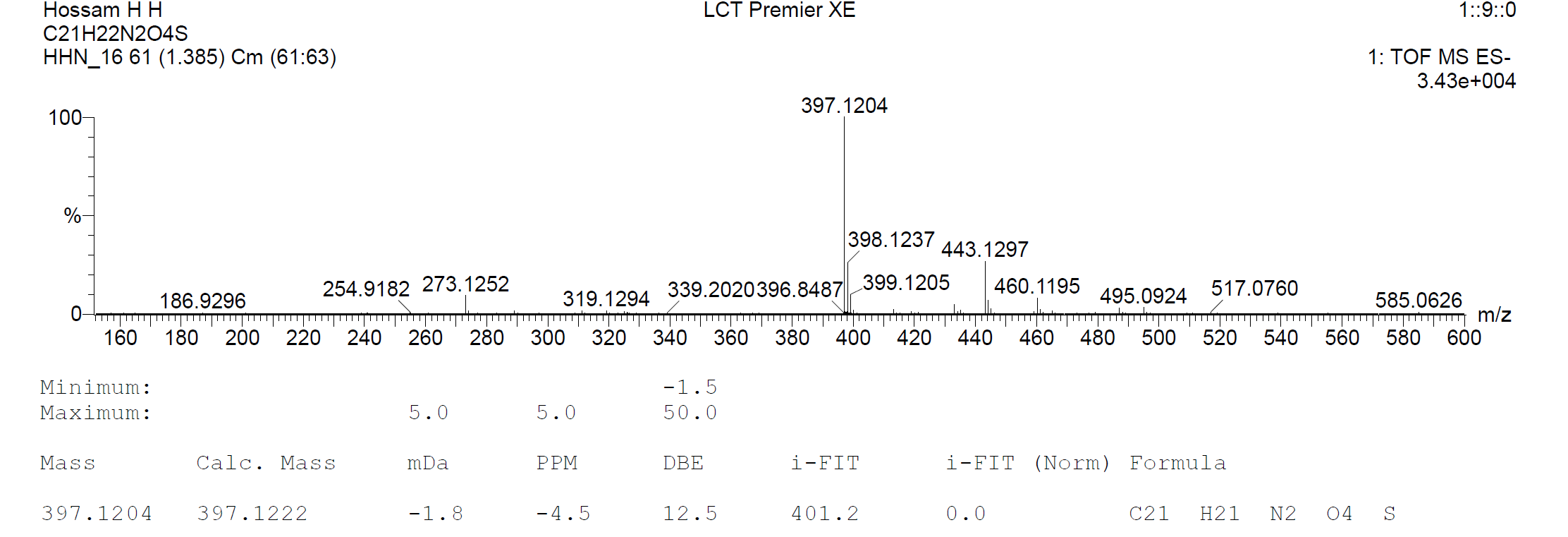

**Figure S33.** HRMS spectrum of compound **4i**.

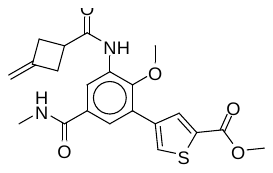

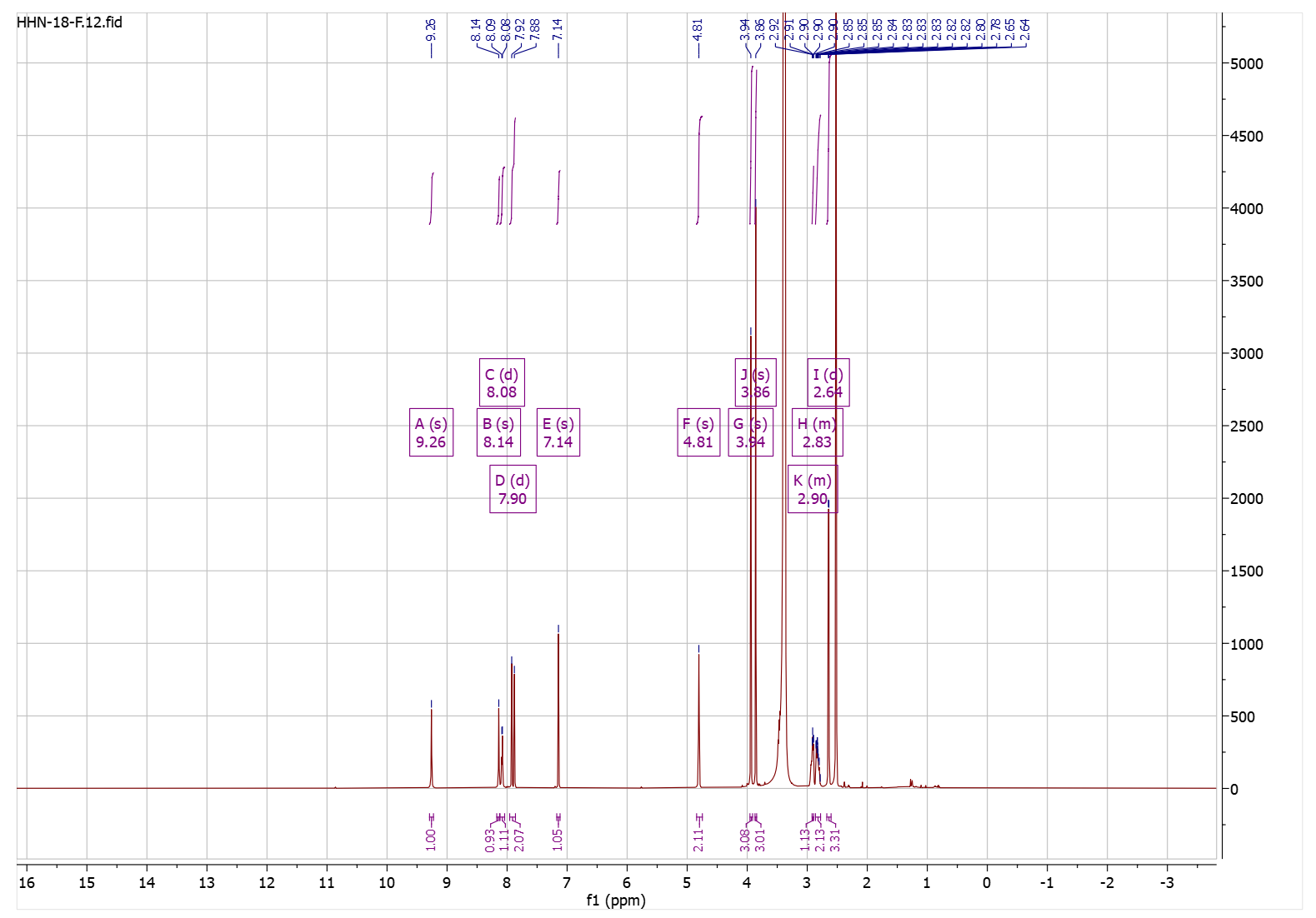

**Figure S34.** ^1^H-NMR spectrum of compound **4j**.

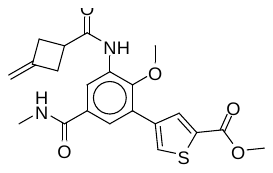

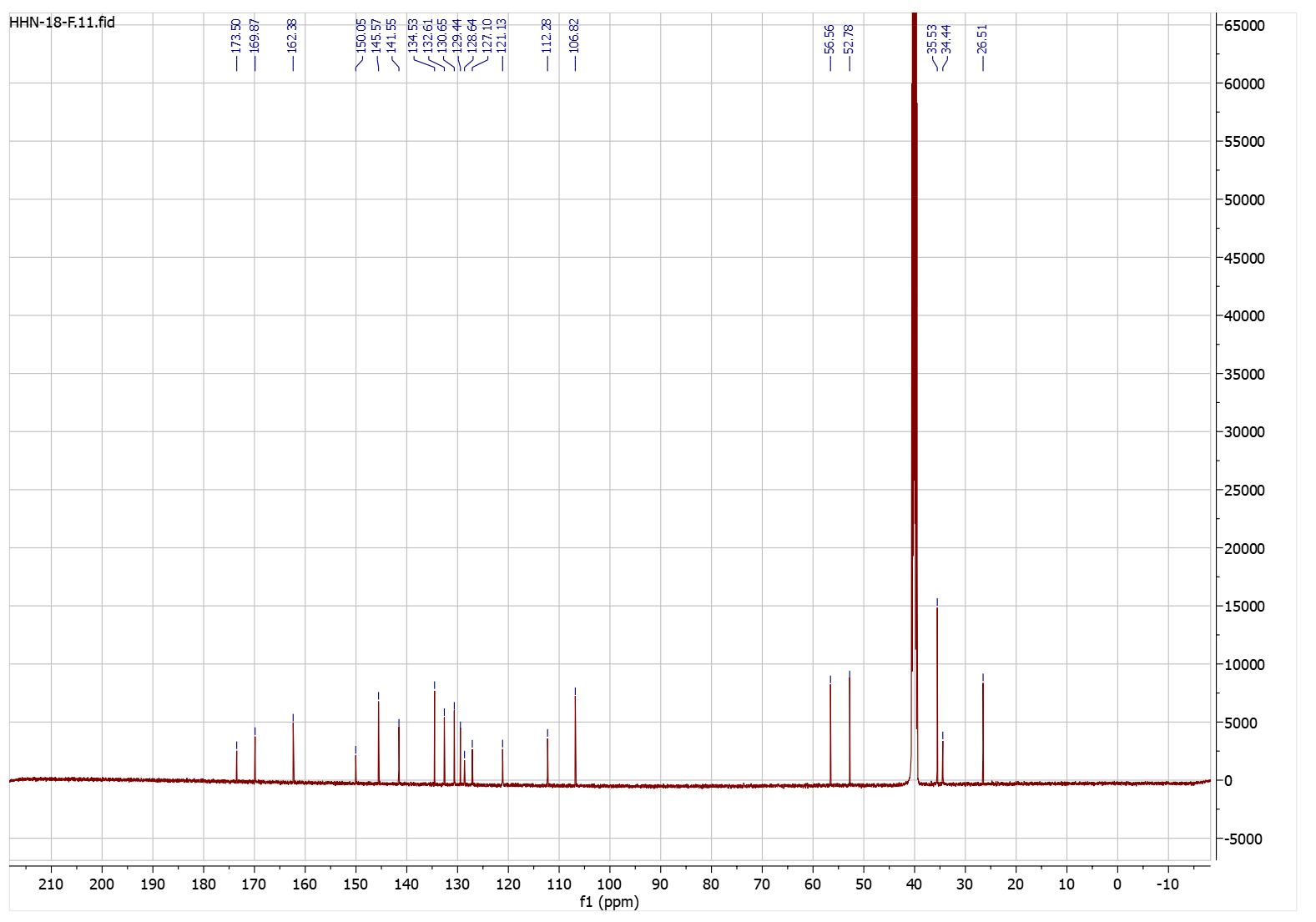

**Figure S35.** ^13^C-NMR spectrum of compound **4j**.

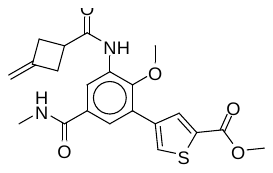

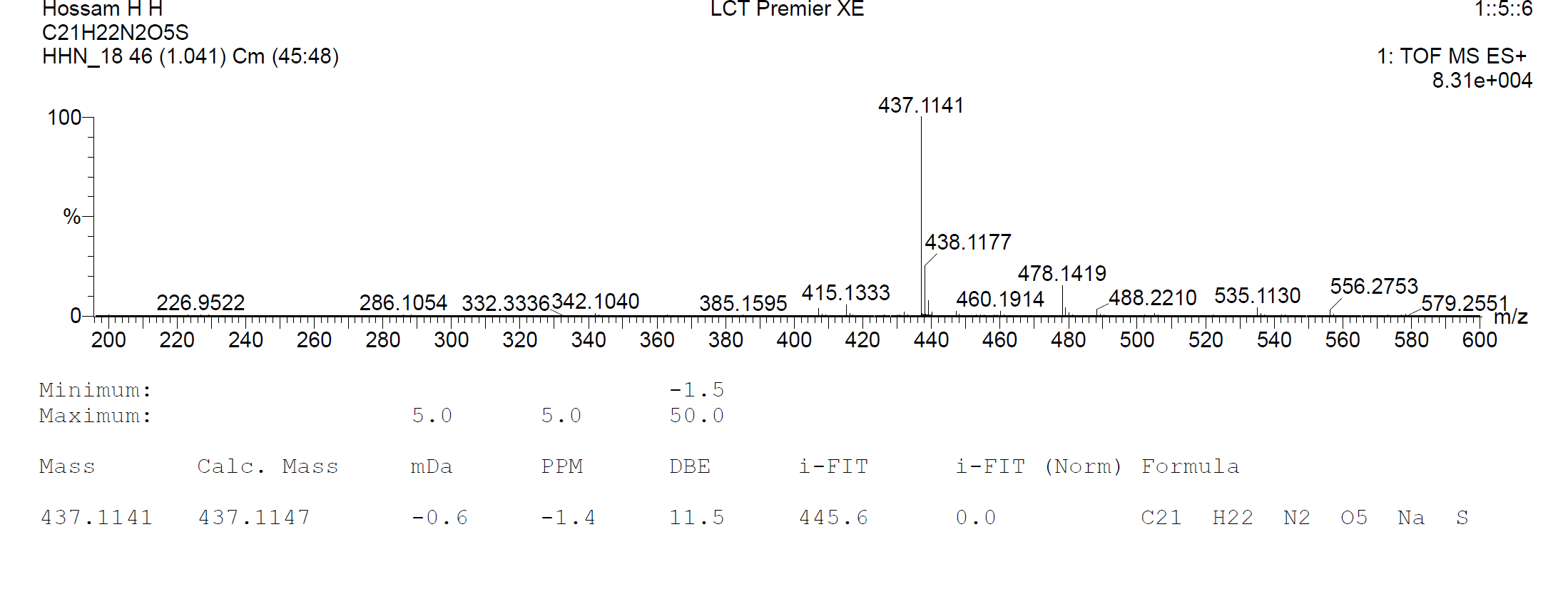

**Figure S36.** HRMS spectrum of compound **4j**.

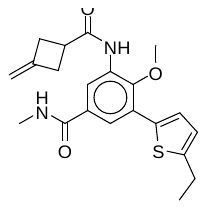

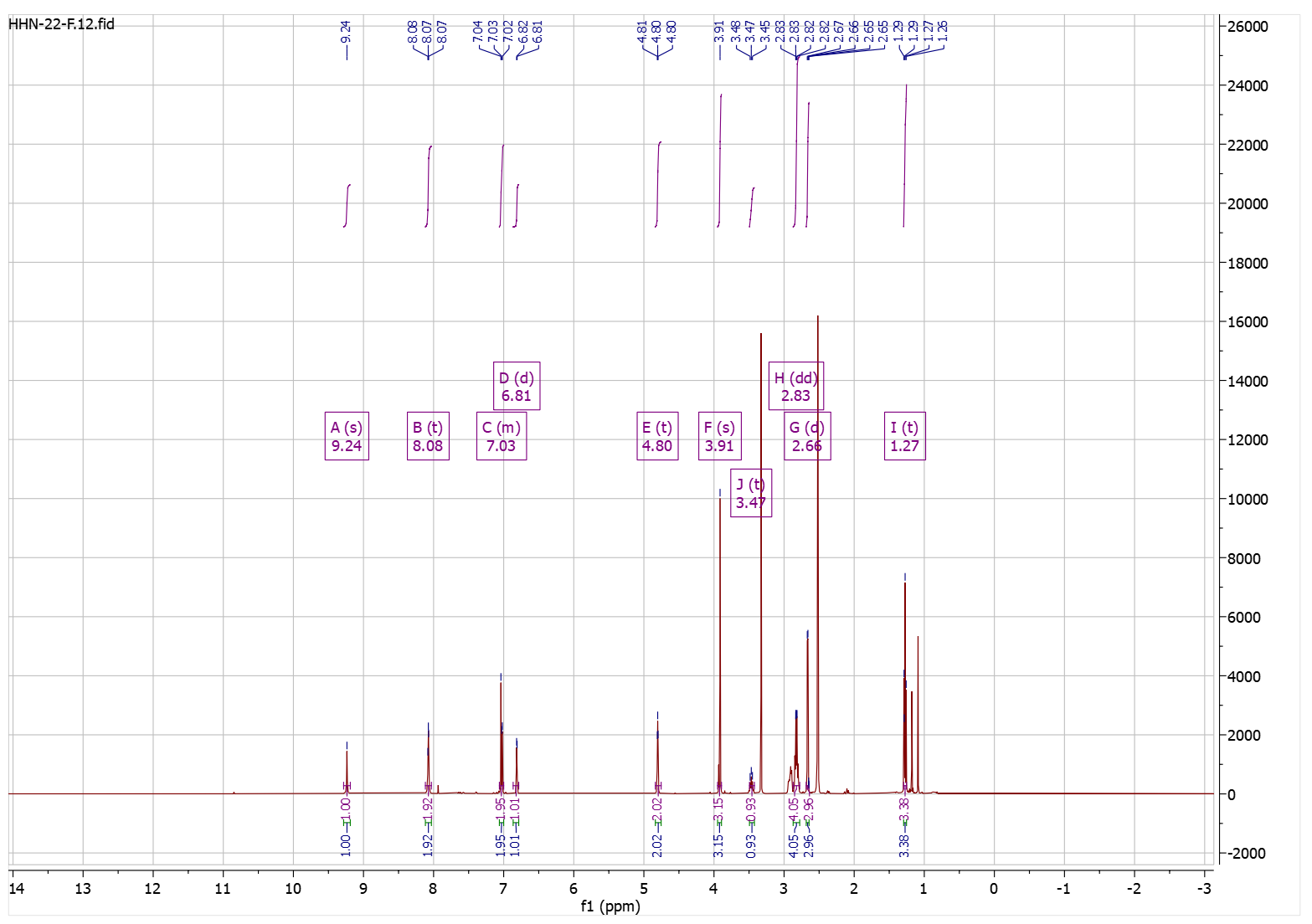

**Figure S37.** ^1^H-NMR spectrum of compound **4k**.

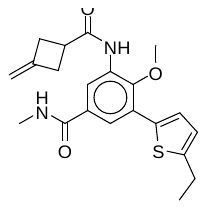

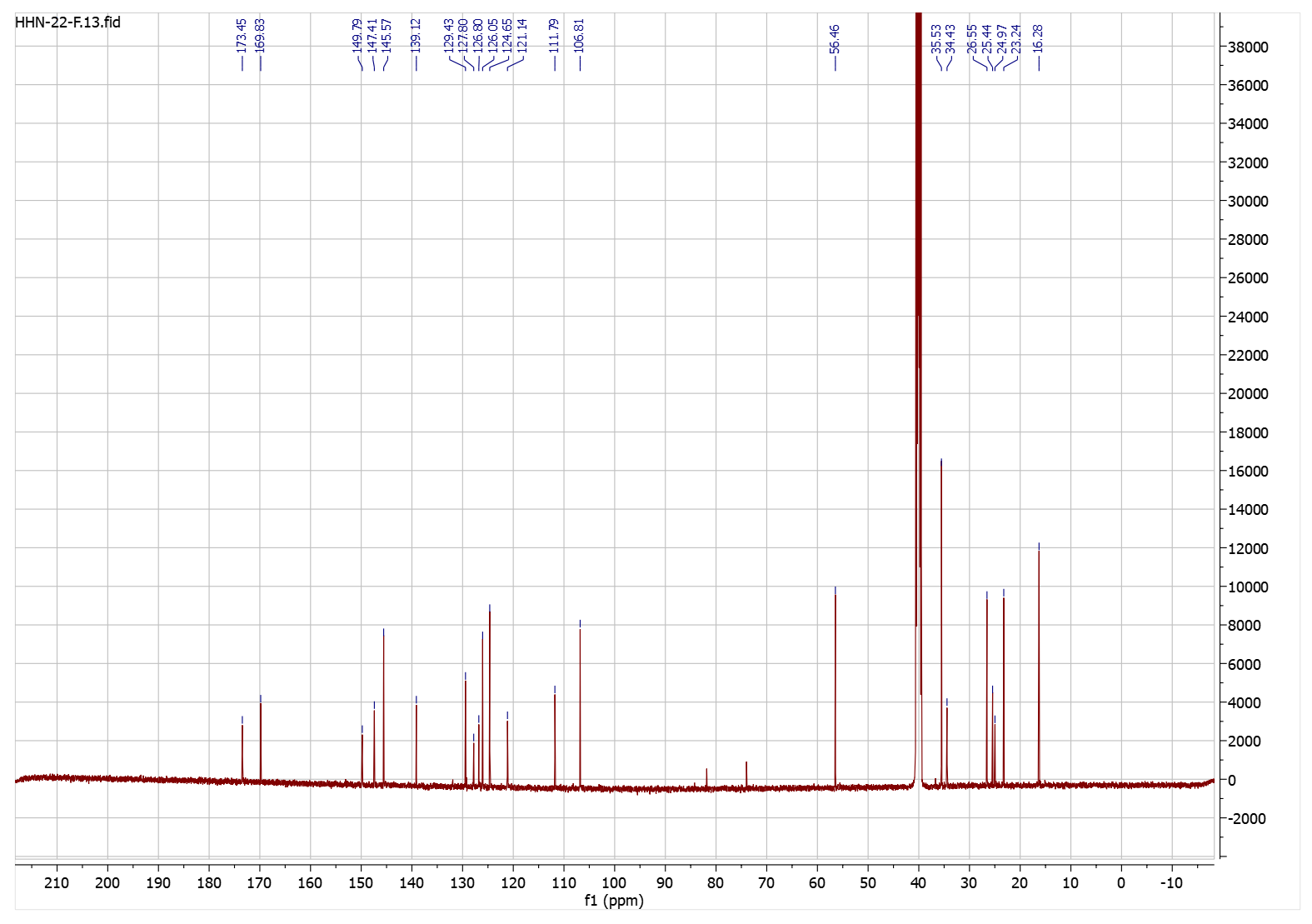

**Figure S38.** ^13^C-NMR spectrum of compound **4k**.

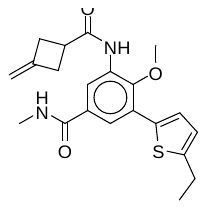

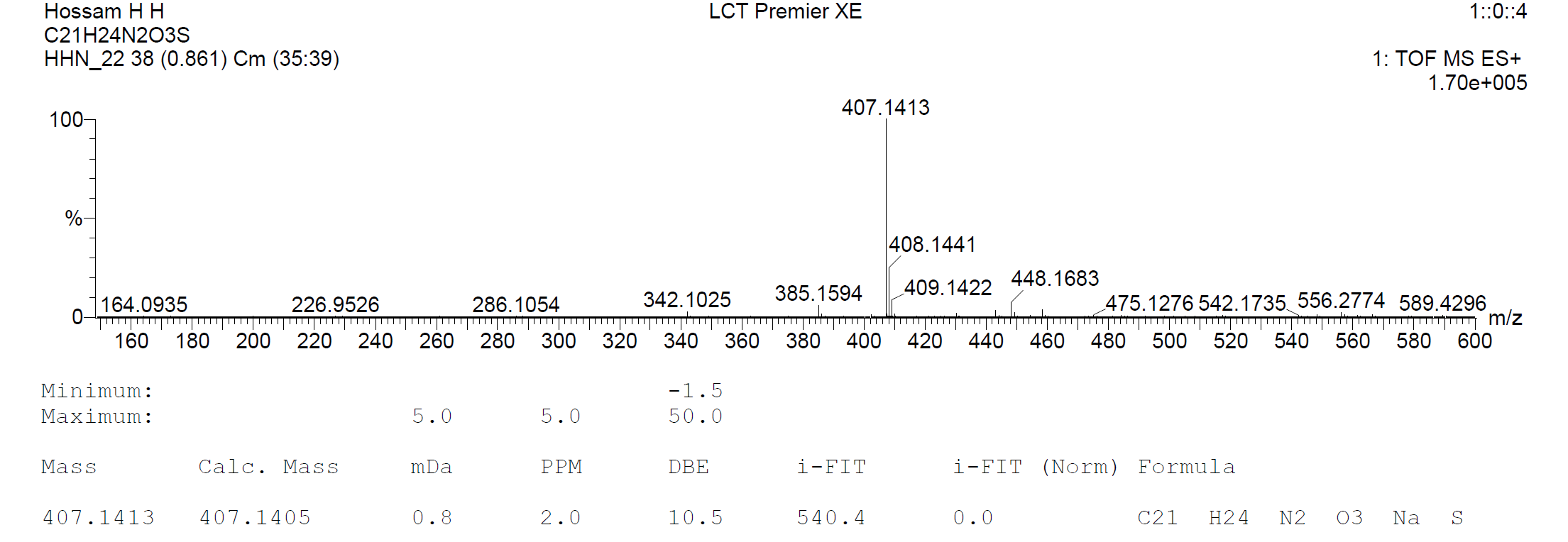

**Figure S39.** HRMS spectrum of compound **4k**.

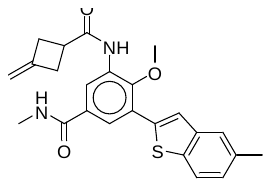

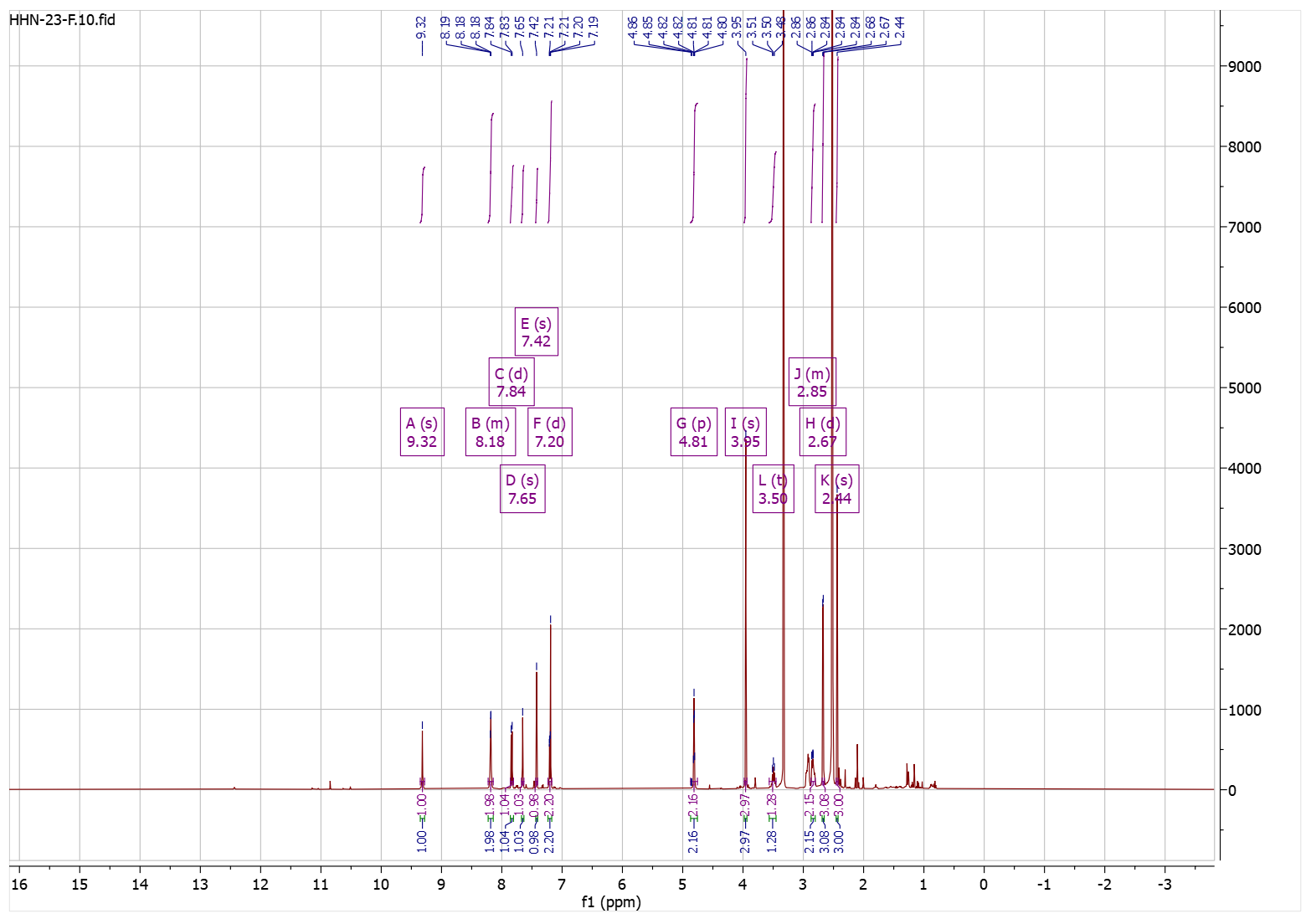

**Figure S40.** ^1^H-NMR spectrum of compound **4l**.

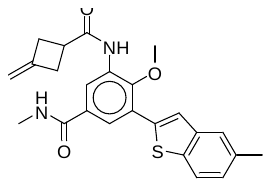

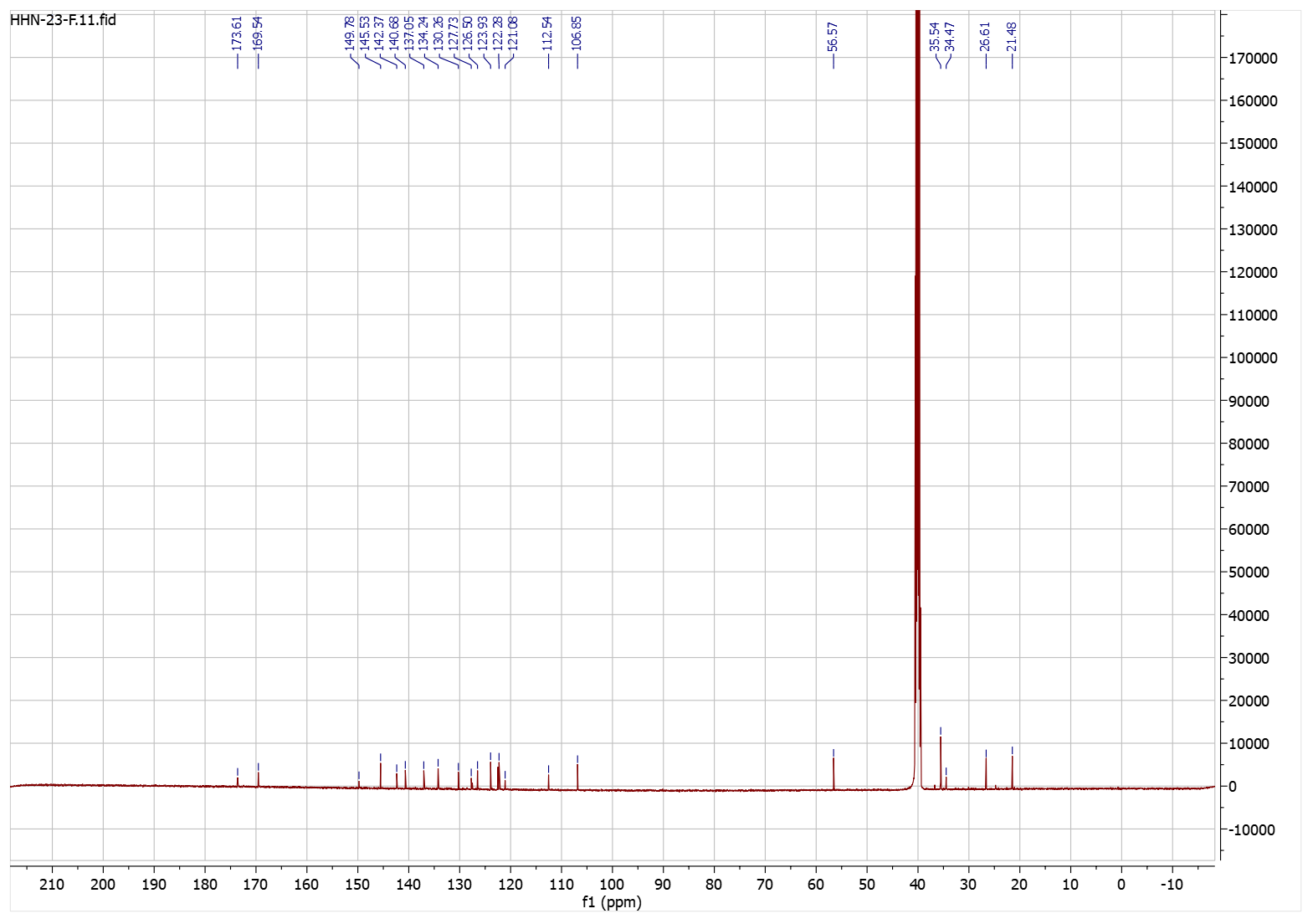

**Figure S41.** ^13^C-NMR spectrum of compound **4l**.

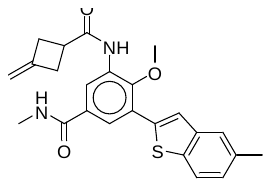

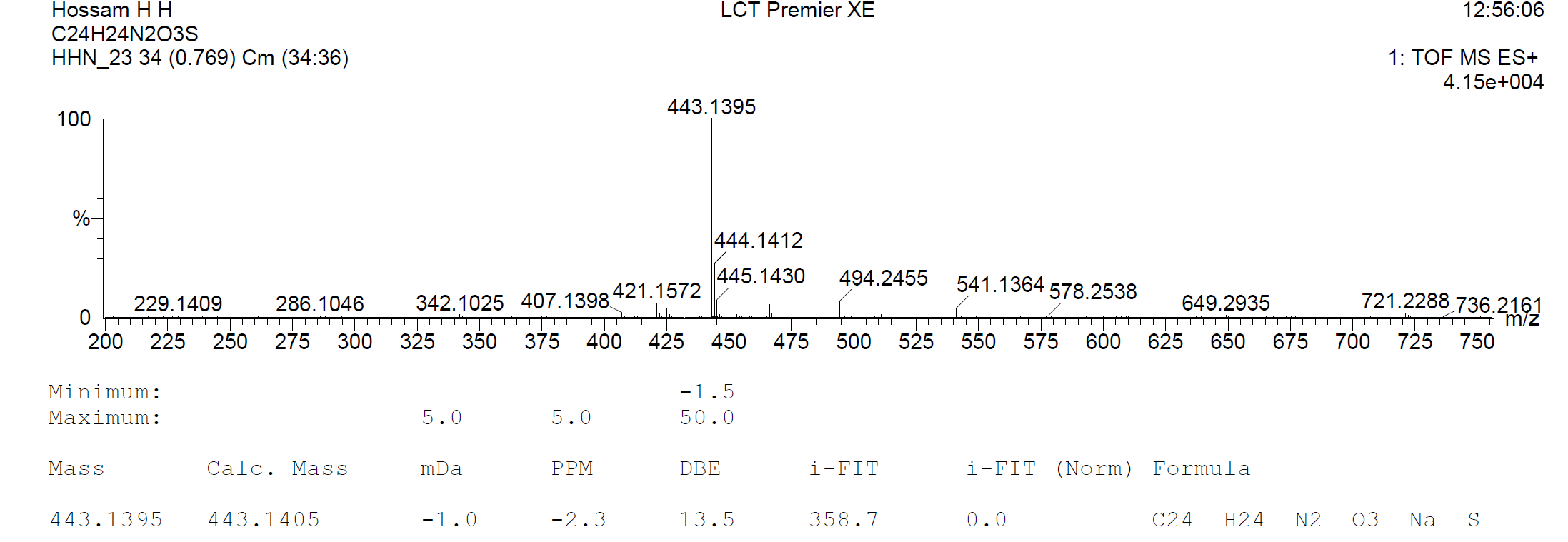

**Figure S42.** HRMS spectrum of compound **4l**.
